# Assessing Generative Model Coverage of Protein Structures with SHAPES

**DOI:** 10.1101/2025.01.09.632260

**Authors:** Tianyu Lu, Melissa Liu, Yilin Chen, Jinho Kim, Po-Ssu Huang

**Affiliations:** Department of Bioengineering, Stanford University, Stanford, CA, USA; Department of Physics, Stanford University, Stanford, CA, USA

## Abstract

Recent advances in generative modeling enable efficient sampling of protein structures, but their tendency to optimize for designability imposes a bias toward idealized structures at the expense of loops and other complex structural motifs critical for function. We introduce SHAPES (Structural and Hierarchical Assessment of Proteins with Embedding Similarity) to evaluate five state-of-the-art generative models of protein structures. Using structural embeddings across multiple structural hierarchies, ranging from local geometries to global protein architectures, we reveal substantial undersampling of the observed protein structure space by these models. We use Fréchet Protein Distance (FPD) to quantify distributional coverage. Different models are distinct in their coverage behavior across different sampling noise scales and temperatures; the frequency of TERtiary Motifs (TERMs) further supports the observations. More robust sequence design and structure prediction methods are likely crucial in guiding the development of models with improved coverage of the designable protein space.

## INTRODUCTION

Navigating the design space of protein structures has traditionally relied on domain expertise guided by energy functions^1^. With the introduction of generative models, structure-based design and hypothesis generation has become more robust. Instead of relying on energy functions, the protein structure space parameterized under a generative model facilitates sample generation and selection according to designability metrics^2^. While generative models are capable of efficiently sampling from protein structure space, evaluation of their capacity to generate all observed structural features in proteins remains understudied and unquantified.

Generative models aim to capture a complex data distribution by applying a learned transformation which turns noise into realistic samples (Figure 1A)^3^. An application of generative modeling is the ability to sample novel, diverse and realistic protein structures. The quality of a sampled backbone is typically evaluated with designability, measured *in silico* by computing the Root Mean Square Deviation (RMSD) of the predicted structures from the sequences designed for it with the original backbone. In our analysis, we designed eight sequences with ProteinMPNN^4^ and predicted their structures with ESMFold^5^. A protein backbone is said to be designable if at least one predicted structure has RMSD *<* 2.0 Å ngstroms to the designed backbone. Many experimental successes have been reported, making designability a robust metric to filter designs^6–8^. However, sampled structures are often idealized, containing higher proportions of alpha helices and beta sheets than native structures deposited in the Protein Data Bank (PDB) (Figure 1B)^9^. This biased sampling of secondary structures motivates the following questions: 1. What regions of protein structure space are not being covered by generative models optimized for designability? 2. How can we quantify and interpret the distributional coverage of protein structure space? 3. What limitations does biased sampling place on the ability to design functional structural elements?

**Figure 1.**
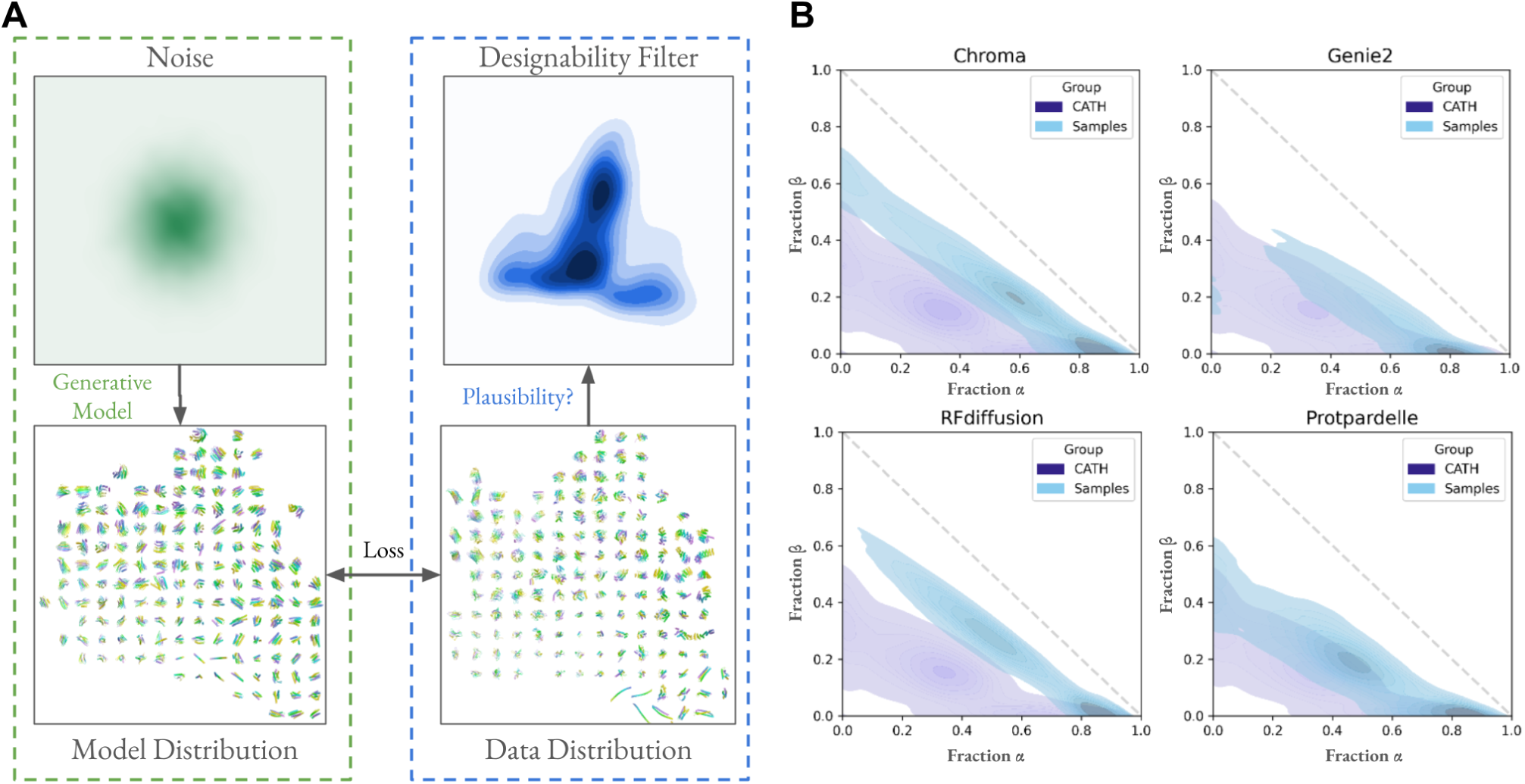
Generative models capture a biased set of protein structure space. (A) Generative models of protein structures convert noise to samples which are optimized to match the data distribution, e.g. CATH. Models are optimized for the ability to draw samples with high designability, which implicitly imposes a filter that over-emphasizes the designable subspace of protein structures and under-emphasizes the undesignable subspace. (B) Secondary structure elements more prominently show alpha helices and beta sheets in sampled structures compared to native structures in CATH.

To answer these questions, we present the SHAPES framework to analyze the behavior of generative models of protein structures. We quantify distributional similarity with Fréchet Protein Distance (FPD), analogous to the Fréchet ProtT5 Distance introduced by Alamdari *et al*.^10^ but using embeddings of protein structures instead of embeddings of protein sequences. The SHAPES evaluation framework consists of sampling a set of structures from a generative model, computing embeddings and the FPD of the embeddings with a reference dataset to quantify distribution similarity. We examine five different models: Chroma^11^, Genie2^12^, Protpardelle^13^, RFdiffusion^6^, and Multiflow^14^. They are all based on diffusion or flow-matching models, but each is trained with different structural representations and have different sampling schemes. RFdiffusion relies on a pretrained structure prediction model (RoseTTAFold) while other models are trained from scratch. Each residue is represented by a frame (rotation and translation), where both are denoised during sampling. Multiflow also uses residue frames but introduces a discrete flow-matching objective for sequence prediction trained jointly with the structure flow-matching objective. Chroma introduces correlated noise that models the polymer chain structure and the scaling of the radius of gyration of proteins. Genie2 uses Frenet-Serret frames during denoising formed by triplets of adjacent alpha-carbon atoms. Protpardelle treats each residue as atomic coordinates instead of frames and is not equivariant to rotations, unlike the network architectures in other models. These differences result in different parameterizations of the distribution of protein structures.

We show that many models do not cover the full diversity of structural elements at all structural hierarchies, from nearest neighbor geometries, to local amino acid environment shells, to global protein architectures. While coverage improves with higher temperatures and noise scales during sampling, such samples also reveal unique pathologies of generated structures which make them less designable than low-temperature samples. Finally, we use TERtiary Motifs (TERMs)^15^ to validate the FPD trends and show that complex functional motifs involving loops are more likely to be found in samples drawn from models with greater coverage of the PDB.

## RESULTS

### Optimizing Designability Leads to Complexity Reduction

Designability aims to answer the question of whether or not there exists a sequence which can fold into a given backbone. By definition, this criterion is satisfied for all accurately modeled structures deposited in the PDB. However, we find that 43.7% of structures in CATH^16^ are not designable, even when using the *native* sequence, in agreement with recent works which show similar results^14,17^. Using designability as a metric to guide the development of generative models and for ranking designs inevitably steers sampling towards the *designable* subset of protein structures and does not measure the ability to model the full set of observed protein structures.

For generative models optimized to maximize designability, what structural features do they introduce? To understand this, we partially add noise to 21,663 structures generated with Protpardelle at high temperature, which are on average less designable than low temperature samples, then denoise the structures with RFdiffusion which produces more designable structures on average. Indeed, designability improves across all lengths (Figure 2B), but is coupled with a reduction in loop content and an increase in alpha and beta secondary structures (Figure 2A). Notably, the RFdiffusion-induced vector field which transports secondary structure density from the Protpardelle to the Partial Diffusion distributions consistently shows movement towards higher secondary structure content, except near the diagonal boundary where beta content is replaced with alpha content. We show an example in which the front-facing loop is remodeled into two helices with a minimal turn (Figure 2C). We posit that the gain in designability is partly attributed to complexity reduction, in particular the erasure of loops which are difficult to design and predict. This behavior is valuable in design tasks which require engineering structural rigidity but could be a limitation in design tasks which require structural flexibility, such as engineering for allostery^18^.

**Figure 2.**
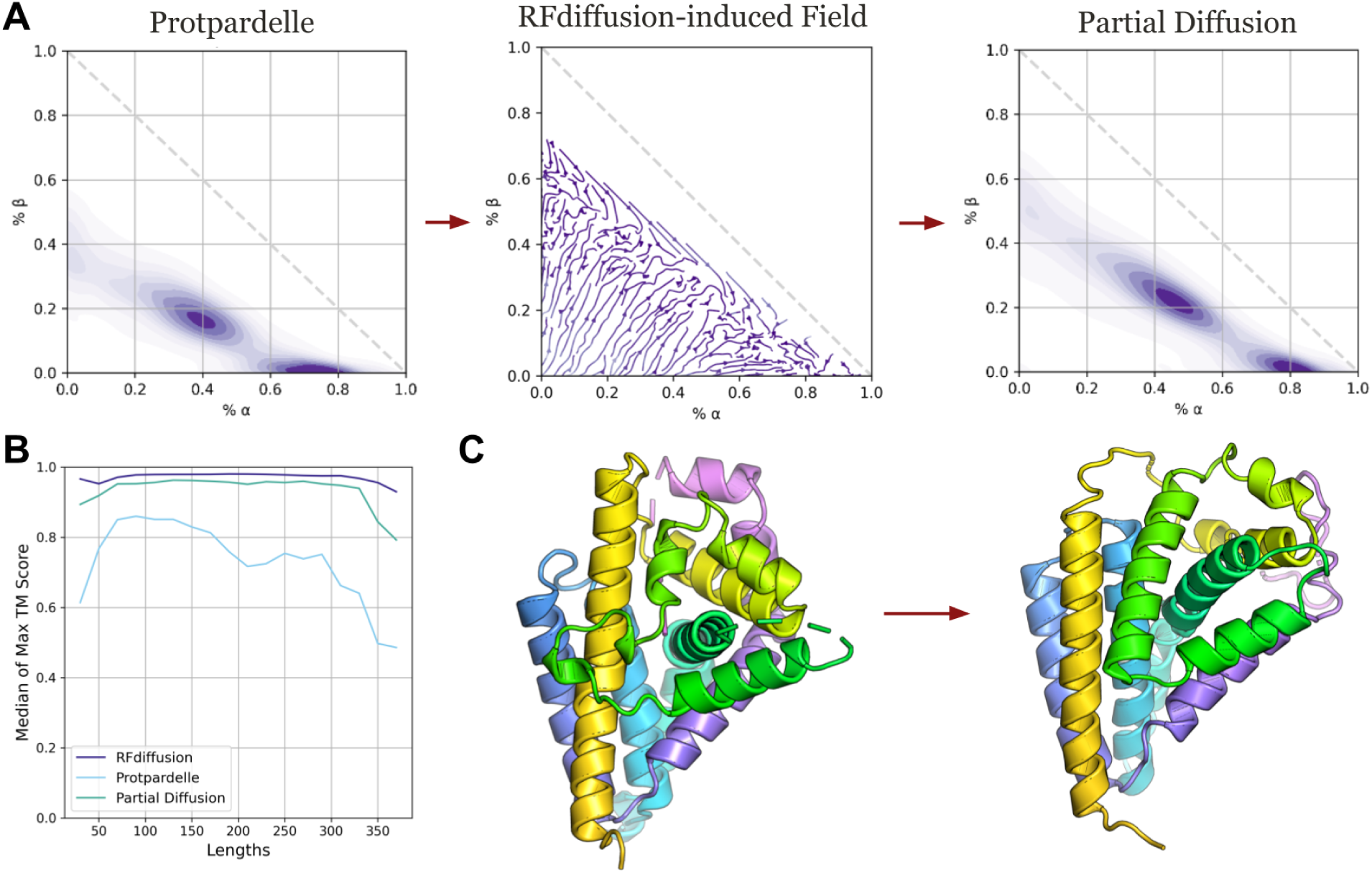
Partial diffusion with RFdiffusion reduces structural complexity. (A) 20 steps of partial diffusion using RFdiffusion applied to 21,663 samples of Protpardelle leads to reduced loop content and more alpha and beta content. Partial diffusion by RFdiffusion induces a vector field in secondary structure content. We approximate the true vector field using the start and end secondary structure content before and after partial diffusion. (B) The max TM score is taken over each group of eight ProteinMPNN designed sequences for each structure. The median is taken over all structures for every length. (C) Example of a high temperature sample from Protpardelle (left) and the structural edits made by 20 steps of RFdiffusion partial diffusion at the default sampling temperature.

### SHAPES Reveal Undersampled Regions of Protein Structure Space

The increase in secondary structure content does not fully elucidate the differences between designable and undesignable regions of protein structure space, as both designable and undesignable CATH structures have similar secondary structure distributions (Supplementary Figure 1). To capture more fine-grained features of protein geometry, we use learned representations of protein structures at all structural hierarchies. Local amino acid nearest neighbor geometries are represented by Foldseek tokens, local amino acid environments including second shell contacts and beyond are represented by ProteinMPNN and ESM3 embeddings, and the geometry of protein architectures are represented by ProtDomainSegmentor embeddings (Figure 3A). Foldseek tokens are learned through an autoencoding objective of nearest neighbor geometric features^19^. They represent a discrete structural alphabet used for accurate and rapid retrieval in large structural datasets. ProteinMPNN encoder embeddings are used by its decoder to predict amino acid identity, thus the embedding is rich in information on the structural context which surrounds all residue types^4^. ESM3 encoder embeddings are used by its decoder to predict masked coordinates rather than masked residue type, but again it necessitates the representation of diverse structural context^20^. ProtDomainSegmentor embeddings are used to predict the CATH architecture for each residue^21^. Indeed, principal components of such embeddings exhibit more separation between designable and undesignable structures (Supplementary Figure 1).

**Figure 3.**
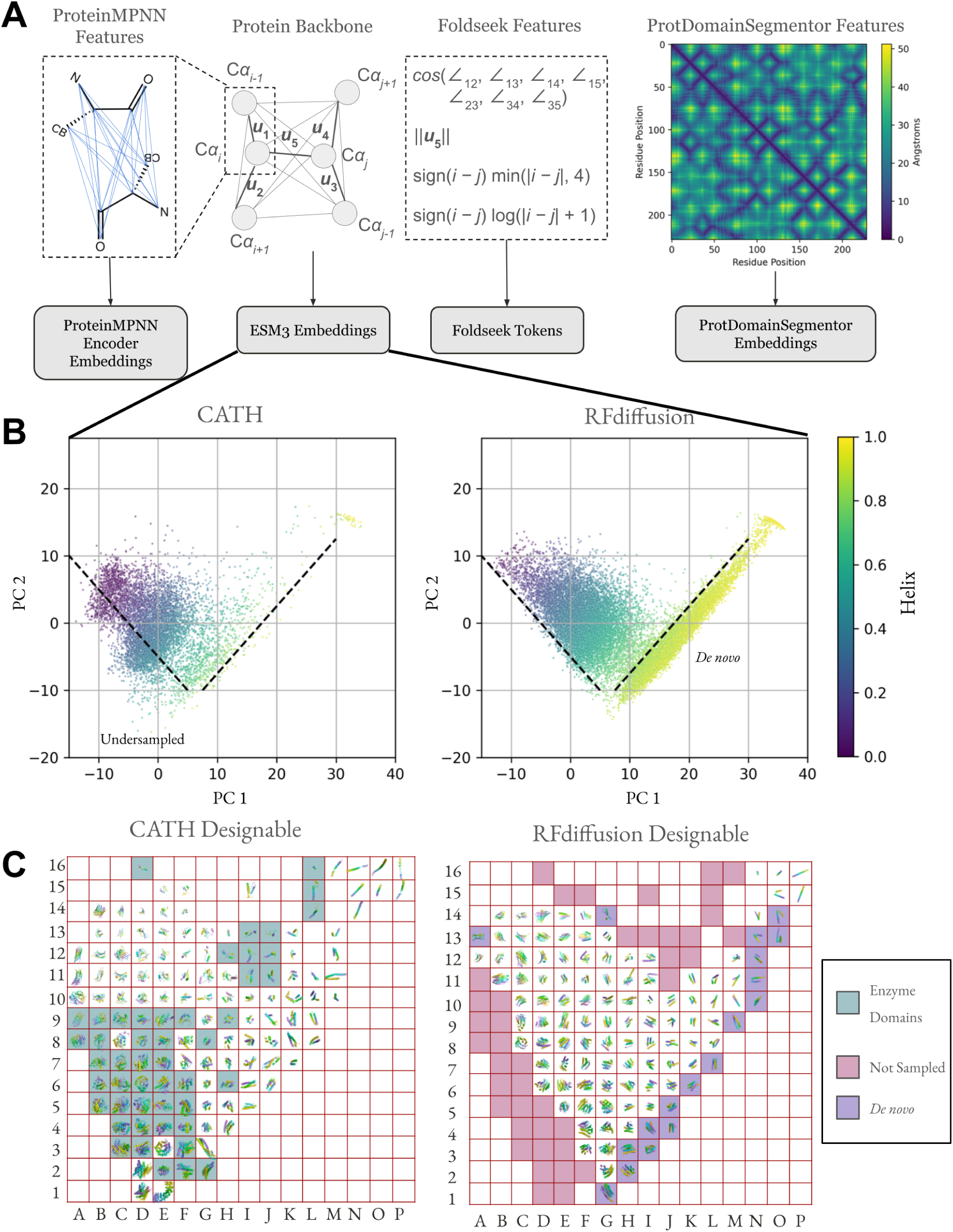
Protein structure embeddings reveal undersampled and *de novo* structure space. (A) Protein structure embeddings used in SHAPES. After mean-pooling across the sequence dimension, the dimensionality is 128 for ProteinMPNN and ESM3, 4096 for ProtDomainSegmentor. Foldseek tokens are discrete and cannot be mean-pooled, thus we count the frequency of each token in each length range (Supplementary Figure 11). (B) First two principal components of mean-pooled ESM3 embeddings colored by helix content determined by DSSP^23,24^. The indicated dashed guide lines denote visual boundaries of native structure space not sampled (Undersampled) and novel regions of protein structure space only observed in samples but not in native structures (*De novo*). (C) Rasterized visualization of panel B with 16 equally spaced grid squares in each principal component axis. A representative structure from each grid was chosen at random. Empty grid squares indicate the absence of any structure in the enclosed region. *De novo* alpha helices are shaded along the lower-right diagonal and the structures from CATH which do not have corresponding structures in the samples are shaded along the left and top rims. The structures are displayed in CATH raster plot are given in the Supplementary Information.

To better understand the distributional coverage behavior of generative models of protein structure, in particular the subspaces which samples tend to over-sample and under-sample, we drew 64,989 structures each from Chroma^11^, Genie2^12^, Protpardelle^13^, RFdiffusion^6^, and 21,663 from Multiflow^14^, matching the length distribution in Ingraham *et al*.’s CATH dataset^16,22^. We compared samples to ground truth structures from the CATH dataset as it represents a broad, expert-curated distribution of all known protein domains. We computed structure embeddings and visualized the first two principal components. The reference dataset is further filtered by resolution < 3.0 Å, R_free_ < 0.25 and no NMR structures.

Using ESM3 mean-pooled encoder embeddings, the first two principal components show a distinct streak in sampled structures not present in native CATH structures, indicating novel structural elements, along with a region present in CATH but not present in sampled structures (Figure 3B). The equivalent plots for ProteinMPNN and ProtDomainSegmentor embeddings are given in Supplementary Figure 2. We rendered the structures in Figure 3C and show that the streak is formed by idealized alpha helical structures and the undersampled region, mostly alphabeta mixtures, is enriched in enzyme domains.

We show rasterized plots for all five models stratified by designable and undesignable along with the underlying CATH distribution for ESM3 and ProtDomainSegmentor embeddings in Supplementary Figures S14 to S71. We omit ProteinMPNN raster plots for sampled structures as the sampled distribution is too narrow for structure visualization to be insightful. The designable structures are typically more concentrated and less diverse than the undesignable structures. In addition to undersampling of undesignable CATH structure space, there exist large regions of *designable* CATH structure space more enriched in beta sheets and loops such as immunoglobulins that are undersampled by all models. This pattern is consistent across models and structure embeddings. Pathologies in undesignable samples are also revealed: flexible tails with a rigid core, lever-arm effects where a flexible linker can cause large RMSD in the rigid bodies it links, poor packing, unpaired or poorly paired beta strands, an isolated beta sheet, little to no secondary structure, and chain breaks. However, not all undesignable structures exhibit visually notable pathologies and it is plausible that such undesignable samples can become designable with improved sequence design models.

### Fréchet Protein Distance (FPD) reveal undersampling of fold distributions

To quantitatively compare the distributional coverage of different models, we compute the Fréchet distance, where lower values indicate greater distribution similarity (Methods, Figure 4, Supplementary Figures 3-4). As observed in Genie2,^12^, increasing the noise injected during sampling broadens the secondary structure distribution coverage. To quantify the effect of sampling temperature, we also sample at medium and high temperature settings from each diffusion-based model 21,336 structures each, except Multiflow as an analogous parameter is not exposed to the user, (Figure 4, Supplementary Figures 3-4) and compute the FPD of each setting. The exact values of the high temperature parameter differs between models and were chosen based on the highest temperature which still can generate plausible samples by visual inspection. For Chroma, inverse temperature 3 coincides with the point beyond which the evidence lower bound starts to drop sharply (Supplementary Figure 2 in Chroma^11^). We also report the FPD computed using designable samples (FPD-D) and undesignable samples (FPD-ND). As sampling temperature increases, the sampled structures generally become more diverse as expected. Relating the effect of sampling temperature back to designability, we observe a consistent trend of fewer designable samples at higher sampling temperatures (Supplementary Figure 5-6). However, this designability-diversity trade-off is distinct from a designability-coverage trade-off: the trend in the FPD gap, defined as the difference between FPD-D and FPD-ND, is distinct between models. In Chroma, while the overall increase in diversity gives better distributional coverage, the distribution coverage of designable samples at higher temperature becomes worse. In RFdiffusion, while diversity visibly increases, we observe a mode shift which leads to worse coverage. In Genie2 and Protpardelle, coverage improves for both designable and undesignable samples. Based on the distinction between diversity vs. distribution coverage, we recommend reporting both FPD-D and FPD-ND to quantify how much of the increase in diversity is due to venturing into non-natural structure space or due to better coverage of the training data distribution, along with reporting designability-coverage trade-offs (Supplementary Figure 7).

**Figure 4.**
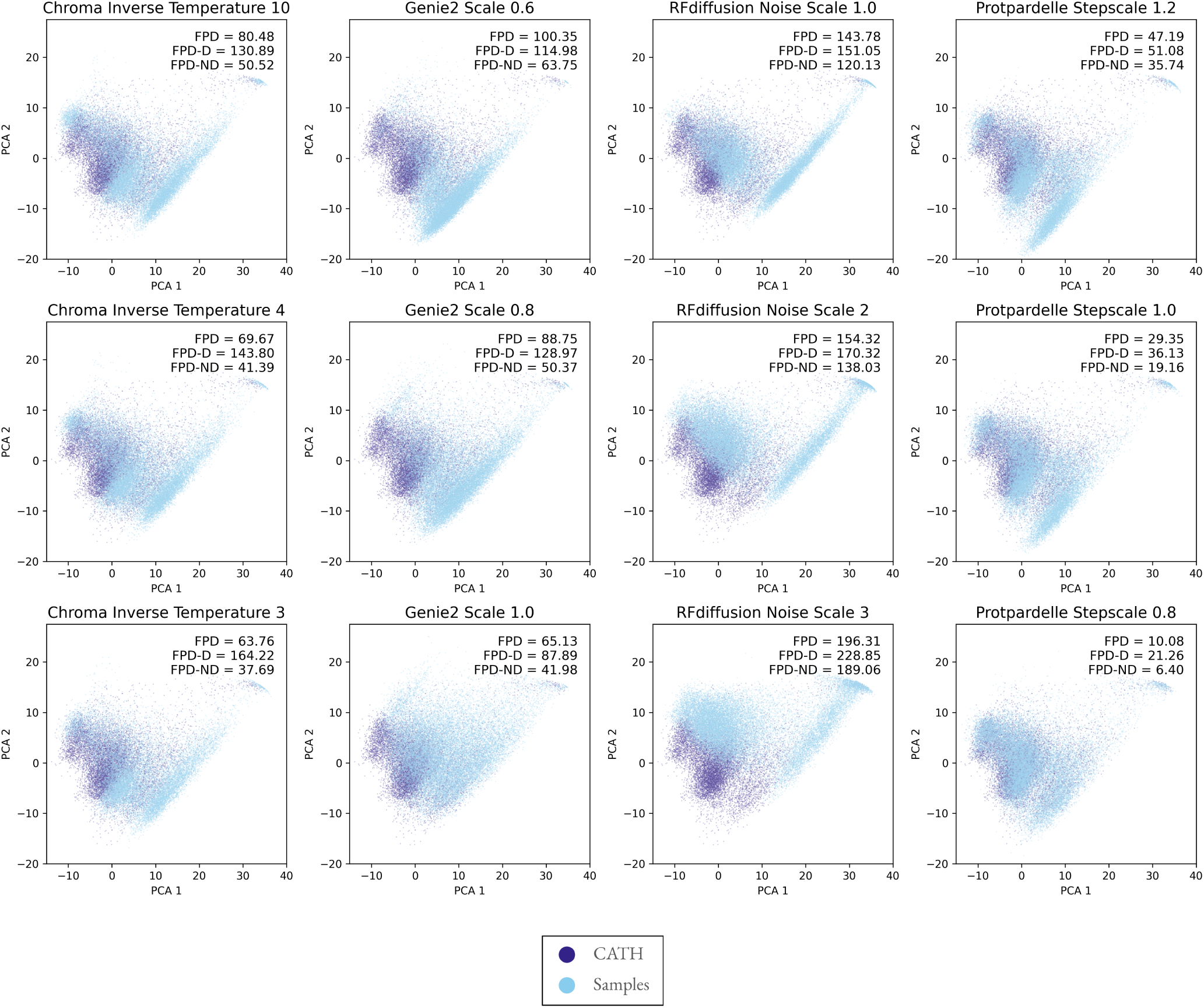
Generative models do not capture the full expressivity of PDB structures. PCA projections of ESM3 mean-pooled encoder embeddings with sampled structure projections overlayed on CATH structure projections. The streak observed in the sampled structures is not present in the native CATH distribution (Figure 3B). The top row corresponds to default sampling temperatures for each model. FPD: Fréchet Protein Distance for all samples to CATH reference set. FPD-D: FPD for designable samples with RMSD < 2.0 Å . FPD-ND: FPD for undesignable samples with RMSD > 2.0 Å . Plots for ProtDomainSegmentor and ProteinMPNN embeddings are given in Supplementary Figures 3-4. Plots for all continuous embeddings with AF3-PDB as the reference distribution instead of CATH are given in Supplementary Figures 8-10.

**Figure 5.**
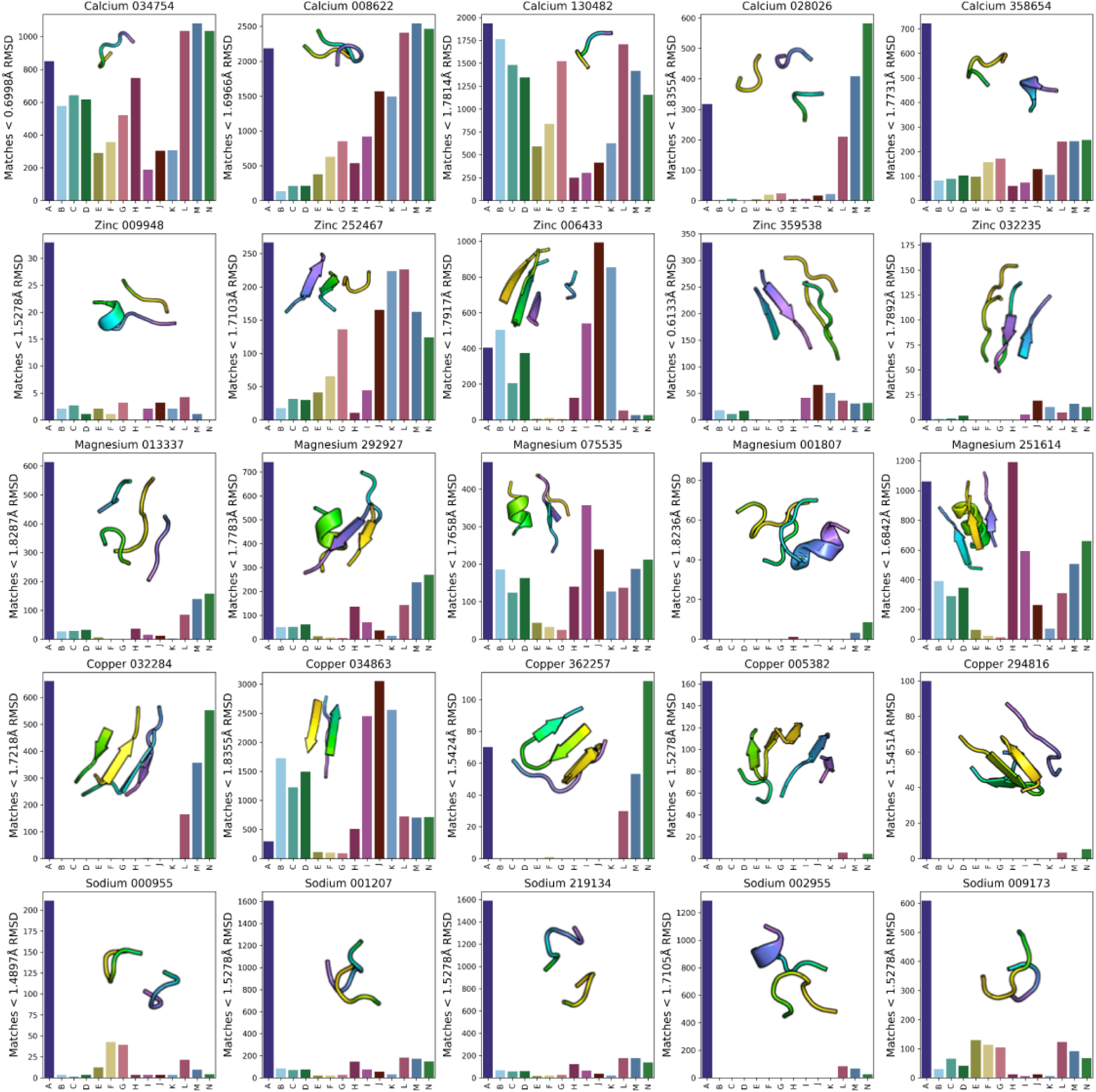
Functional tertiary structural alphabets are absent in samples. A: CATH, B: Chroma Default, C: Chroma Inverse Temperature 3, D: Chroma Inverse Temperature 4, E: Genie2 Default, F: Genie2 Scale 0.8, G: Genie2 Scale 1.0, H: MultiFlow, I: RFdiffusion Default, J: RFdiffusion Noise Scale 2, K: RFdiffusion Noise Scale 3, L: Protpardelle Stepscale 0.8, M: Protpardelle Stepscale 1.0, N: Protpardelle Stepscale 1.2. Counts of metal-binding TERMs in CATH and sampled structure sets when queried by MASTER. The RMSD threshold for each TERM depends on its complexity measured by the number of residues and the number of fragments, where more complex TERMs have less strict RMSD thresholds for a match to be counted (Methods).

While CATH provides an expert-curated distribution of diverse protein domains, solely using CATH as ground truth may be misleading as different models have different training data distributions. We compute the same FPD metrics using another reference distribution, the PDB as clustered in AlphaFold3 (AF3-PDB) (Supplementary Figures 8-10)^25^. We observe the same trends in FPD and the FPD gap across different embedding types and sampling temperatures, indicating that the biased coverage is not due to differences in CATH data leakage into the training set between models or artifacts in CATH domain parsing but rather a general behavior.

### FPD Trends Agree with Frequency Differences in Residue Nearest Neighbor Geometry

While ProteinMPNN and ESM3 capture local residue environments, strictly local geometric features may be over-smoothed given the large number of neighbors ProteinMPNN (48) and ESM3 (16) uses as context. A structure representation which uses geometric features derived from only a single nearest neighbor is Foldseek. As they are discrete, they cannot be mean-pooled. Instead, we compute similarity to a reference distribution of structures by counting the frequencies of Foldseek tokens per length range, representing the nearest-neighbor geometries commonly observed in each bin. Foldseek tokens are grouped per-token (unigram) and per pair of adjacent tokens (bigram). The KL divergence between unigram and bigram Foldseek token frequencies of CATH or AF3-PDB structures with sampled structures agree with trends in FPD using continuous embeddings, where higher sampling temperatures and noise scales give lower KL divergences. Stratifying by length ranges, we also reveal a length-dependent bias in coverage of native local structural elements (Supplementary Figure 11). Chroma, Protpardelle, and especially Multiflow (Supplementary Figure 12) have greater mismatch for longer protein lengths while Genie2 coverage is more even throughout. In particular, there is a large distribution mismatch for protein lengths below 100 amino acids in RFdiffusion samples. The coverage is not uniform across length ranges, as RFdiffusion matches the native CATH distribution more closely for proteins longer than 150 amino acids. Interestingly for RFdiffusion, increased noise scale improves coverage for proteins shorter than 150 amino acids but worsens coverage for longer proteins, in agreement with the higher FPD at higher noise scales. For all other models, the effect of sampling temperature is consistent, with higher temperature giving better coverage. The agreement with FPD trends using ProteinMPNN and ESM3 embeddings confirms that the conclusions drawn for continuous embeddings are not due to embedding artifacts but are general across embedding types. The same trends are also observed with both CATH and AF3-PDB as the reference distribution.

### Functional Tertiary Structural Alphabets are Underrepresented

The incomplete coverage protein structure space indicates that there exist tertiary structural elements present in native structures but are absent in the samples. To evaluate this, we queried sampled structure sets with recurring metal-binding TERtiary Motifs (TERMs)^15^ (Figure 5). We elect to use TERMs because they form a tertiary structural alphabet derived from a set cover of the PDB such that each TERM is a structural building block and has realizations in native structures. The rank order of TERMs is based on the amount of structural novelty each TERM introduces which we can use to gain intuition on a generative model’s ability to sample frequent vs. rare TERMs. The motifs range from short loops to double and triple loop contacts and fragments of secondary structure elements. Notably, TERMs are more interpretable than the embeddings used to compute FPD. We used MASTER^26^ to rapidly search structure sets, with a dynamic RMSD threshold as defined in Mackenzie *et al*.^15^ for a match to count.

Some TERMs, such as Calcium 034754, Calcium 008622, and Calcium 130482, are prevalent across different model samples. In contrast, some TERMs, such as Magnesium 001807, Copper 005382, Copper 294816, and Sodium 002955, are entirely absent from almost all model samples except Protpardelle. In general, Protpardelle is able to cover all TERMs, in agreement with achieving the lowest FPD of the models benchmarked. For most TERMs, the number of matches in the 21,336 CATH structures is greater than the number of matches found in the 21,336 sampled structures per model setting, indicating widespread undersampling, except for Copper 034863 which is much more oversampled by Chroma and RFdiffusion.

## DISCUSSION

To address the three questions motivating this study, using SHAPES we show that structures containing loops and loops mixed with alpha-beta structures, in which enzymes are prevalent, are not covered by most generative models. The degree to which each model covers protein structure space is quantified using FPD for comparison between different models. Specifically, RFdiffusion is optimized for highly designable samples with high secondary structure content, Genie2 generalizes to novel protein architectures while retaining consistent local geometric features with high noise scales, and Protpardelle can sample diverse loops. Distribution spread can be improved across all models by increasing sampling temperature and noise scale at the expense of designability and sometimes the coverage of the native distribution. This biased sampling imposes a limitation on the ability to sample functional structural elements.

The ability to draw unbiased samples from the protein fold space is crucial to solving motif scaffolding problems. Conditional generation is the process of drawing samples from P(scaffold | motif) = P(scaffold, motif) / P(motif) which may be computationally intractable when the likelihood of generating the motif in any scaffold is near zero. We posit that the unpredictable performance of models in motif-scaffolding benchmarks, which vary from motif to motif^6,20^, can in part be attributed to the frequency each motif is found in unconditional samples. When unconditional samples do not cover scaffolds which host motifs, especially those which exist in the PDB, conditional sampling could force the sampling trajectory out-of-distribution and generate unrealistic samples. While generating idealized *de novo* proteins is highly impactful, as demonstrated by the design of picomolar binders and highly active enzymes^7,8,27^, designing subtle but functional mechanisms inherent to natural protein structures can be challenging when sampling from non-natural or biased distributions of structures. For example, designing an enzyme that is amenable to optimization by directed evolution may be more facile with scaffolds sampled from a native structure distribution than with highly idealized scaffolds, since naturally abundant structures are products of evolution.

SHAPES offers unique advantages over using pairwise TM score or the number of clusters as diversity metrics. Mean pairwise TM score below 0.6 can be obtained trivially by mode collapse on two dissimilar folds. The number of clusters can be arbitrarily increased by generating a single alpha helix with increasing lengths. In both cases, SHAPES features are able to detect inadequate distribution coverage. Importantly, SHAPES highlights undersampled regions in which their precise identification is critical if the goal is to train a generative model which covers the full conformational diversity, thus also capturing the full functional diversity, that exists in a target data distribution. Nonetheless, current metrics of designability, diversity, and novelty are still useful as they all offer different perspectives on the performance of generative models of protein structures. The usefulness of a generative model for a protein designer depends on the objective, whether representation learning is used to predict function^28,29^, generating diversity within a single fold^30^, or extrapolating beyond natural protein structures and thereby removing the structural vestiges of evolution that hamper recombinant expression and yield. FPD is dependent on the choice of reference structures, so use cases in which the goal is to inherently sample from a custom set of structures, such as nanobody design, can be easily handled.

Designability currently guides samples towards a non-natural structure distribution. As natural structures incorporate flexible elements such as loops to achieve function, we anticipate that increased robustness in sequence design and structure prediction models will expand the space of designable protein functions such that designability can guide samples towards any desired region of protein structures while not sacrificing its predictive power on the likelihood of experimental success. As the known protein structure universe continues to expand^31^ and protein design goals become more multi-objective and ambitious, we hope that SHAPES can guide the development of the next generation of generative models of protein structures.

## LIMITATIONS

For consistency, we elect to use the standard sequence design method in the field, Protein-MPNN^4^. Use of other sequence design methods may give different designability results, in particular using the built-in sequence design models in Chroma and Multiflow. Also for consistency, we elect to use ESMFold for structure prediction instead of AlphaFold2 despite its limitations^32^. We reasoned that given the very low designability of AlphaFold2 in single sequence mode (1.34% on CATH with native sequences)^17^ and the absence of multiple sequence alignments to run AlphaFold2 in MSA-mode for ProteinMPNN-designed sequences, a fast and relatively robust single-sequence model such as ESMFold is most appropriate. Different structure-prediction models would also give different designability results. Results for the coverage behavior of conditional sampling may be different than the unconditional setting analyzed here due to different forms of guidance used during sampling and cases in which an unconditional model is fine-tuned to obtain a conditional model. We leave analysis of conditional samples to future work.

While SHAPES does not rely on the sequence design capability of ProteinMPNN, it nonetheless relies on representations learned from models trained for unrelated tasks. An alternative is to use embeddings directly extracted from a diffusion model^3,33^. Other embeddings can be used, as long as they can capture features which can discern samples apart, such as using LigandMPNN^34^ embeddings to evaluate generative models of all-atom protein structures.

## METHODS

### SHAPES

Using SHAPES consists of the following steps:

1. Choose a target training data distribution. Here we choose CATH^16^ and AF3-PDB^9^.
2. Sample structures at varying lengths from a generative model. Here we match the length distribution of the reference dataset.
3. Compute structure embeddings: Foldseek, ProteinMPNN, ESM3, and ProtDomainSegmentor
4. Compute Fréchet Protein Distance (FPD) for each continuous embedding and KL-divergence for Foldseek tokens.
5. Visualization of distribution coverage.

### Fréchet Protein Distance (FPD)

Given *N* data points from a ground truth distribution *p*_data_(**x**) and *M* samples from a model *p*_sample_(**x**), we compute embeddings 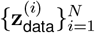 and 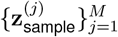. The distributional similarity of sampled structures to reference structures can be quantified by computing the Fréchet Protein Distance (FPD) given by Equation 1.

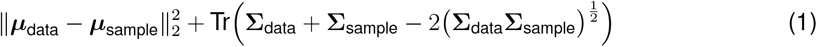

where ***µ***_data_ and ***µ***_sample_ are the mean vectors of the reference structures and the sampled structures respectively, and **Σ**_data_ and **Σ**_sample_ are the covariance matrices of the reference structures and the sampled structures respectively. In practice, fewer samples, on the order of 2000 can be used to estimate the FPD computed with a large number of samples, on the order of 20000 (Supplementary Figure 13).

### TERMs

The metal-binding motifs are from Figure S9 of Mackenzie *et al*.^15^. The formula used to compute the RMSD threshold for each motif is given by Equation 1 of Mackenzie *et al*.:

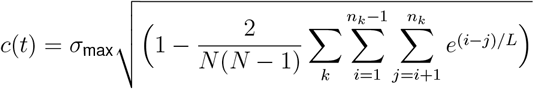

where *N* is the number of residues, *k* indexes the segment, such that the *n*’th segment has length *n*_*k*_, *L* is a correlation length set to 20 as recommended. Here, *σ*_max_ = 2.0 which is double the recommended value as the recommended value of 1.0 gave zero matches for most models for most motifs. *σ*_max_ = 1.0 for Calcium 130482 and *σ*_max_ = 1.5 for Copper 034863 as more than 5000 matches were found with *σ*_max_ = 2.0.

## Data and code availability

- All original code has been deposited at the GitHub repository https://github.com/ProteinDesignLab/protein_shapes and data generated has been deposited at Zenodo under the DOI 10.5281/zenodo.14166398.

## ACKNOWLEDGMENTS

The authors thank Richard Shuai for comments on the manuscript. T.L is supported by the Stanford Graduate Fellowship. M.L. is supported by Stanford Bioengineering department’s Summer Research Experiences for Undergraduates (REU) program. P.-S.H. is supported by NIH (R01GM147893).

## AUTHOR CONTRIBUTIONS

Conceptualization, T.L., J.K., P.-S.H; methodology, T.L., M.L. and Y.C.; data collection, T.L., M.L.; manuscript writing, T.L., M.L. and P.-S.H.; supervised research, P.-S.H.

