## Supplementary Material for "Assessing Generative Model Coverage of Protein Structures with SHAPES"

#### CATH structures in Figure 3C

A8: 1jtdB00 hydrolase/inhibitor; A9: 3r0sA01 transferase; A10: 2w38A01 transferase; A11: 3ljyA00 cell adhesion; A12: 4rlcA00 transport protein; B5: 1ga6A00 hydrolase/hydrolase inhibitor; B6: 1ejdA01 transferase; B7: 1deuB00 hydrolase; B8: 4b5qA00 hydrolase; B9: 1ee6A00 lyase; B10: 2w47A00 hydrolase; B11: 2vm9A02 cell adhesion; B12: 2xwxA03 chitin-binding protein; B13: 3rbsA02 contractile protein; B14: 4fqeA00 membrane protein; C3: 1gaiA00 hydrolase; C4: 2dc0A00 hydrolase; C5: 3eu8A00 hydrolase; C6: 4u4eA00 transferase; C7: 4x9mA01 oxidoreductase; C8: 1j31A00 structural genomics, unknown function; C9: 3fe4B00 lyase; C10: 4g1vA02 oxidoreductase; C11: 1ylnA01 structural genomics, unknown function; C12: 2ichA01 lipid binding protein; C13: 2cg7A01 signaling protein; C14: 3pqhA01 viral protein; D1: 2b2hA00 transport protein; D2: 3hd6A00 membrane protein, transport protein; D3: 4ozvA00 lyase; D4: 1szqA01 structural genomics, lyase; D5: 3qxfA00 hydrolase; D6: 2yxoB00 hydrolase; D7: 1dkuA transferase; D8: 3m84A02 ligase; D9: 2orwB01 transferase; D10: 2w9hA00 oxidoreductase; D11: 4hwmA00 structural genomics, unknown function; D12: 3le2A02 hydrolase; D13: 2retA00 protein transport; D14: 3kvpA00 structural genomics, unknown function; D16: 3ksrA01 hydrolase; E1: 4rv1D00 de novo protein; E2: 3iv7A02 oxidoreductase; E3: 1nc5A00 structural genomics, unknown function; E4: 3qfeB00 lyase; E5: 4iw7A02 transferase; E6: 1e7wB00 oxidoreductase; E7: 5swuA00 hydrolase; E8: 4j37A02 rna binding protein; E9: 3cc8A00 transferase; E10: 3a7fA01 transferase; E11: 3i1aA01 transferase; E12: 5cemA01 transferase; E13: 1mhxA00 immune system; E14: 1o9yC00 structural protein; E15: 3tndB01 translation, toxin; F2: 2j1oA00 transferase; F3: 1tazA00 hydrolase; F4: 3h2zA02 oxidoreductase; F5: 1yisA02 lyase; F6: 4zb7A00 transferase; F7: 3eeaA00 structural genomics, unknown function; F8: 4ljyA02 hydrolase; F9: 1lq9A00 oxidoreductase; F10: 2dsyA00 structural genomics, unknown function; F11: 4iajA00 structural genomics, unknown function; F12: 2cjsC01 exocytosis; F13: 2fe3A02 dna binding protein; F14: 3zoqC00 hydrolase/viral protein; F15: 1mvfD00 immune system; G2: 3l39A01 phosphate-binding protein; G3: 4e40A00 transport protein; G4: 3l9tA02 unknown function; G5: 1w98B02 transferase; G6: 3fymA00 dna binding protein; G7: 2f5jB00 gene regulation; G8: 5f33A03 transferase; G9: 3c3wA01 transcription; G10: 5trbA00 ligase; G11: 3r6fA00 hydrolase; G12: 3mwmA02 transcription; G13: 2i8dA01 structural genomics/unknown function; H4: 3bt5A00 structural genomics, unknown function; H5: 3qsgA02 structural genomics, unknown function; H6: 1vbiA01 oxidoreductase; H7: 1hbkA00 fatty acid metabolism; H8: 3i4uA01 hydrolase; H9: 1vquA01 transferase; H10: 2vt1B00 membrane protein; H11: 1ayeA01 serine protease; H12: 4mouA02 isomerase; H13: 1a1iA01 transcription/dna; I5: 1tqgA00 transferase; I6: 4egwA02 metal transport; I7: 4bwcA02 hydrolase; I8: 4yifF00 dna binding protein; I9: 3lsgA01 transcription regulator; I10: 4csrB00 transcription; I11: 4nekA02 isomerase; I12: 3ephA03 transferase/rna; I13: 1gyxA01 isomerase; I15: 4g6qA02 structural genomics, unknown function; J6: 4a64C03 cell cycle; J7: 1u5pA01 structural protein; J8: 3v7dD01 cell cycle; J9: 1a7wA00 histone; J10: 1e3oC02 transcription; J11: 4wczA02 isomerase; J12: 3ip4C01 ligase; J13: 1vloA03 transferase; K7: 2x2vA00 membrane protein; K8:

2fcwA00 lipid transport/endocytosis/chaperone; K9: 2f1kC02 oxidoreductase; K10: 3mhsB00 hydrolase/transcription regulator/protein binding; K11: 4wv4B00 transcription; K12: 2c9wA02 transcription regulation; K13: 3ufeB01 transcription; L8: 2ic6A00 viral protein; L9: 1lrzA03 antibiotic inhibitor; L10: 4adzA00 transcription; L11: 1j2jB00 protein transport; L14: 5b1aL00 oxidoreductase; L15: 5b1aJ00 oxidoreductase; L16: 3r4yA01 hydrolase; M10: 2qtfA02 nucleotide binding protein; M11: 3okqA00 protein binding; M12: 3hl8A04 hydrolase; M13: 2gpeD01 dna binding protein; M16: 1g1iA00 metal binding protein; N14: 2xqhA02 cell adhesion; N15: 2h8pD00 membrane protein; N16: 4w7yA00 transport protein; O15: 2ymyB00 apoptosis; O16: 3b5nK00 membrane protein; P15: 4wy4D00 membrane protein; P16: 2w6aB00 signaling protein.

#### Supplementary Figures


Figure S1: Secondary structure elements distribution of designable vs. undesignable CATH structures.

Figure S2: First two principal components of ProteinMPNN and ProtDomainSegmentor embeddings of CATH and RFdiffusion structures colored by helix content.

Figure S3: Coverage of CATH protein structure distribution visualized with ProtDomainSegmentor mean-pooled pre-final layer embeddings.

Figure S4: Coverage of CATH protein structure distribution visualized with ProteinMPNN final encoder layer embeddings.

Figure S5: Length-dependence of designability of all sampled structures stratified by temperature.

Figure S6: Designability of all sampled structures stratified by temperature and pLDDT thresholds.

Figure S7: Designability-coverage tradeoff.

Figure S8: ESM3-based FPD and coverage of AF3-PDB.

Figure S9: ProtDomainSegmentor-based FPD and coverage of AF3-PDB.

Figure S10: ProteinMPNN-based FPD and coverage of AF3-PDB.

Figure S11: Coverage of Foldseek token distribution of CATH structures measured by Kullback-Leibler divergence of unigram and bigram token probabilities.

Figure S12: Multiflow coverage.

Figure S13: Errors in estimate of true FPD by sampling fewer structures.

Figure S14 to S71: Rasterized plot with structures spatially arranged by the first and second principal components of ProtDomainSegmentor, ProteinMPNN encoder, and ESM3 latent space.

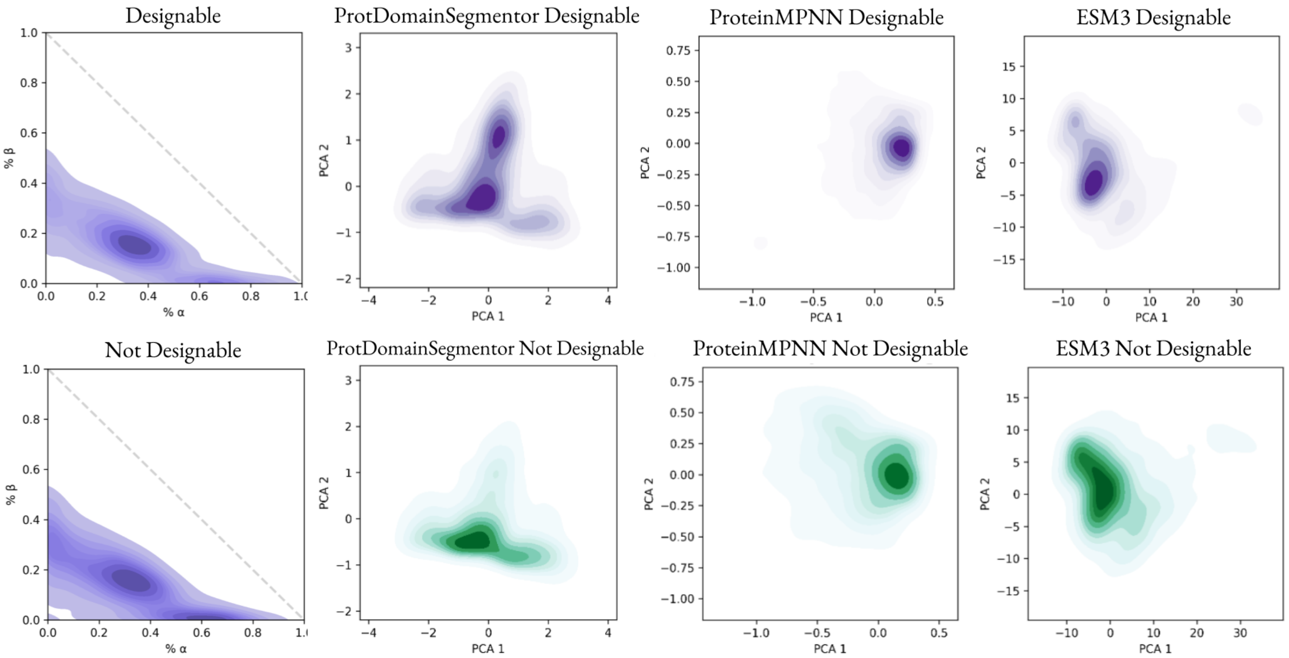


**Supplementary Figure 1. Secondary structure elements distributions do not explain designable vs. undesignable structures.** Secondary structure element frequencies computed with DSSP for CATH structures. While the distribution of undesignable structures involves additional density for structures with few secondary structures, there is no pronounced right-upward shift of the designable distribution as in the sampled structures. Kernel density estimates of embeddings of CATH structures shown more pronounced differences of designable vs. undesignable structure space.

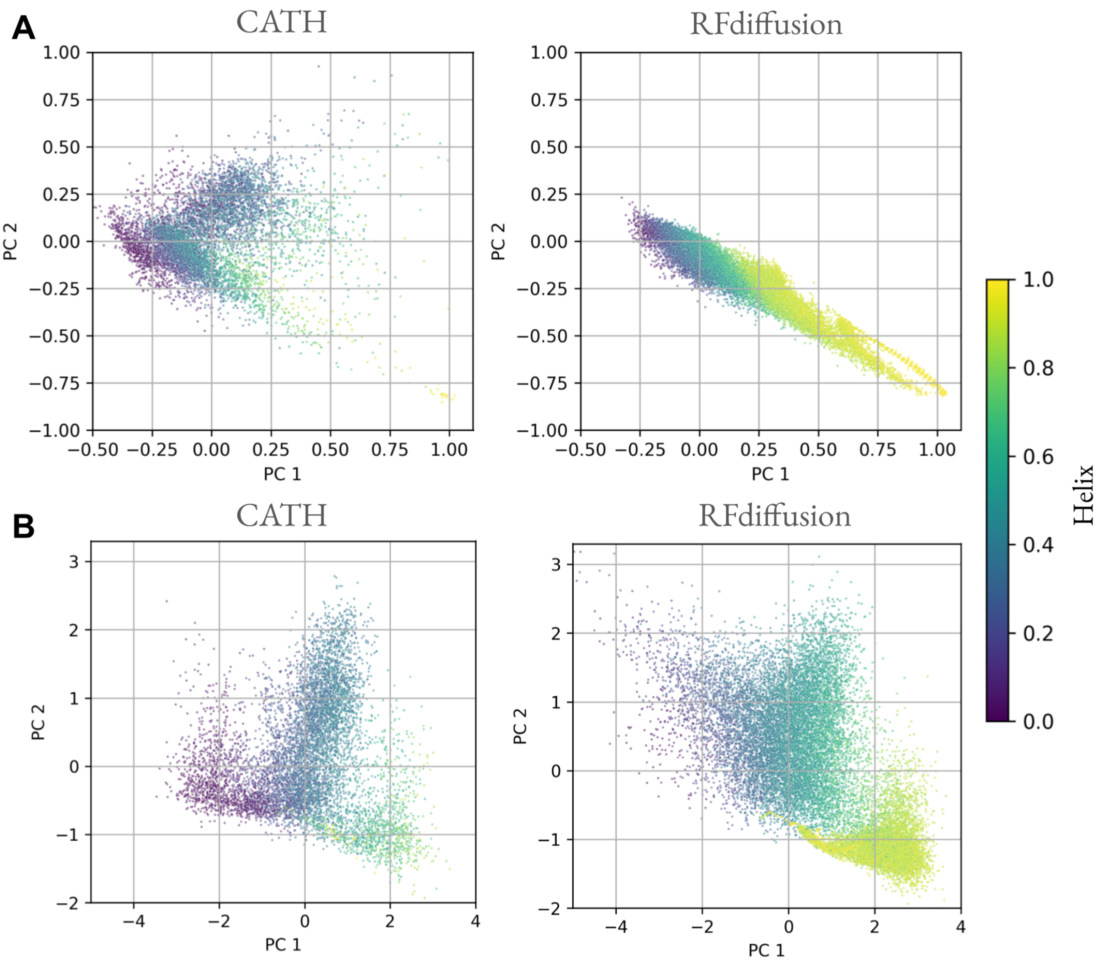


**Supplementary Figure 2. CATH and RFdiffusion distributions colored by helix content.** (A) First two principal components of mean-pooled ProteinMPNN encoder final layer embeddings. The *de novo* structures are prominent in the lower-right of the RFdiffusion plot with the same gradient in helix content as present in the ESM3 encoder embeddings. (B) First two principal components of mean-pooled ProtDomainSegmentor pre-final layer embeddings.

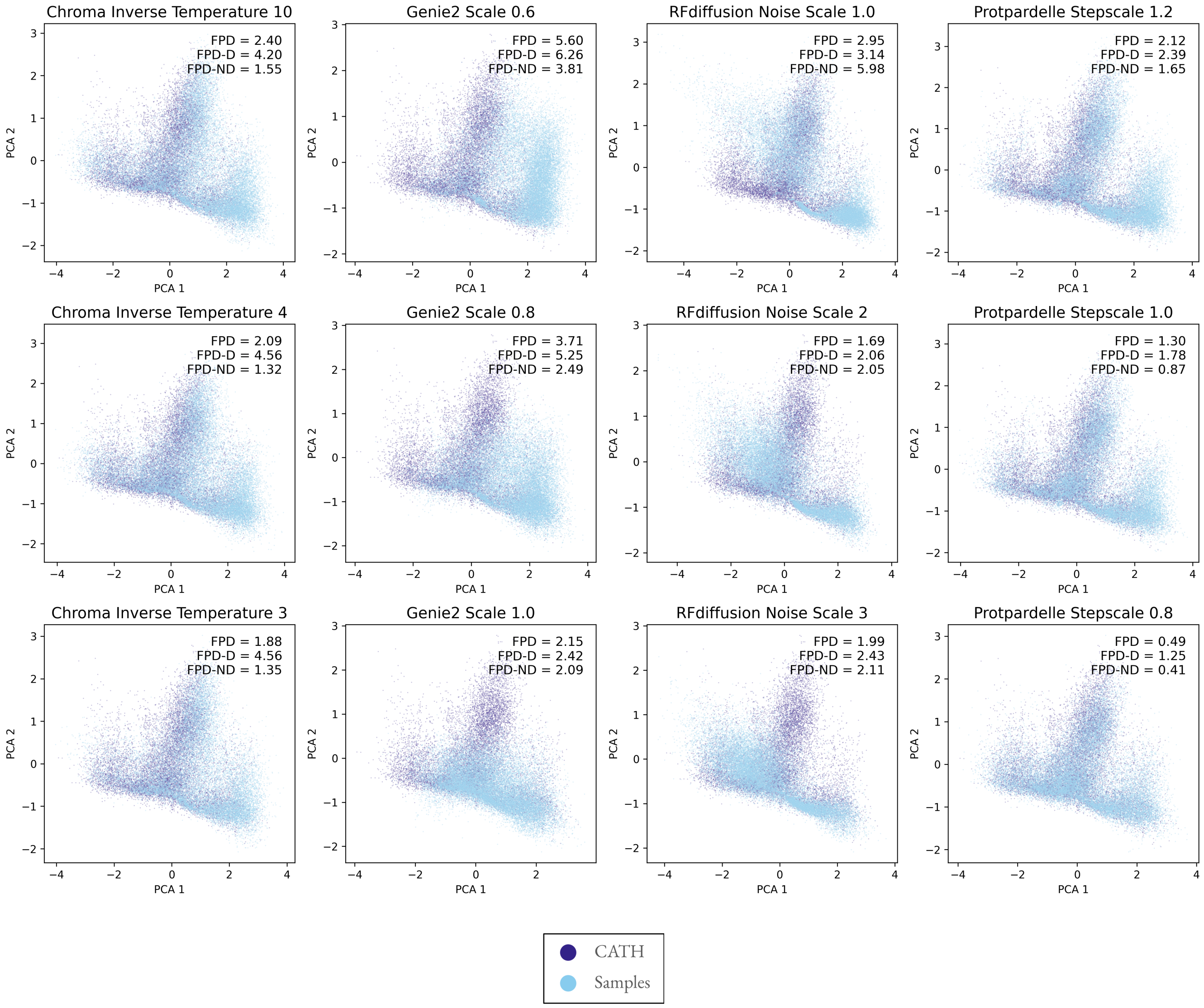


**Supplementary Figure 3. ProtDomainSegmentor feature coverage is more complete.** The pre-final layer embeddings from ProtDomainSegementor organize protein architectures by similarity as it is directly projected to predict the CATH architecture of an input protein structure. Sampled structures which do not overlay on the dark background of CATH structures indicate folds which are not observed in CATH architectures.

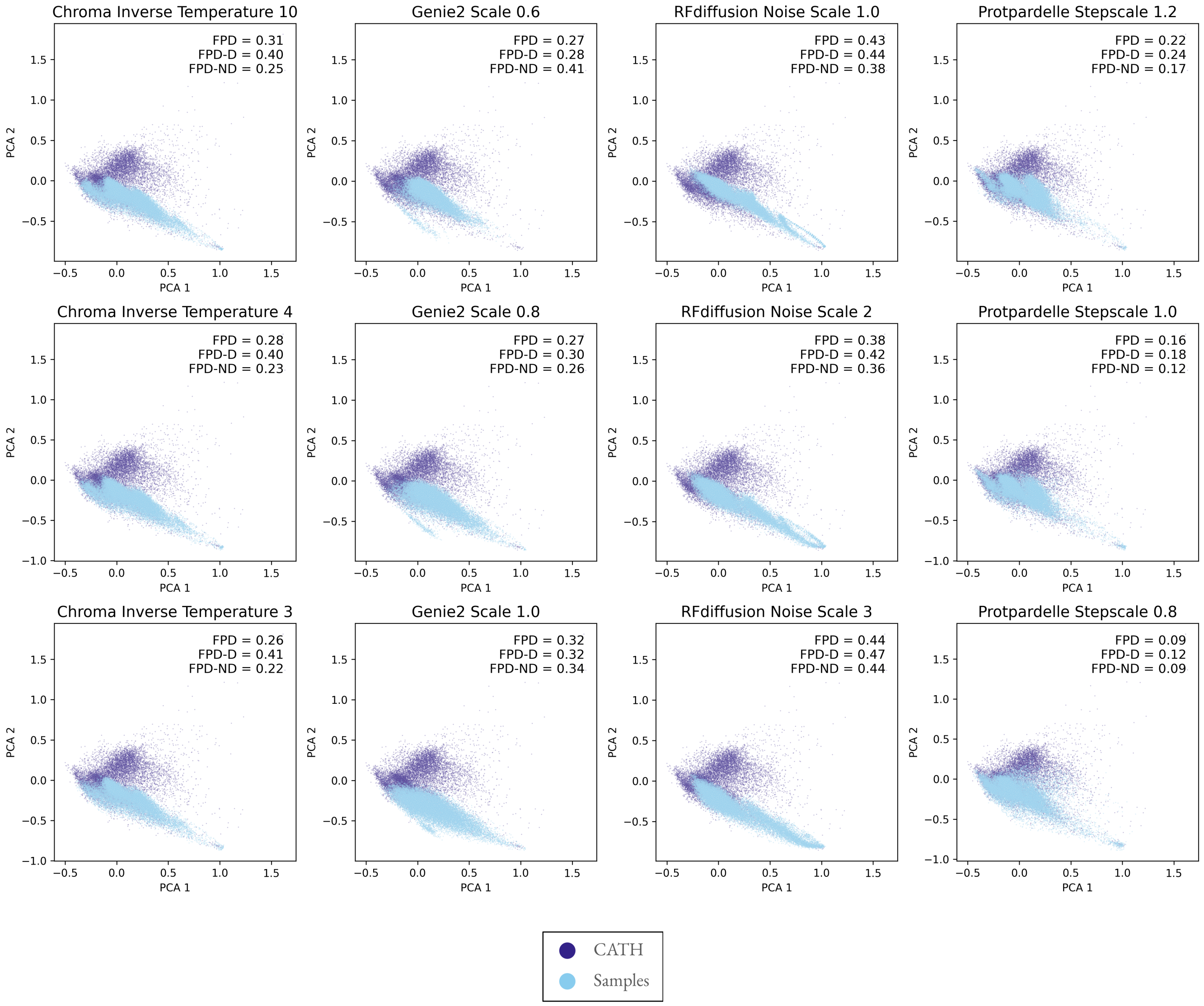


**Supplementary Figure 4. Local amino acid environments of native structures are not fully captured by sampled structures** First two principal components of mean-pooled ProteinMPNN final encoder layer embeddings show that samples do not fully cover local amino acid environment features observed in native structures.

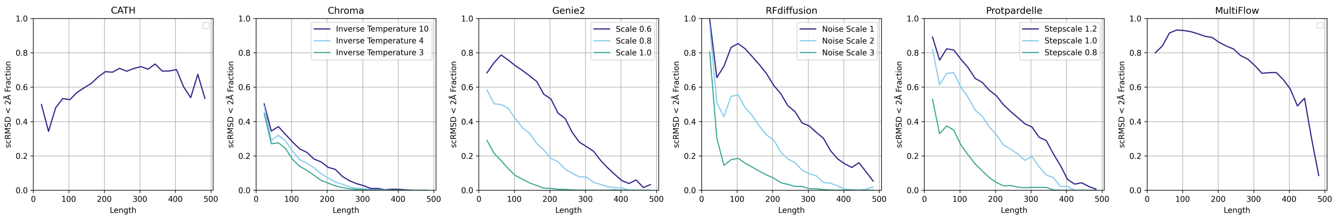


**Supplementary Figure 5. Increased sampling temperature consistently reduces designability of samples.** For comparison of generated backbones versus native backbones, the CATH designability reported here is computed from 32 ProteinMPNN sequences designed given the native backbone, not native sequences. A structure is designable if RMSD $<$ 2Å and pLDDT $>$ 80.

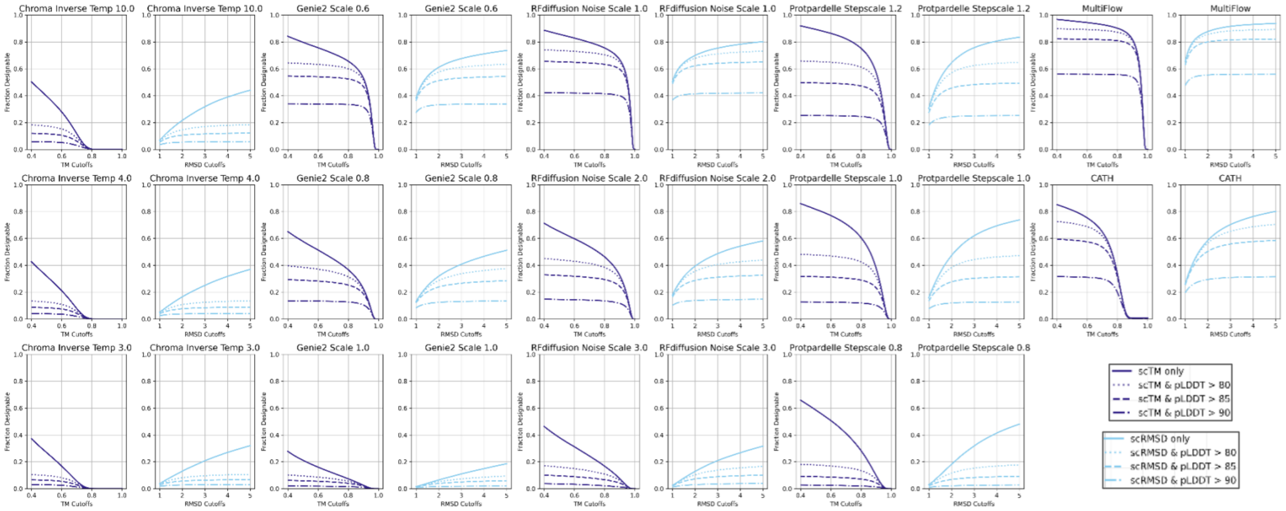


**Supplementary Figure 6. Threshold-dependent and pLDDT-dependent designability statistics.** An additional pLDDT threshold is often used to filter designs and included in the designability metric alongside scRMSD or scTM. Visualizing the effect of different scRMSD and scTM cutoffs stratified by pLDDT thresholds reveal different behaviors: Multiflow produces highly designable backbones with few filtered out by an additional pLDDT thresholds. Conversely, including a pLDDT threshold filters out more Protpardelle generated structures while more samples pass scTM-only and scRMSD-only thresholds. The designable fraction is reported across all lengths.

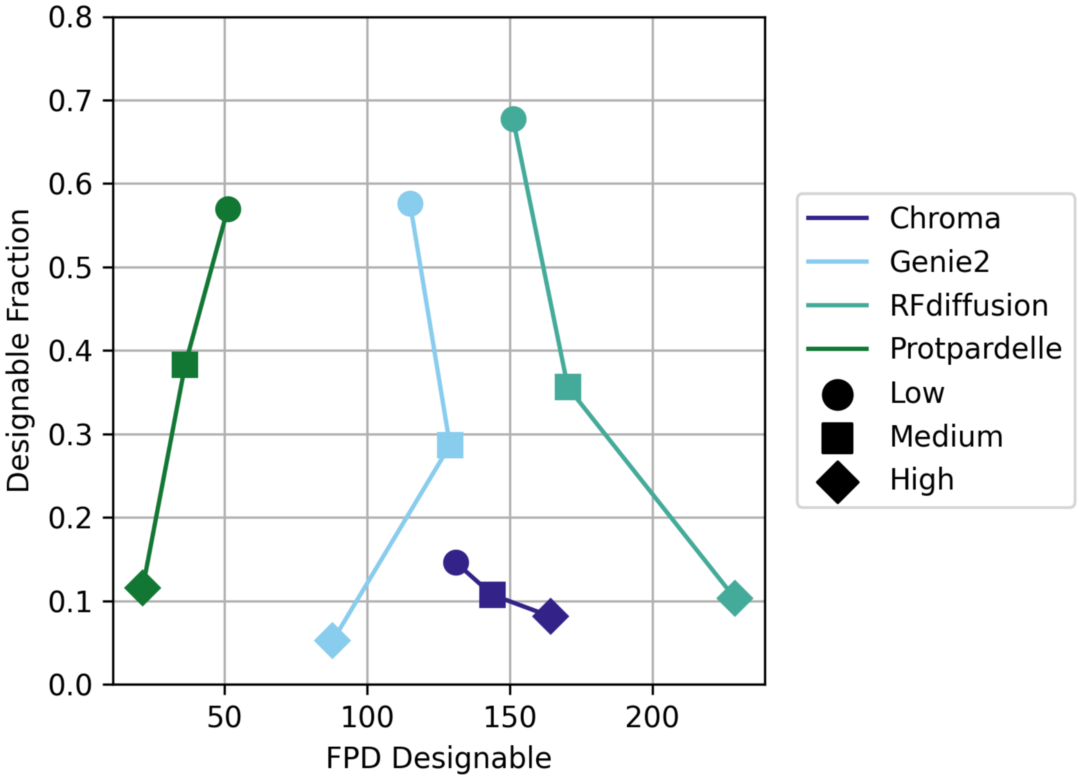


**Supplementary Figure 7. Trade-off between designability and coverage.** For all models, higher sampling temperature leads to lower designability. Only for Protpardelle and Genie2 does it also lead to better coverage, whereas the coverage becomes worse for Chroma and RFdiffusion. The FPD is reported for only the designable subset of sampled structures.

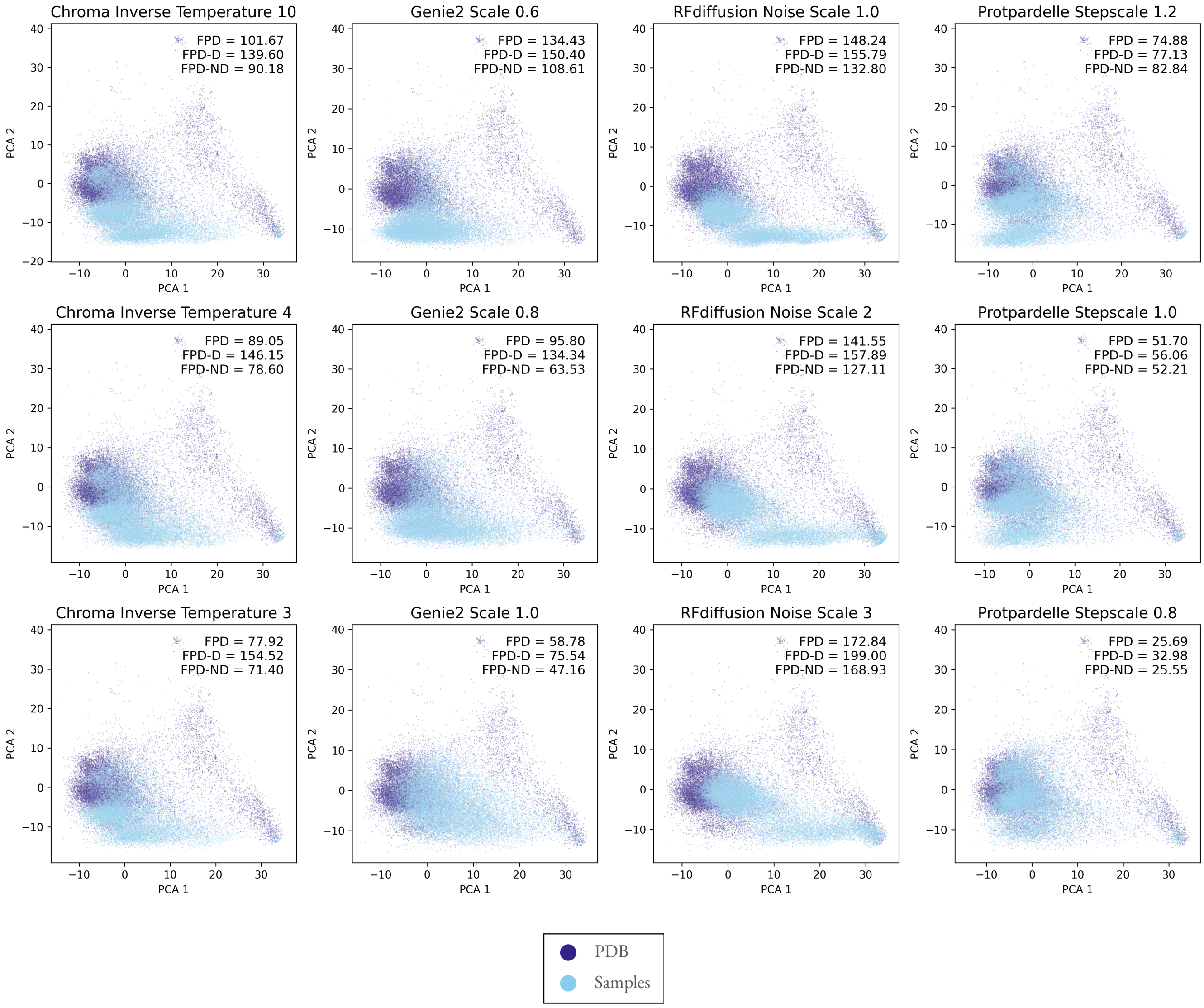


**Supplementary Figure 8. ESM3-based FPD and coverage of AF3-PDB.** While changing the reference distribution results in a different organization of the first two principal components, the streak of de novo helical structures is still prominent.

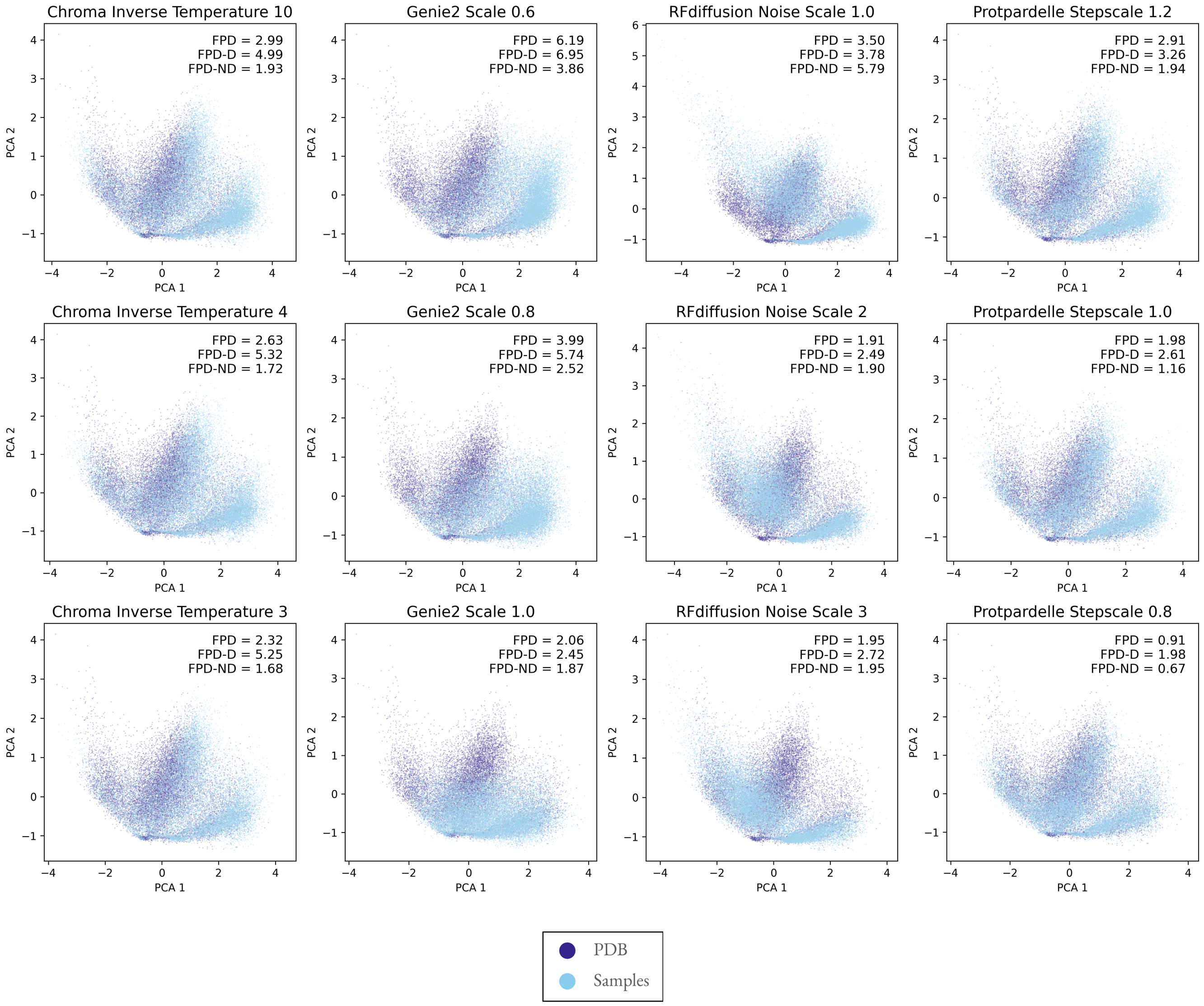


**Supplementary Figure 9. ProtDomainSegmentor-based FPD and coverage of AF3-PDB.** Consistent with CATH as the reference structure set, coverage of protein domains is generally better than coverage of local amino acid environments across all models.

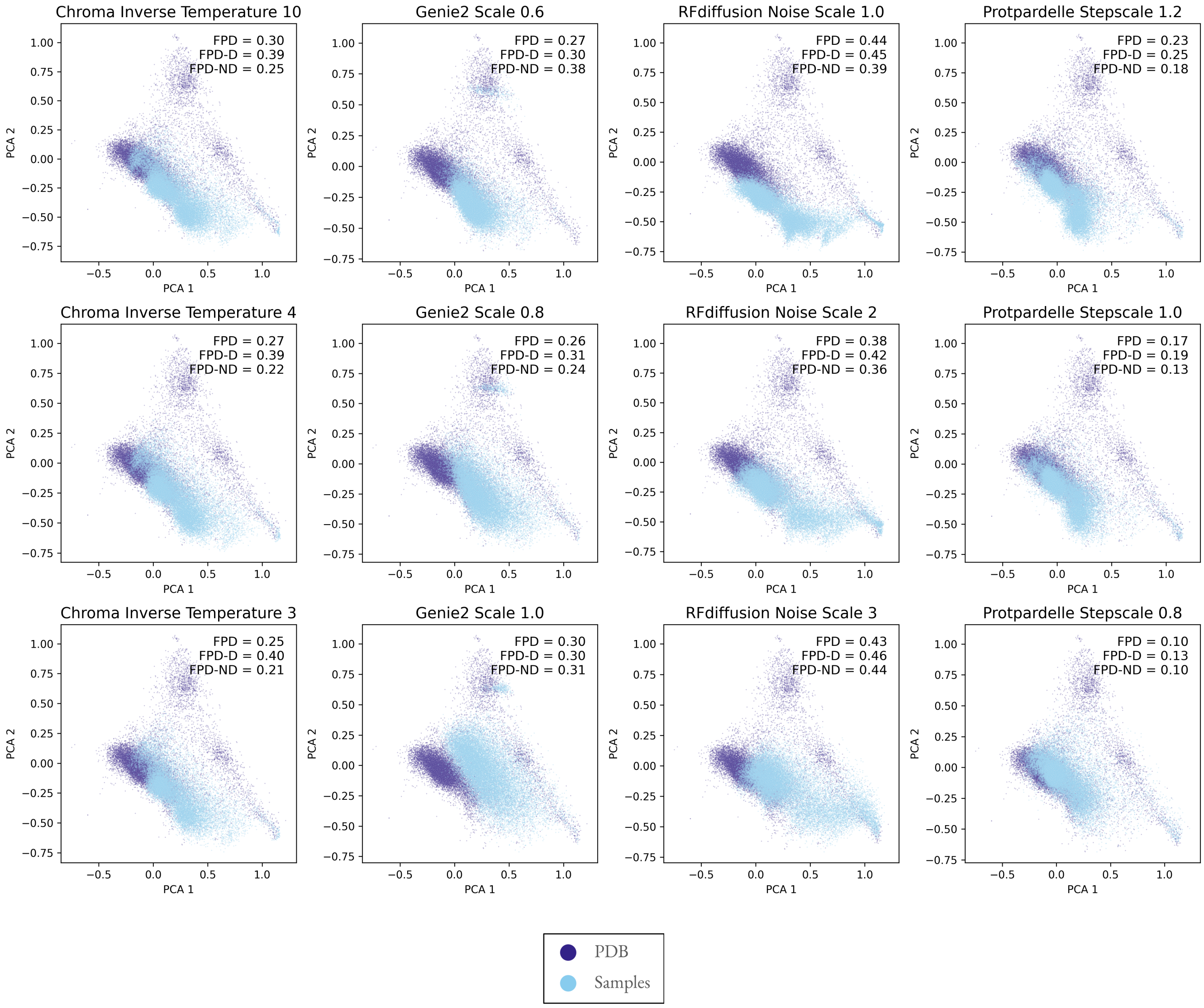


**Supplementary Figure 10. ProteinMPNN-based FPD and coverage of AF3-PDB.** The difference in FPD magnitudes compared to ESM3 embeddings is due to the magnitudes of the structure embeddings. Structural coverage of ProteinMPNN features in AF3-PDB is incomplete.

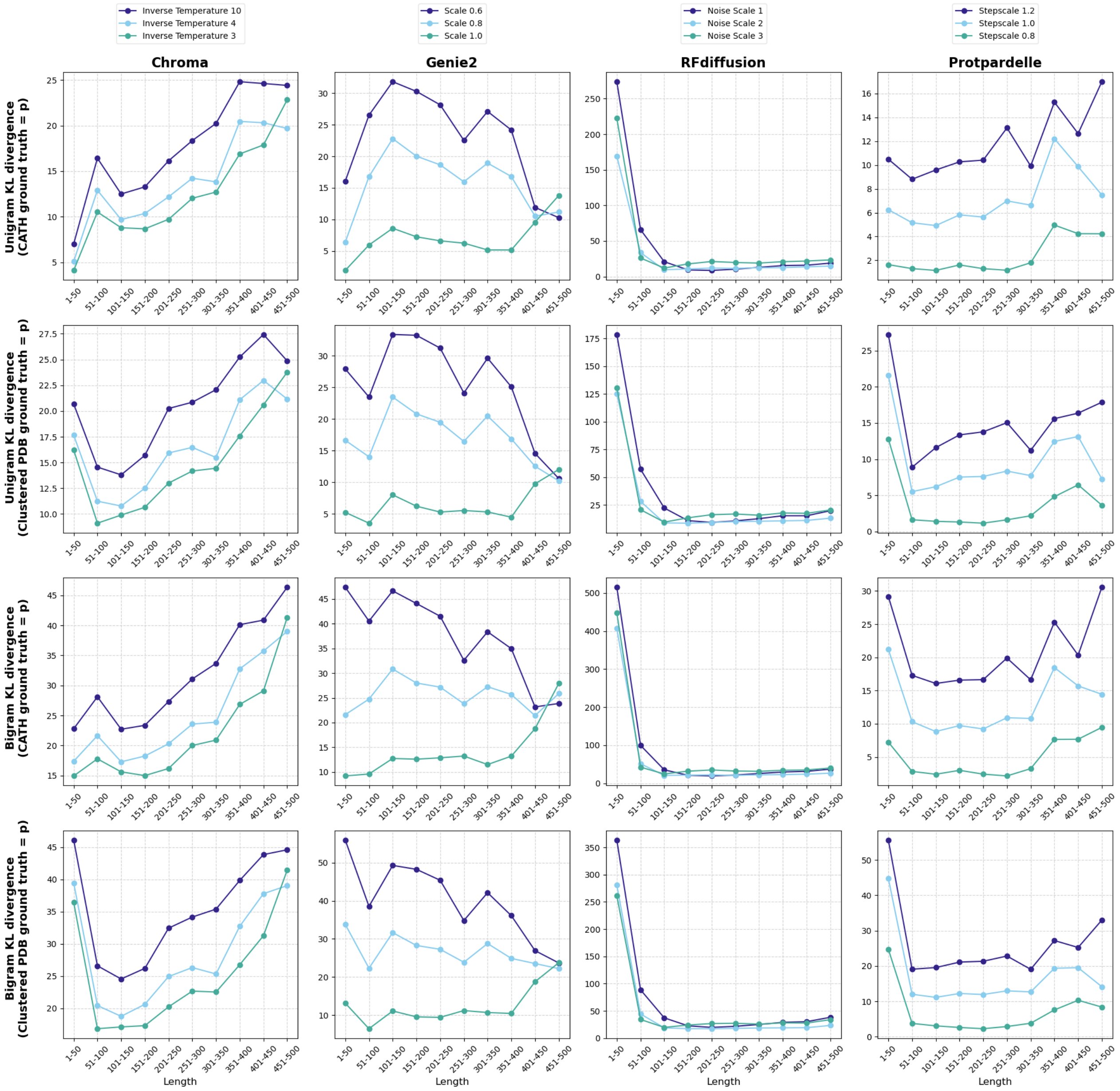


**Supplementary Figure 11. Foldseek-derived nearest neighbor residue geometric features are undersampled and display length-dependent bias.** KL(data||sample) of unigram and bigram Foldseek token frequencies. Lower values indicate the modes in the data are covered by samples while higher values indicate missing mode coverage. KL: Kullback-Leibler divergence.

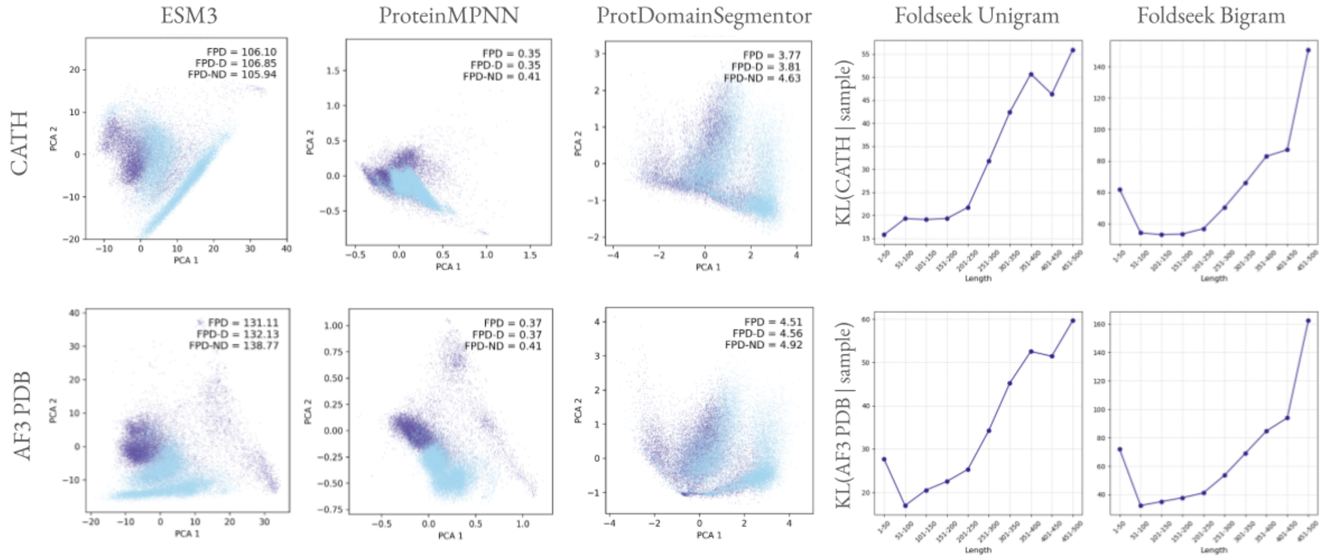


**Supplementary Figure 12. Multiflow coverage.** The distinct streak in ESM3 embeddings is also present in Multiflow samples. Similar to other models, coverage of protein architectures is generally better than the coverage of local features. Interestingly, Foldseek tokens show worse coverage of native nearest-neighbor features at longer lengths.

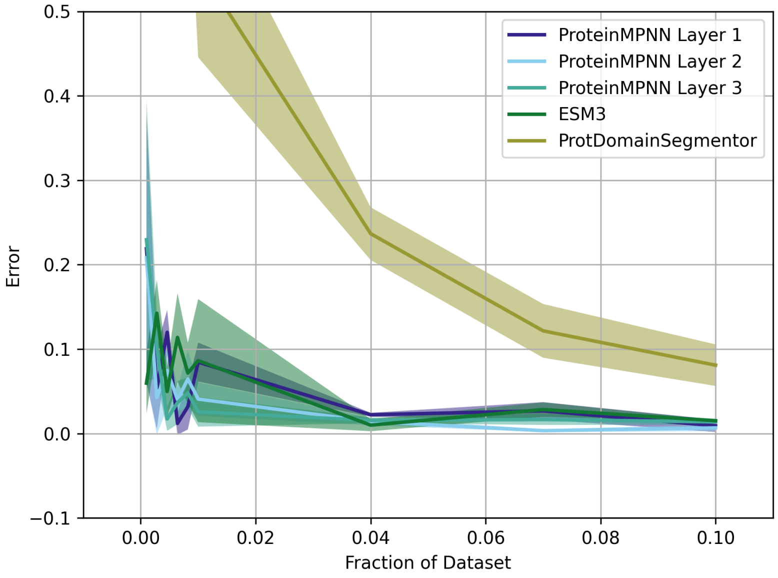


**Supplementary Figure 13. Fewer samples can be used to estimate FPD.** To reduce computational demand of computing FPD, 10% of the full set of samples, 2,166 out of 21,663 structures, can be used to estimate the FPD computed using the full set.

CATH Designable

ProtDomainSegmentor PCA

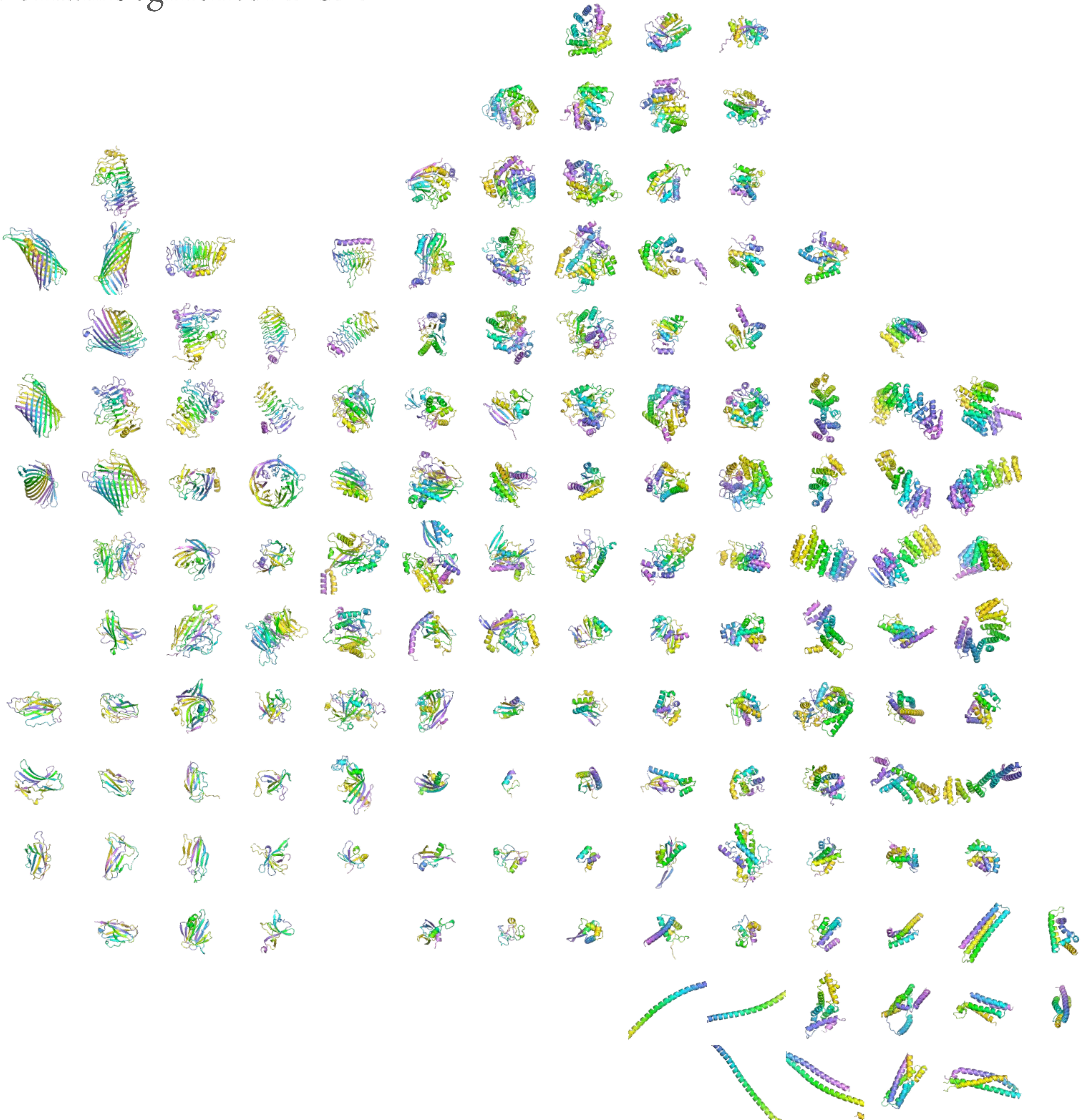

CATH Undesignable

ProtDomainSegmentor PCA

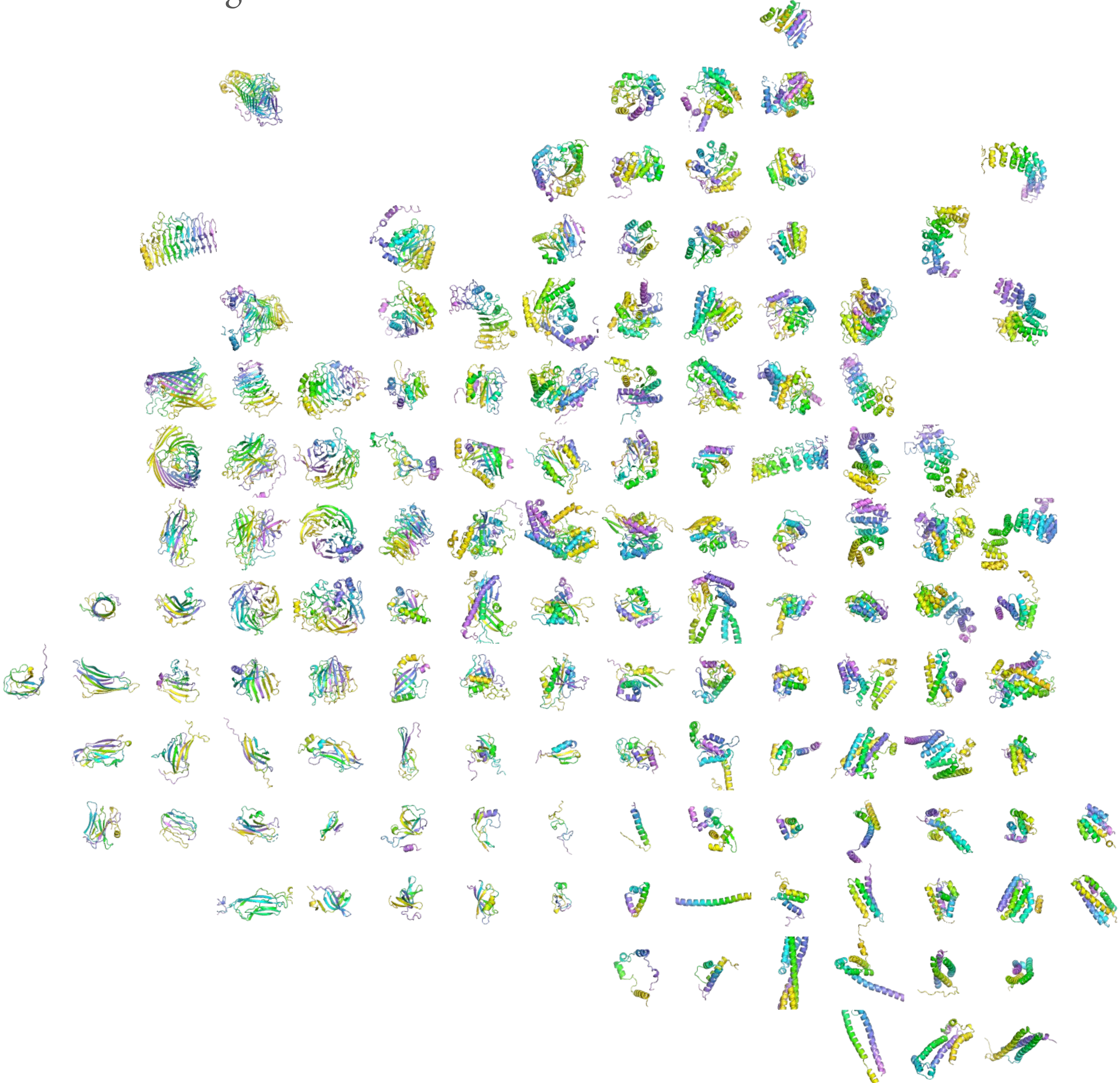

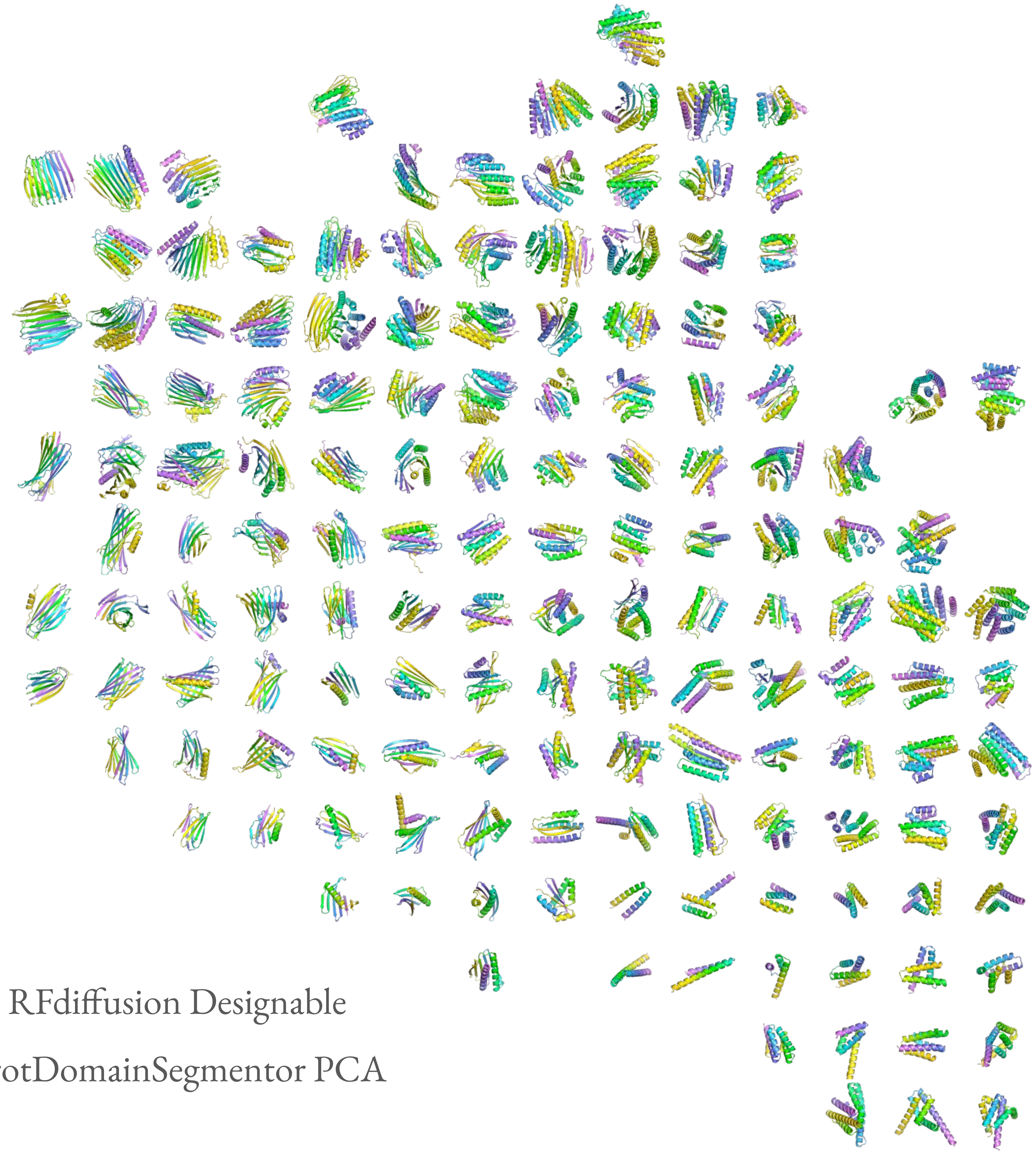

RFdiffusion Designable

ProtDomainSegmentor PCA

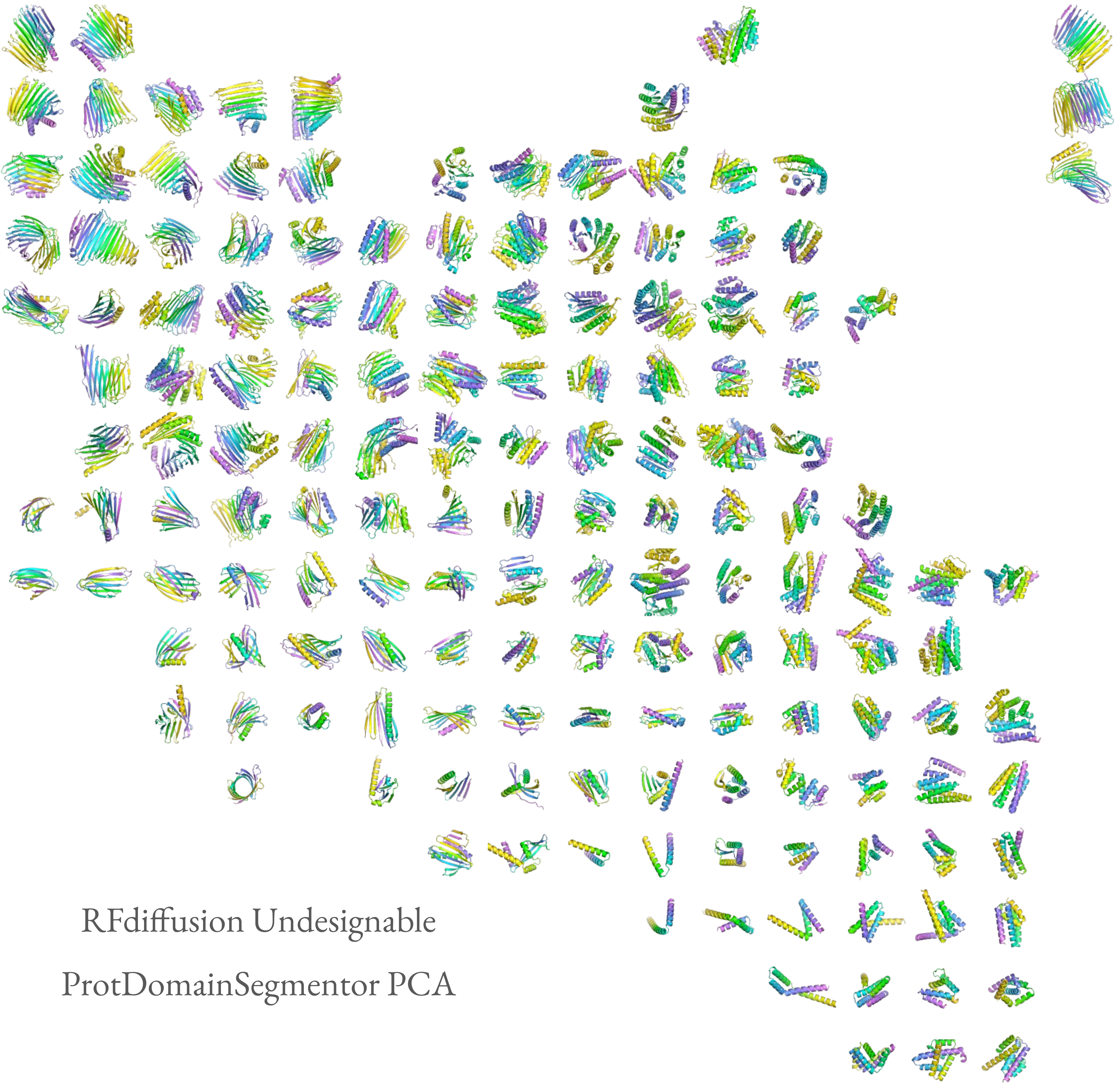

RFdiffusion Undesignable

ProtDomainSegmentor PCA

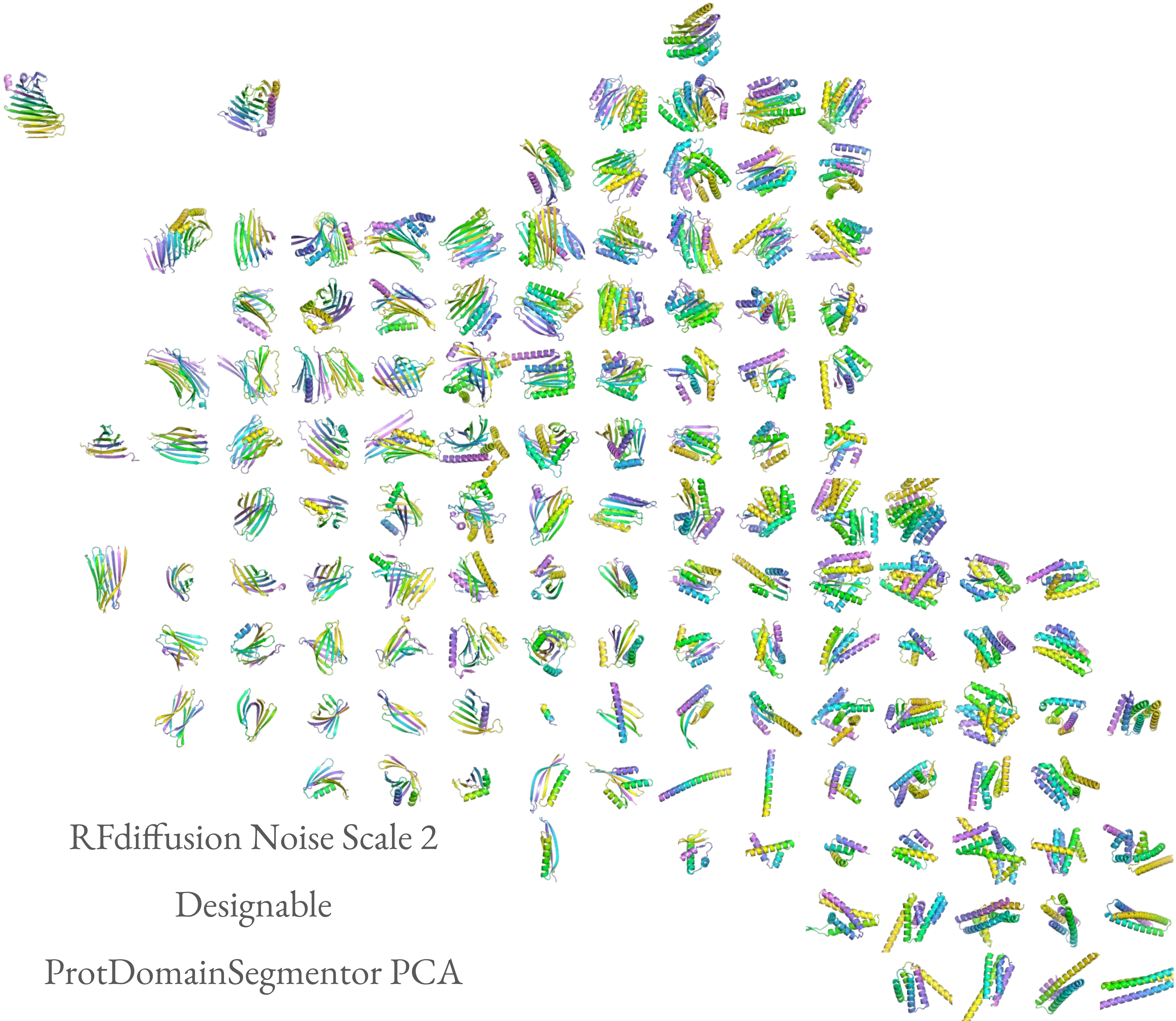

RFdiffusion Noise Scale 2

Designable

ProtDomainSegmentor PCA

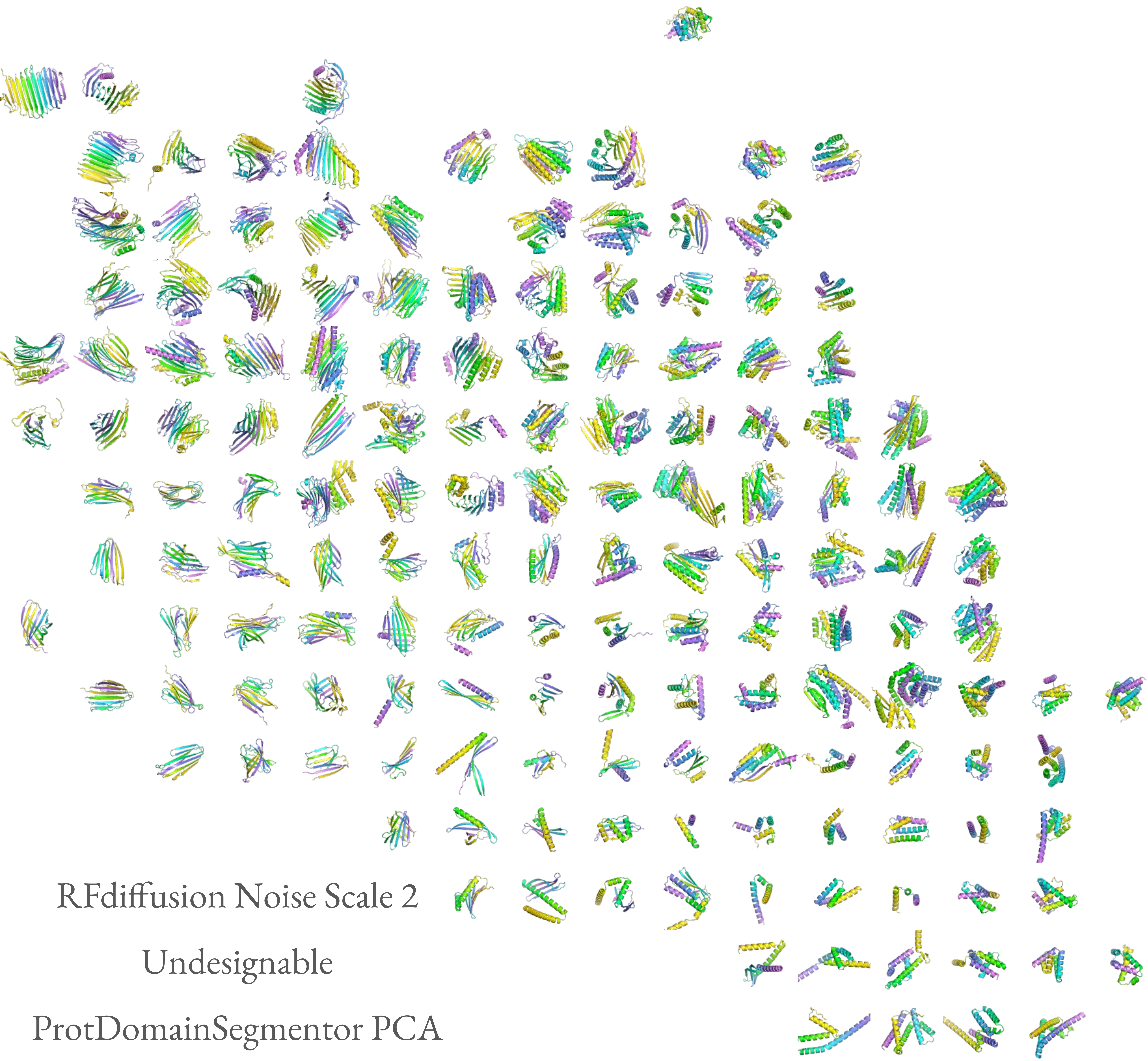

RFdiffusion Noise Scale 2
Undesignable
ProtDomainSegmentor PCA

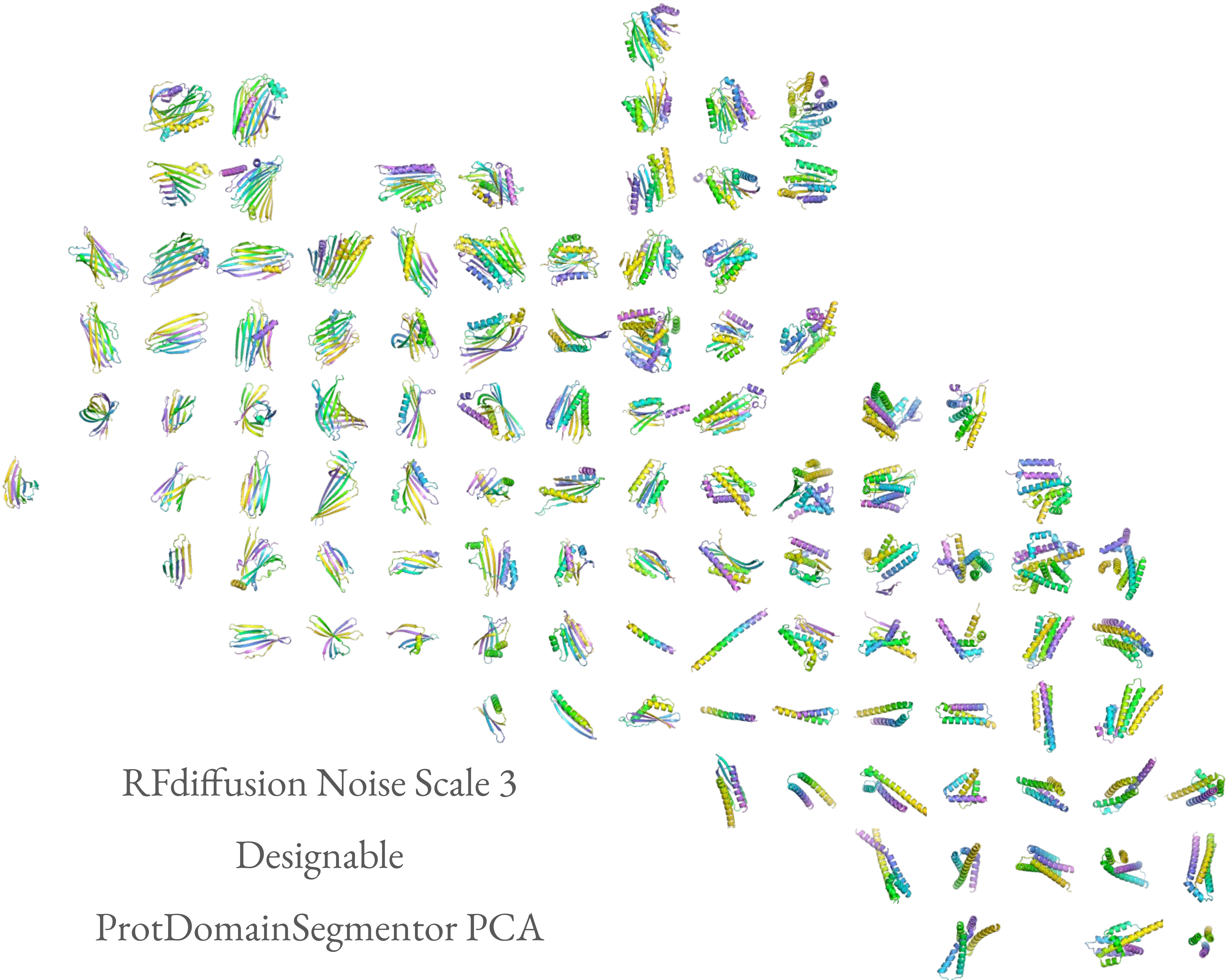

RFdiffusion Noise Scale 3

Designable

ProtDomainSegmentor PCA

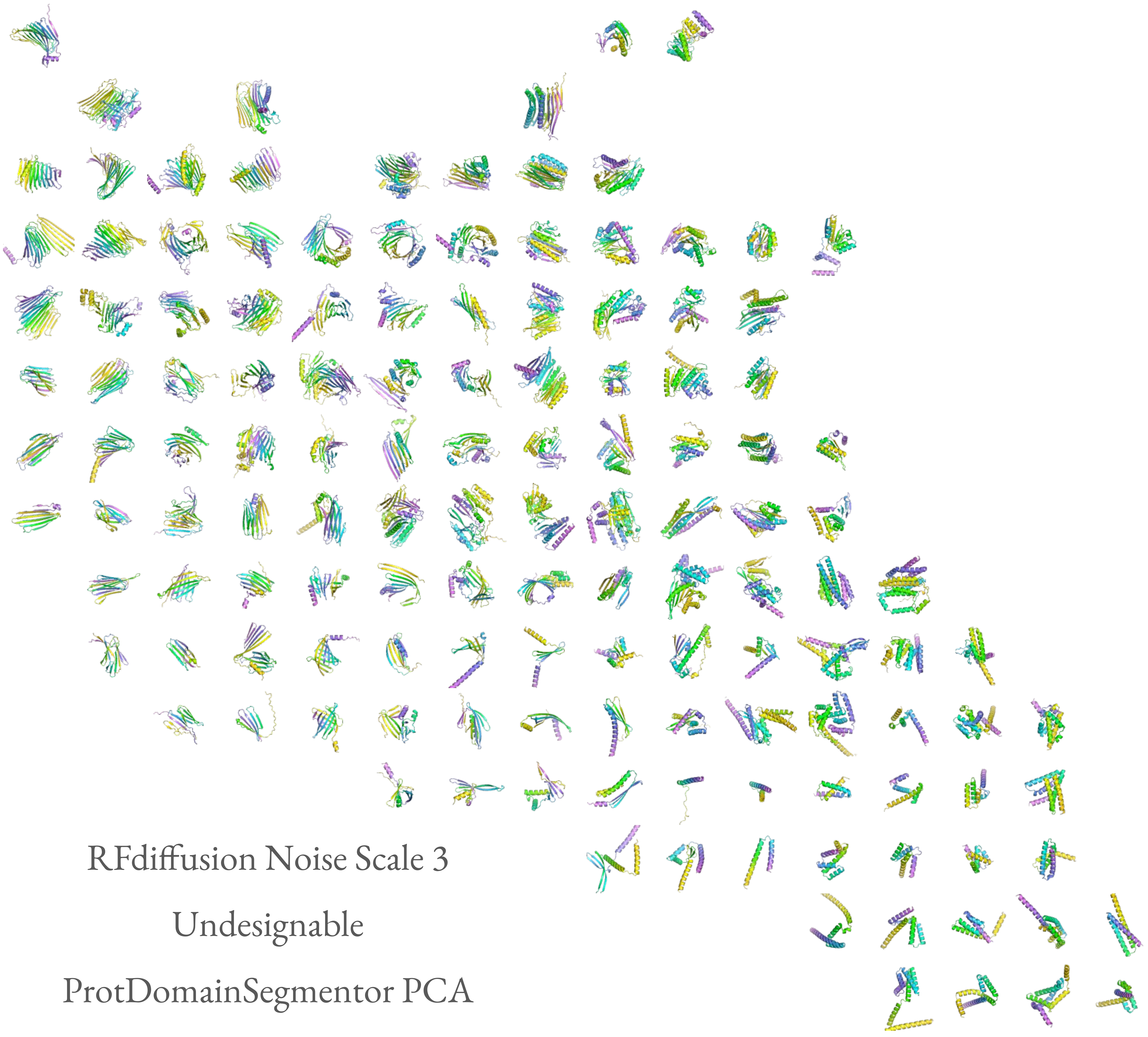
RFdiffusion Noise Scale 3
Undesignable
ProtDomainSegmentor PCA

MultiFlow Designable

ProtDomainSegmentor PCA

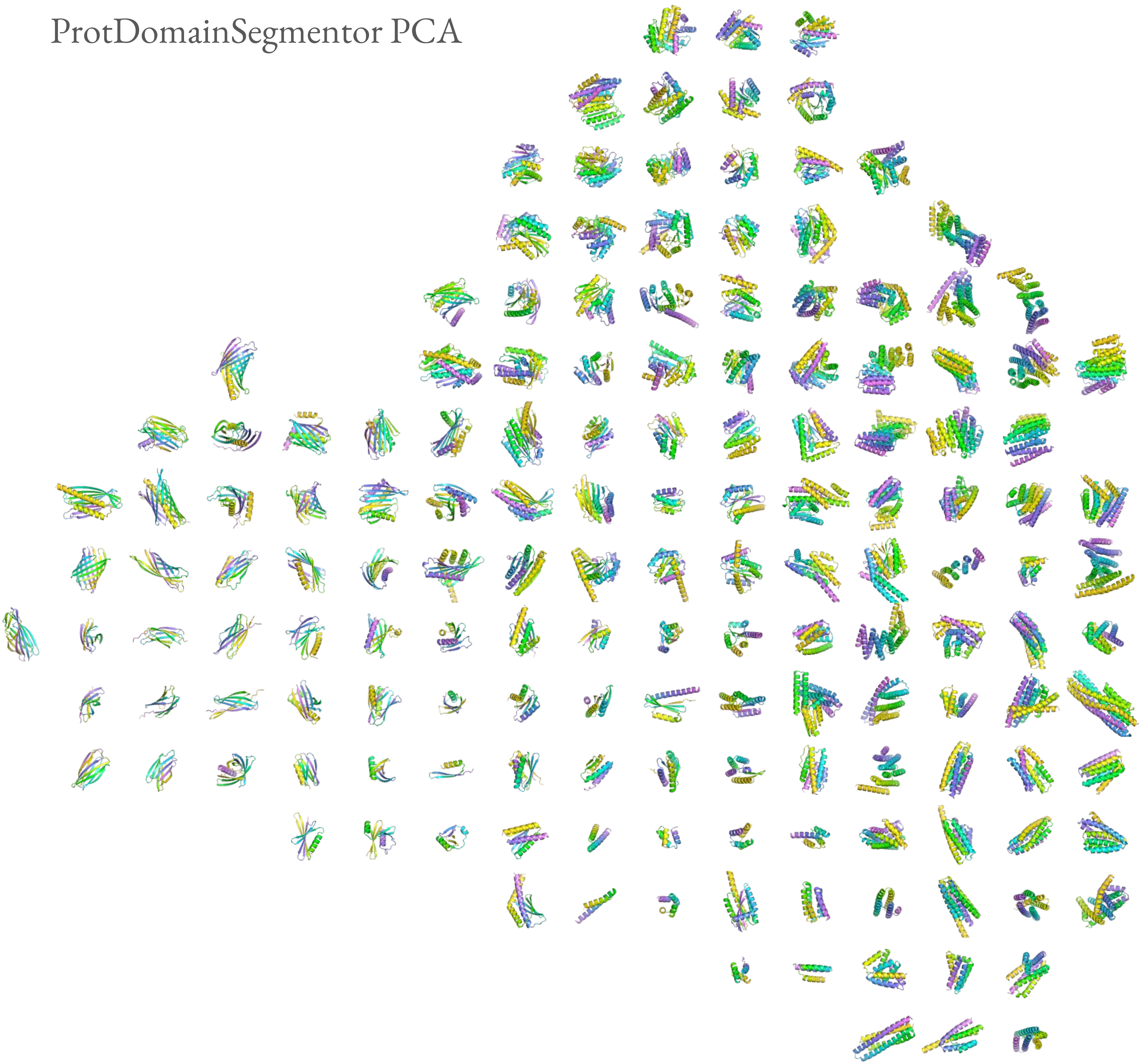

MultiFlow Undesignable

ProtDomainSegmentor PCA

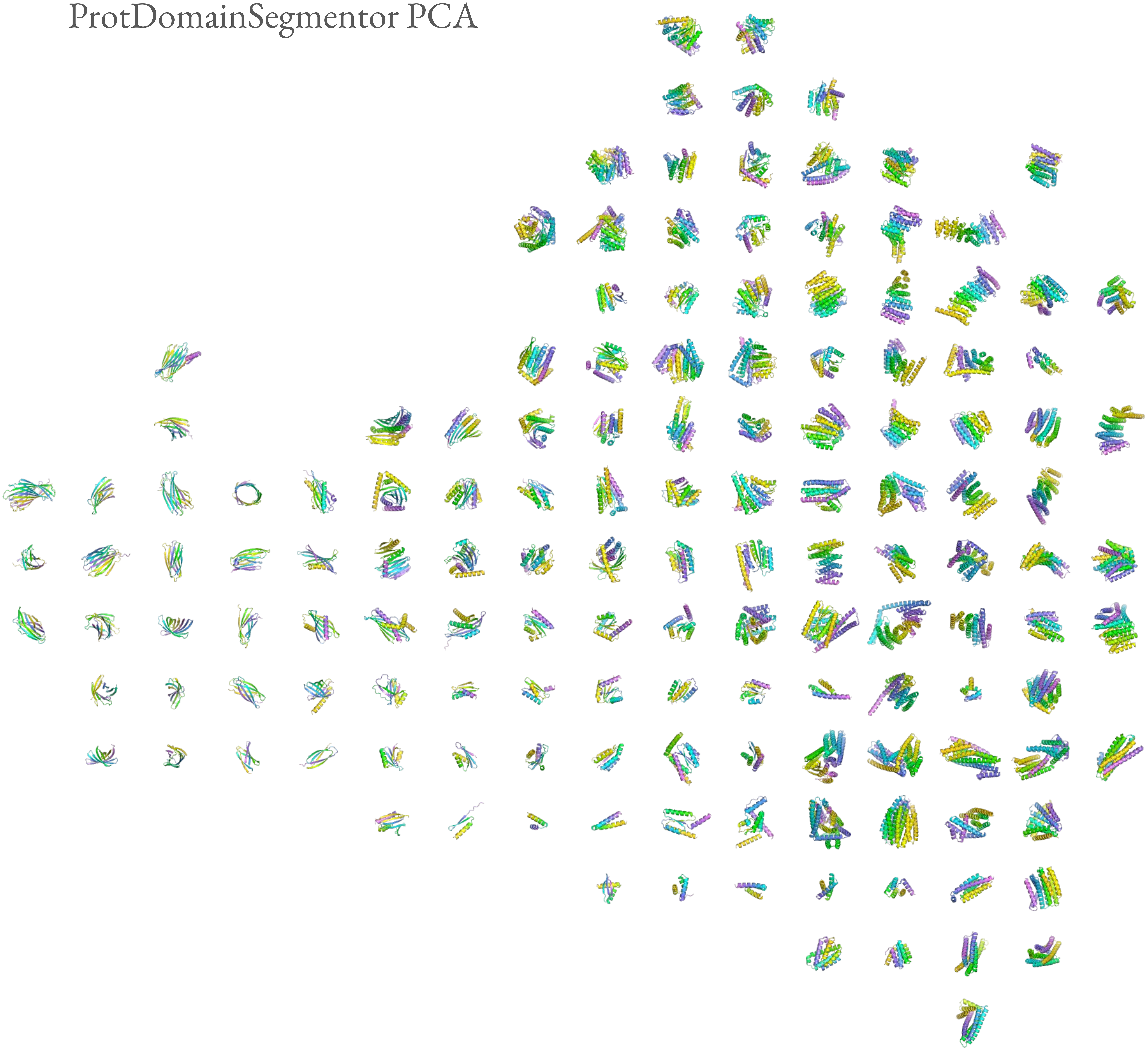

Chroma Designable

ProtDomainSegmentor PCA

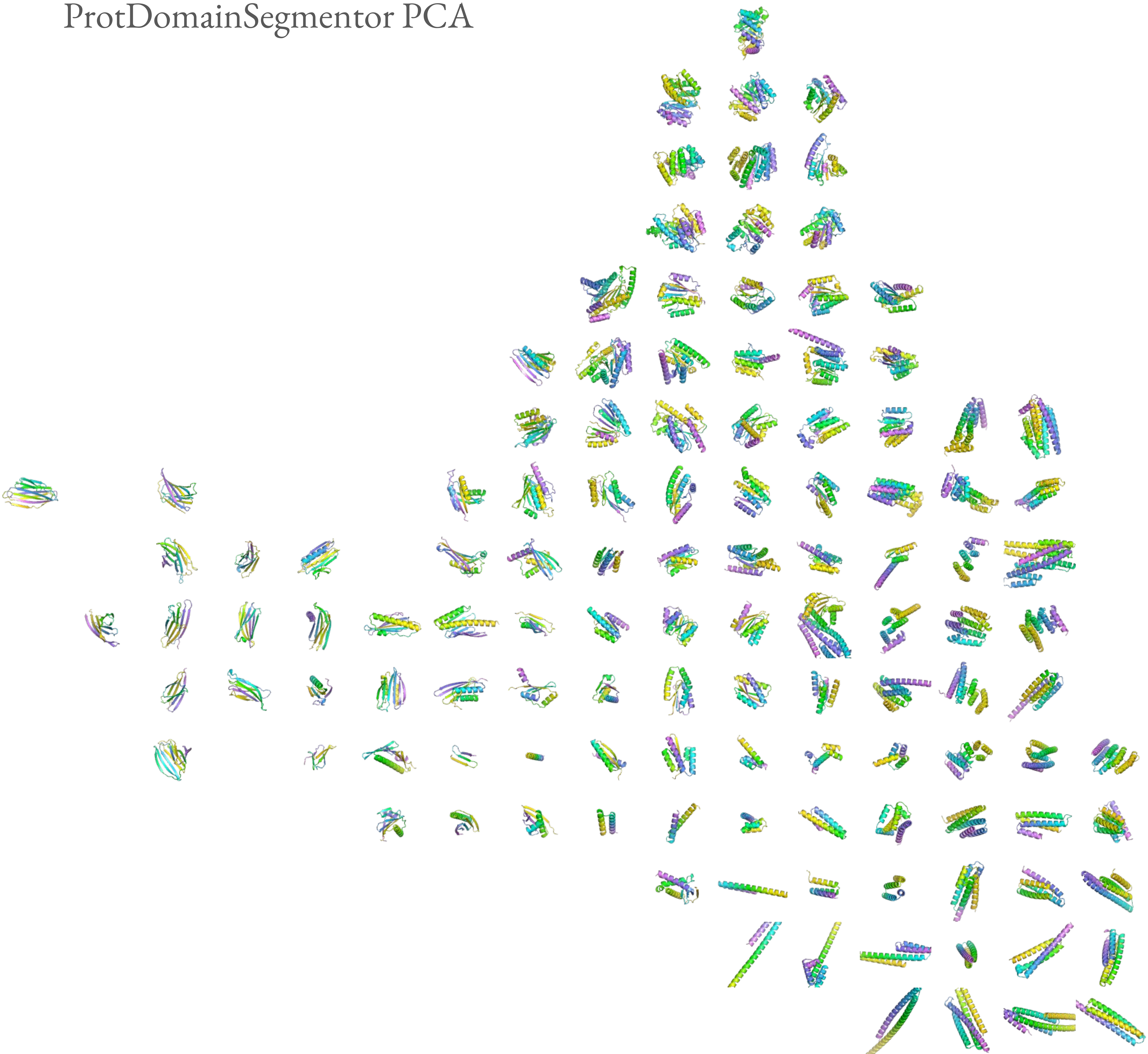

Chroma Undesignable

ProtDomainSegmentor PCA

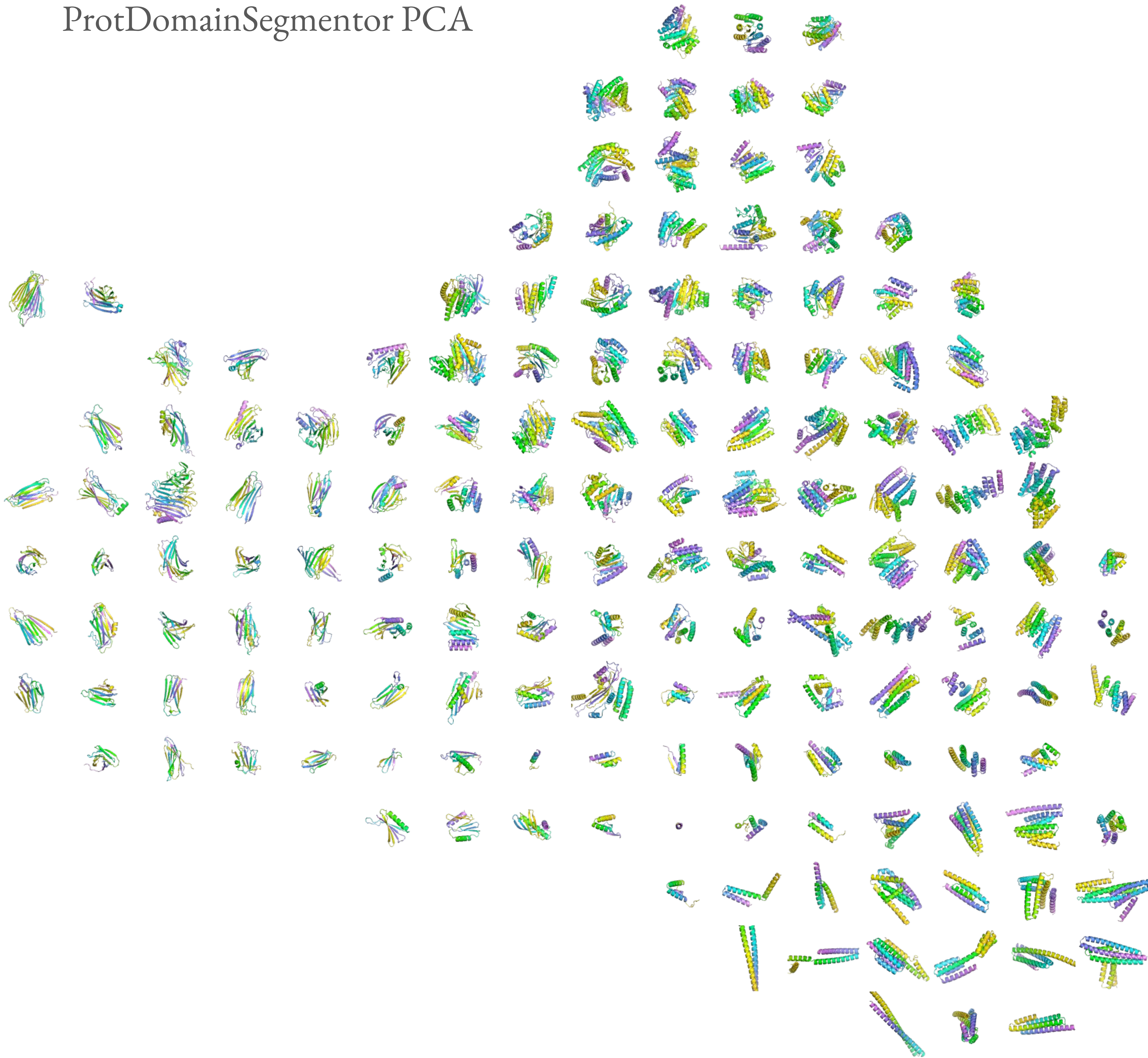

Chroma Inverse Temperature 4

Designable

ProtDomainSegmentor PCA

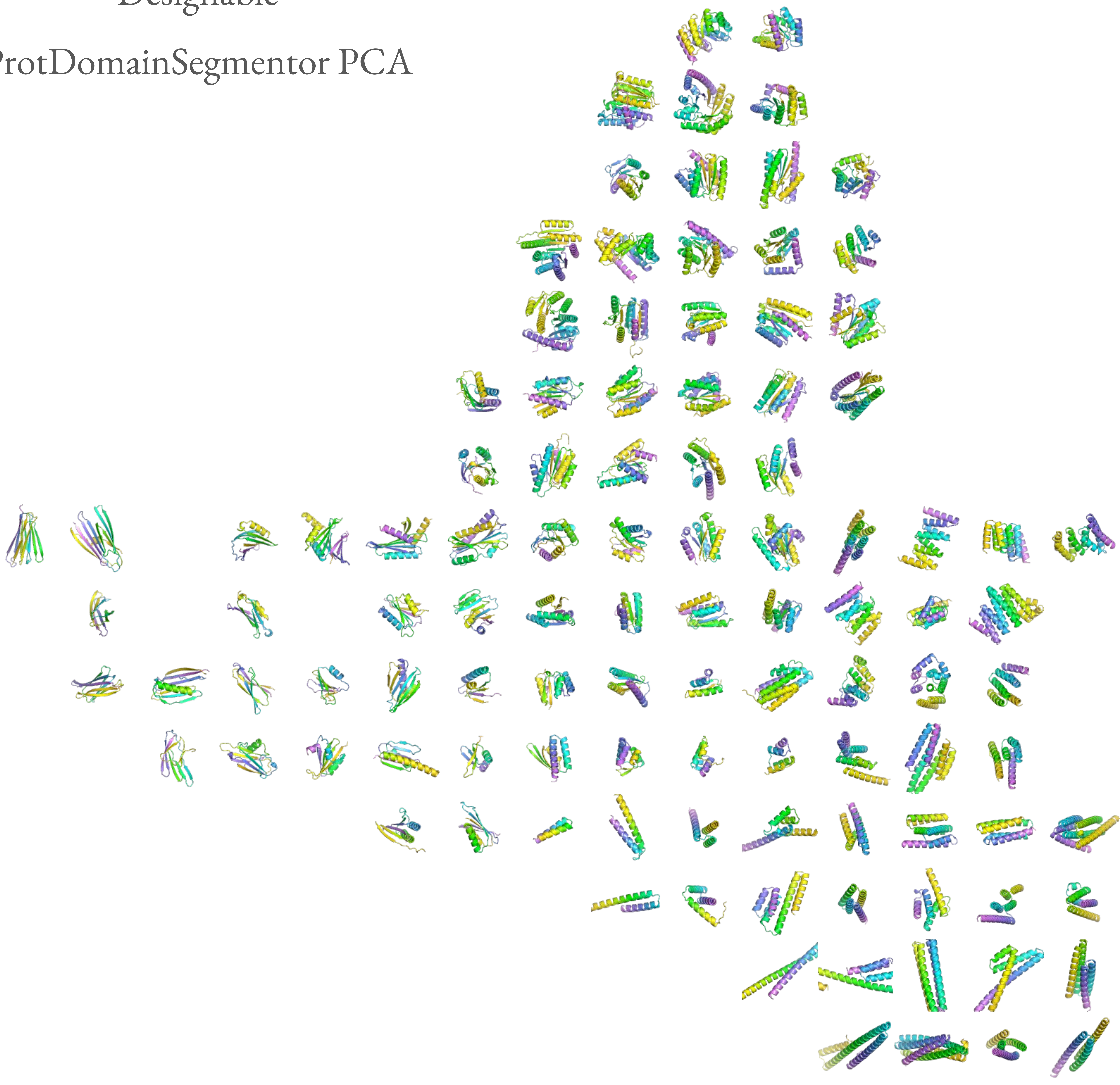

Chroma Inverse Temperature 4

Undesignable

ProtDomainSegmentor PCA

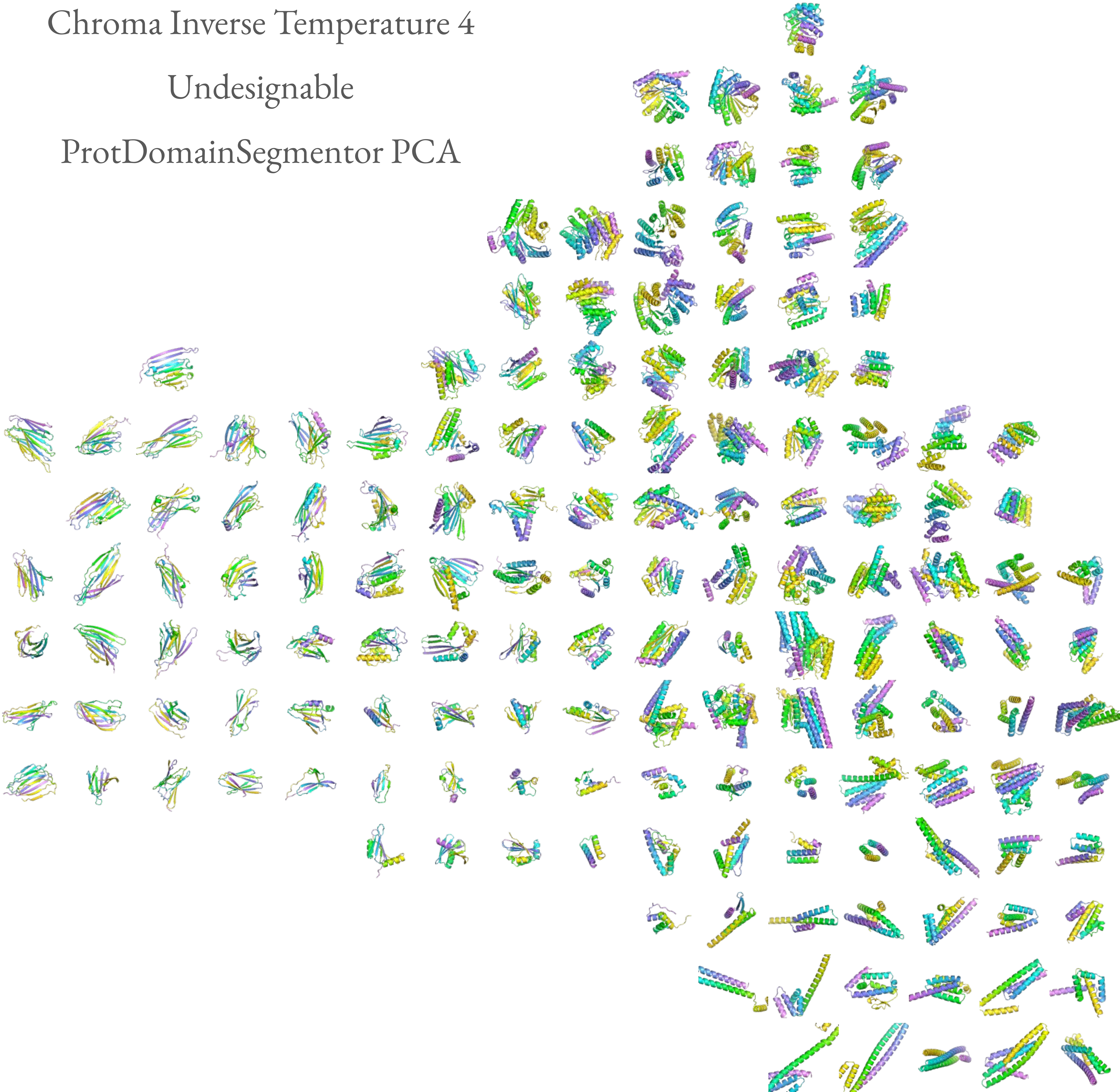

Chroma Inverse Temperature 3

Designable

ProtDomainSegmentor PCA

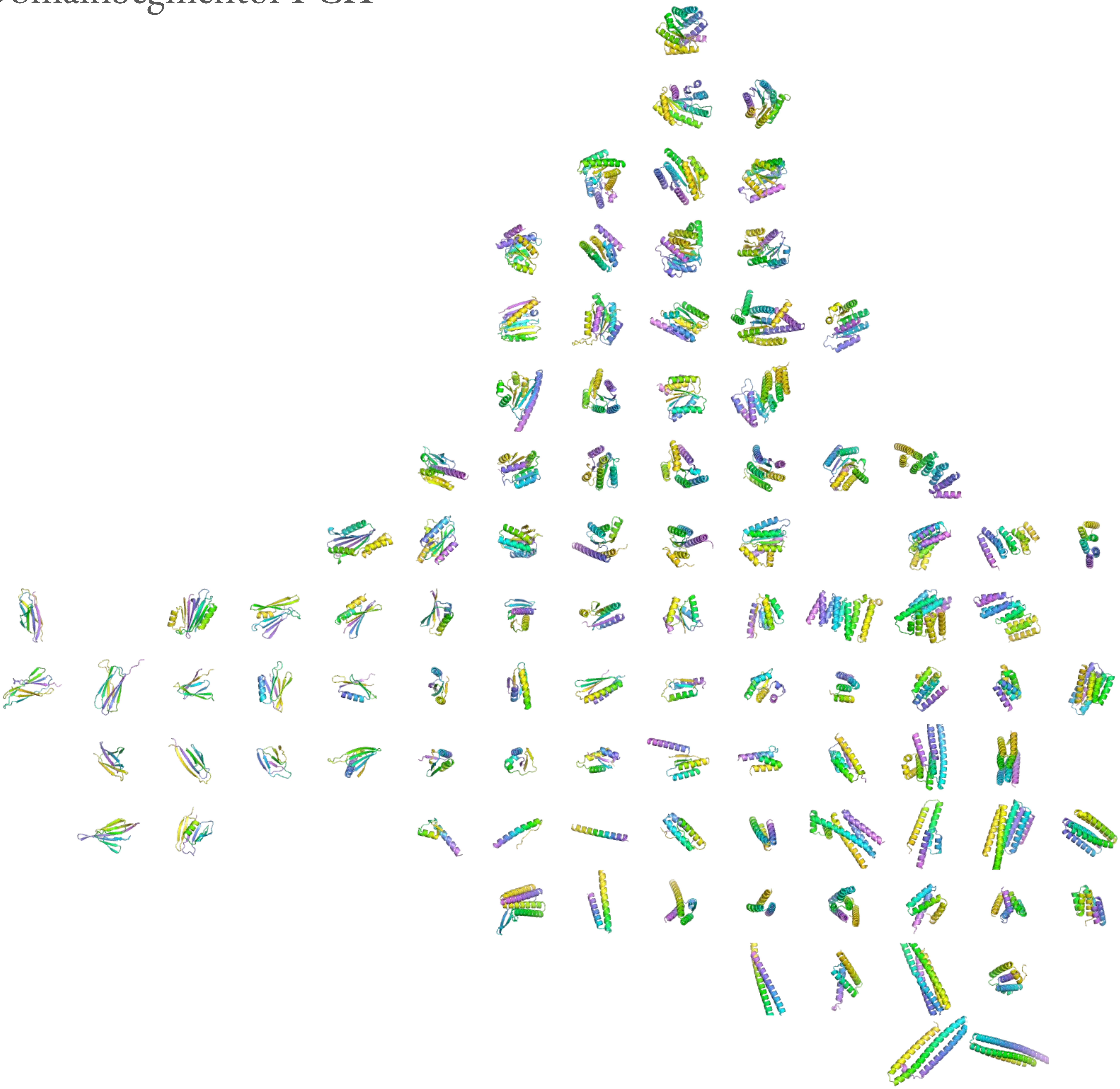

Chroma Inverse Temperature 3

Undesignable

ProtDomainSegmentor PCA

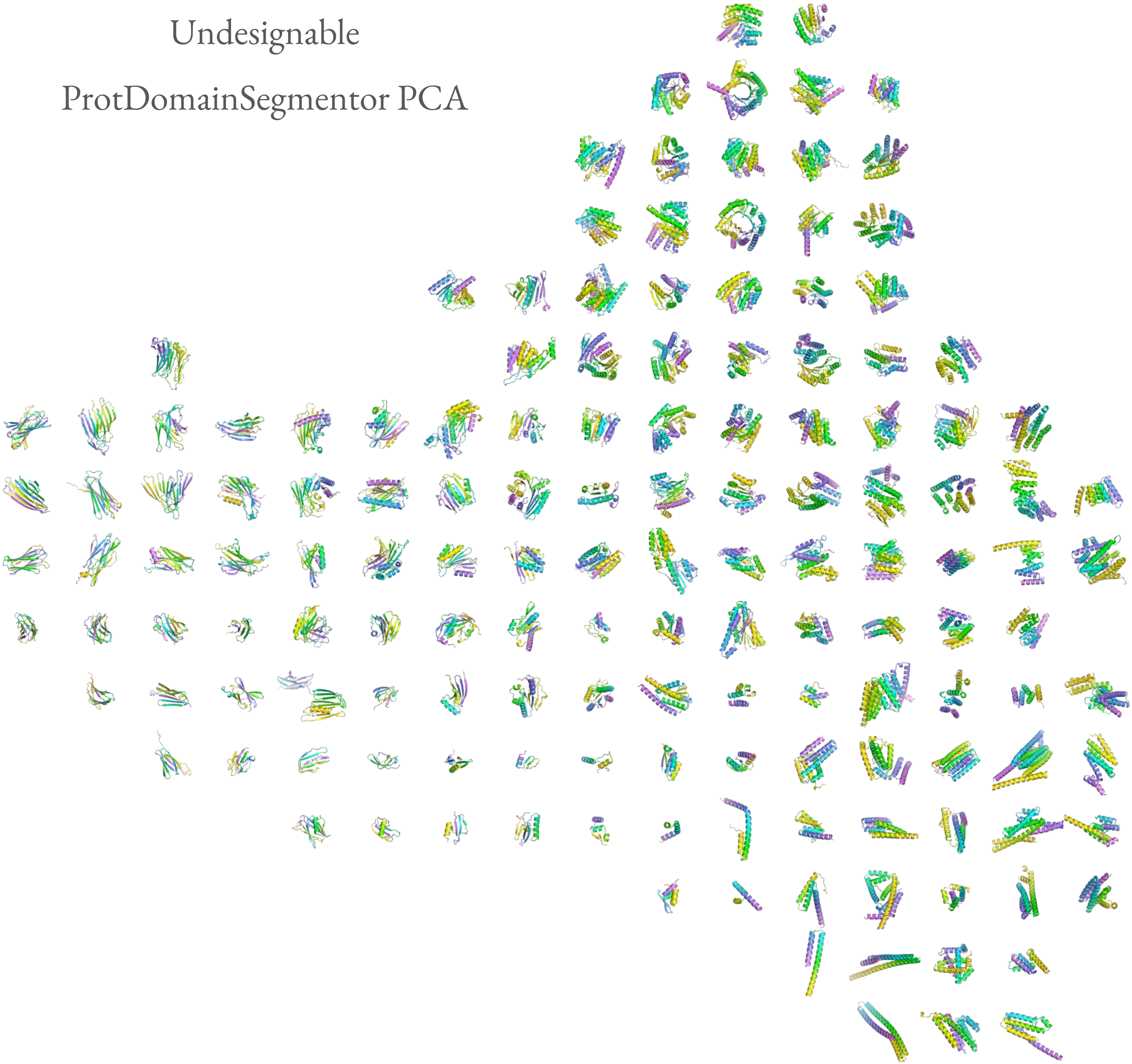

Genie2 Designable

ProtDomainSegmentor PCA

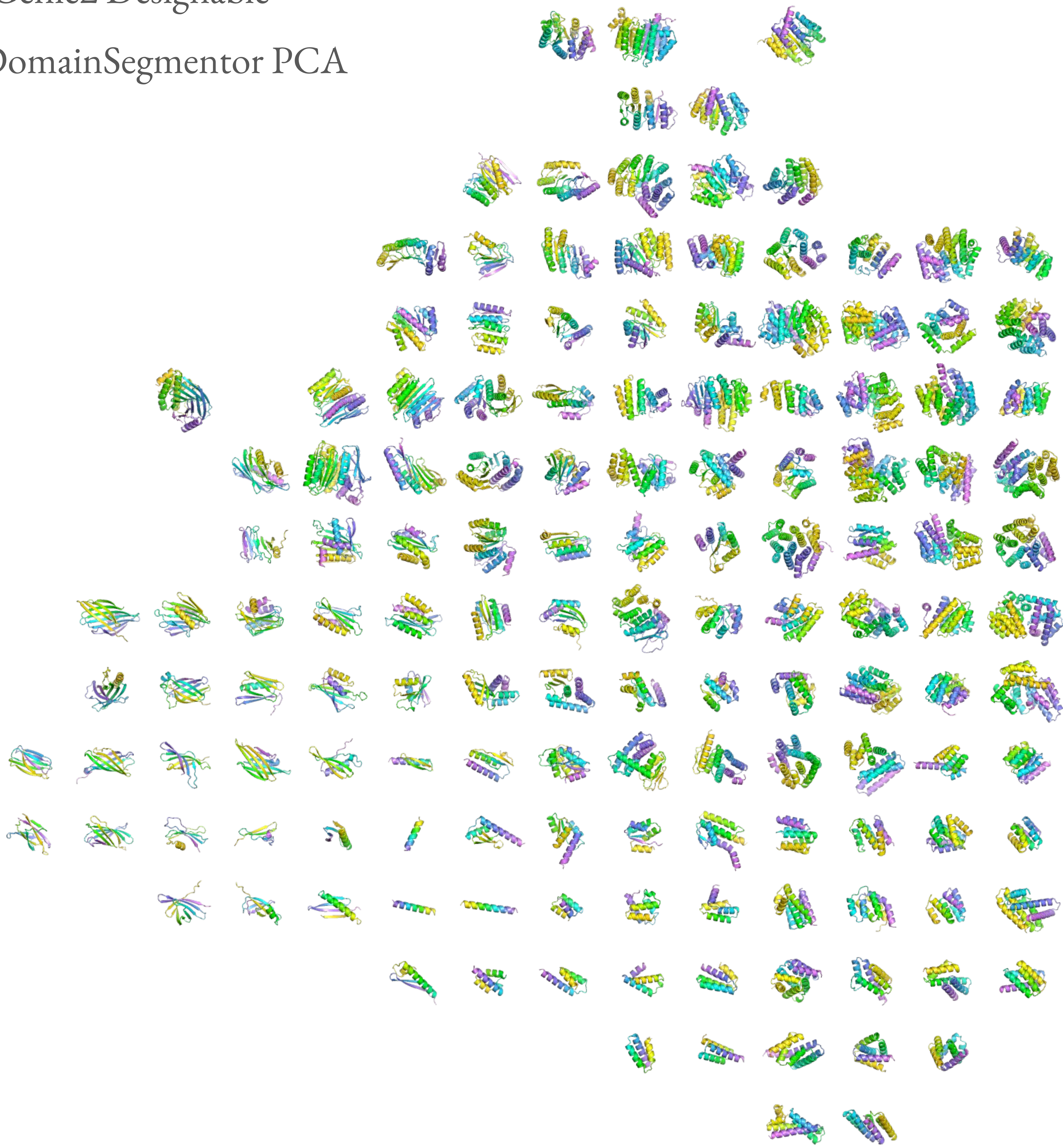

Genie2 Undesignable

ProtDomainSegmentor PCA

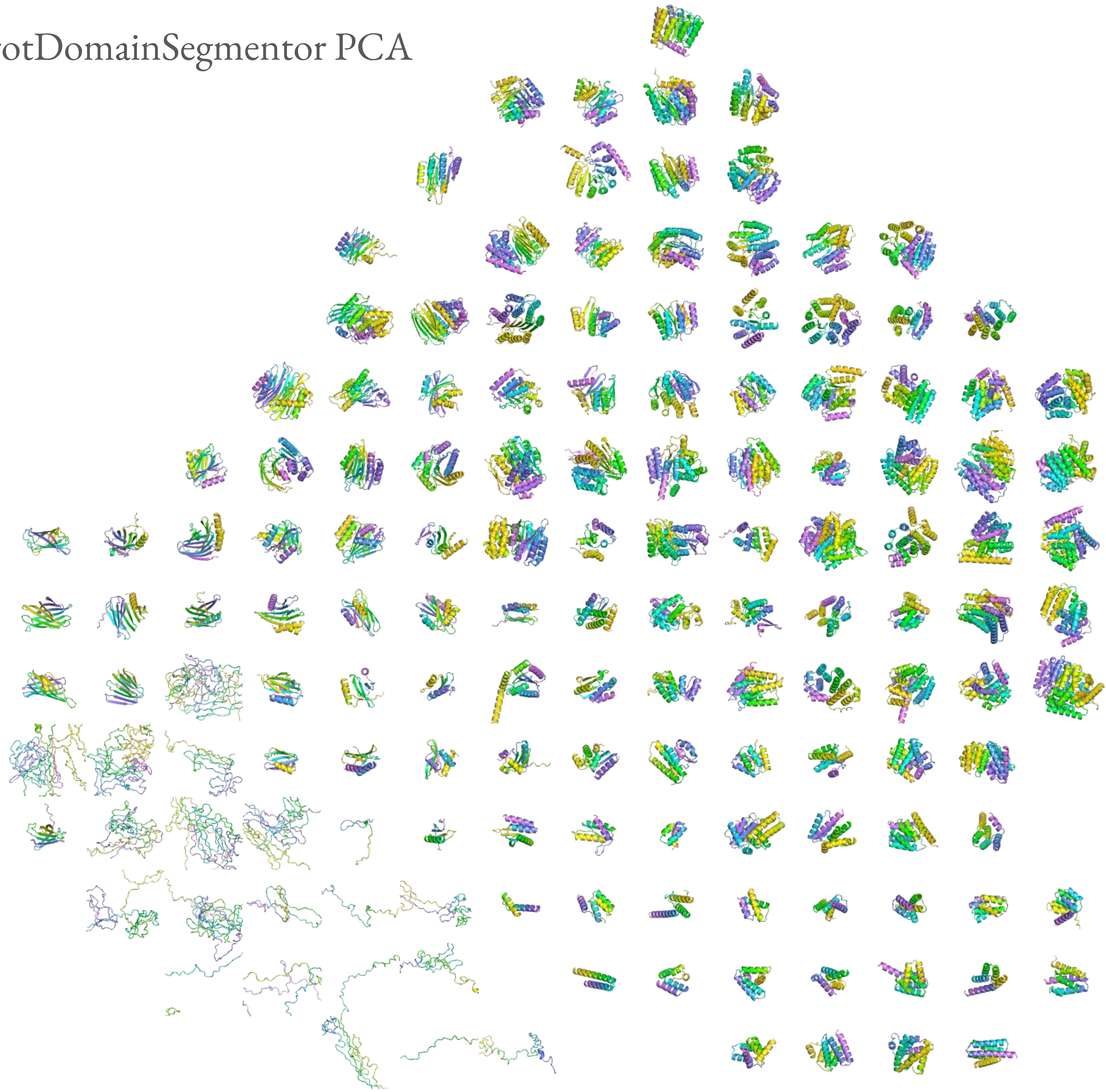

Genie2 Scale 0.8 Designable

ProtDomainSegmentor PCA

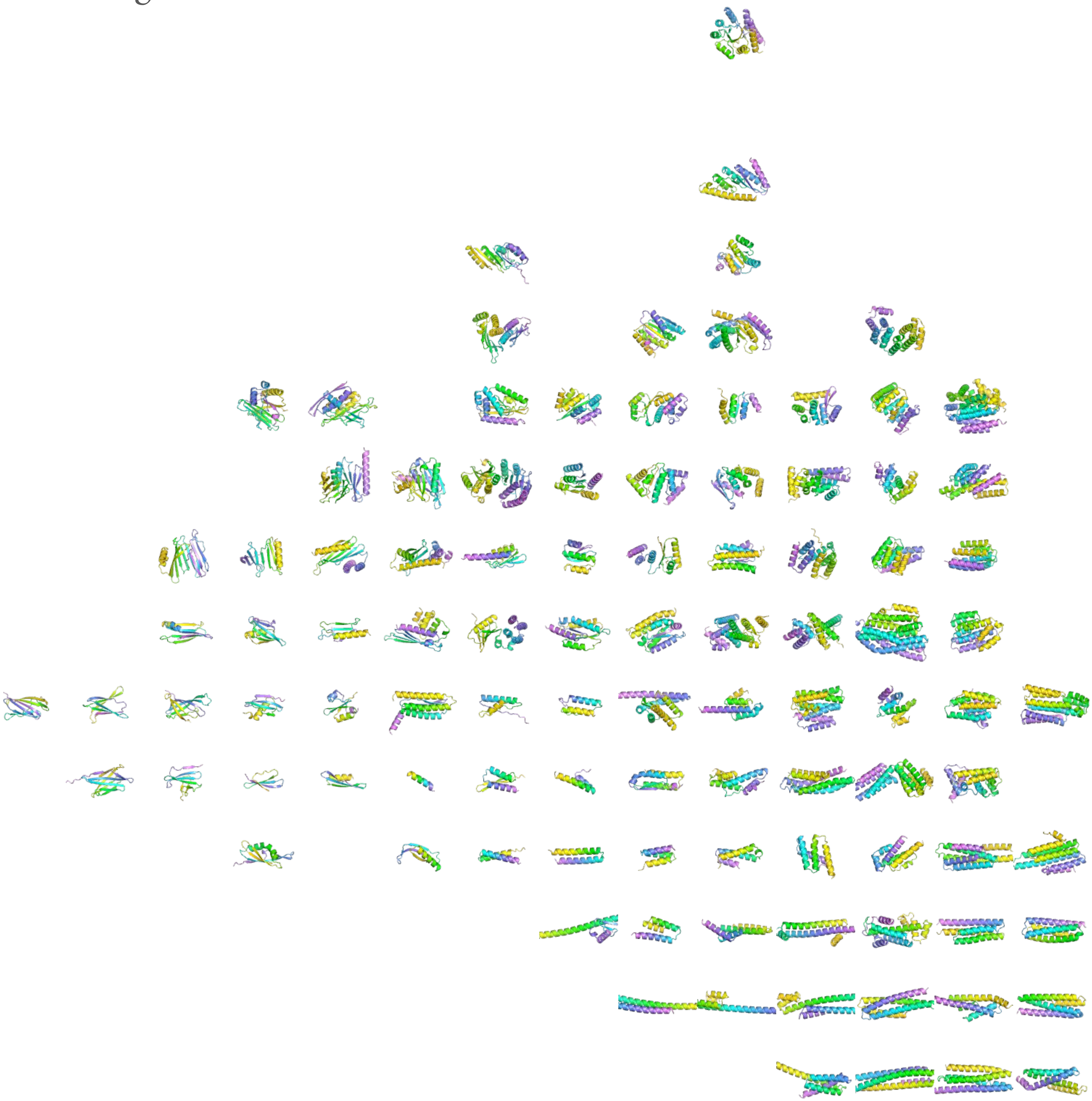

Genie2 Scale 0.8 Undesignable

ProtDomainSegmentor PCA

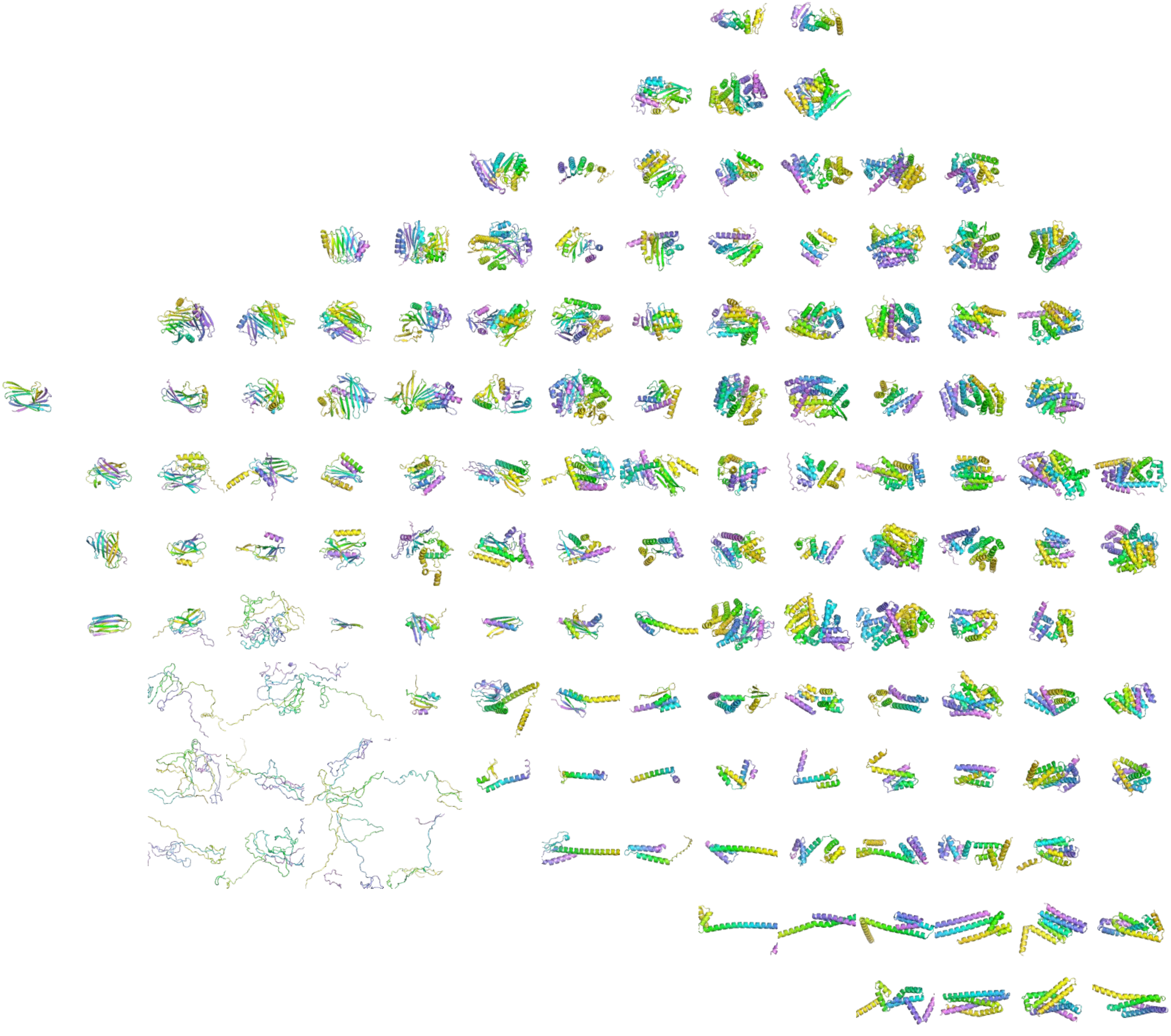

Genie2 Scale 1.0 Designable

ProtDomainSegmentor PCA

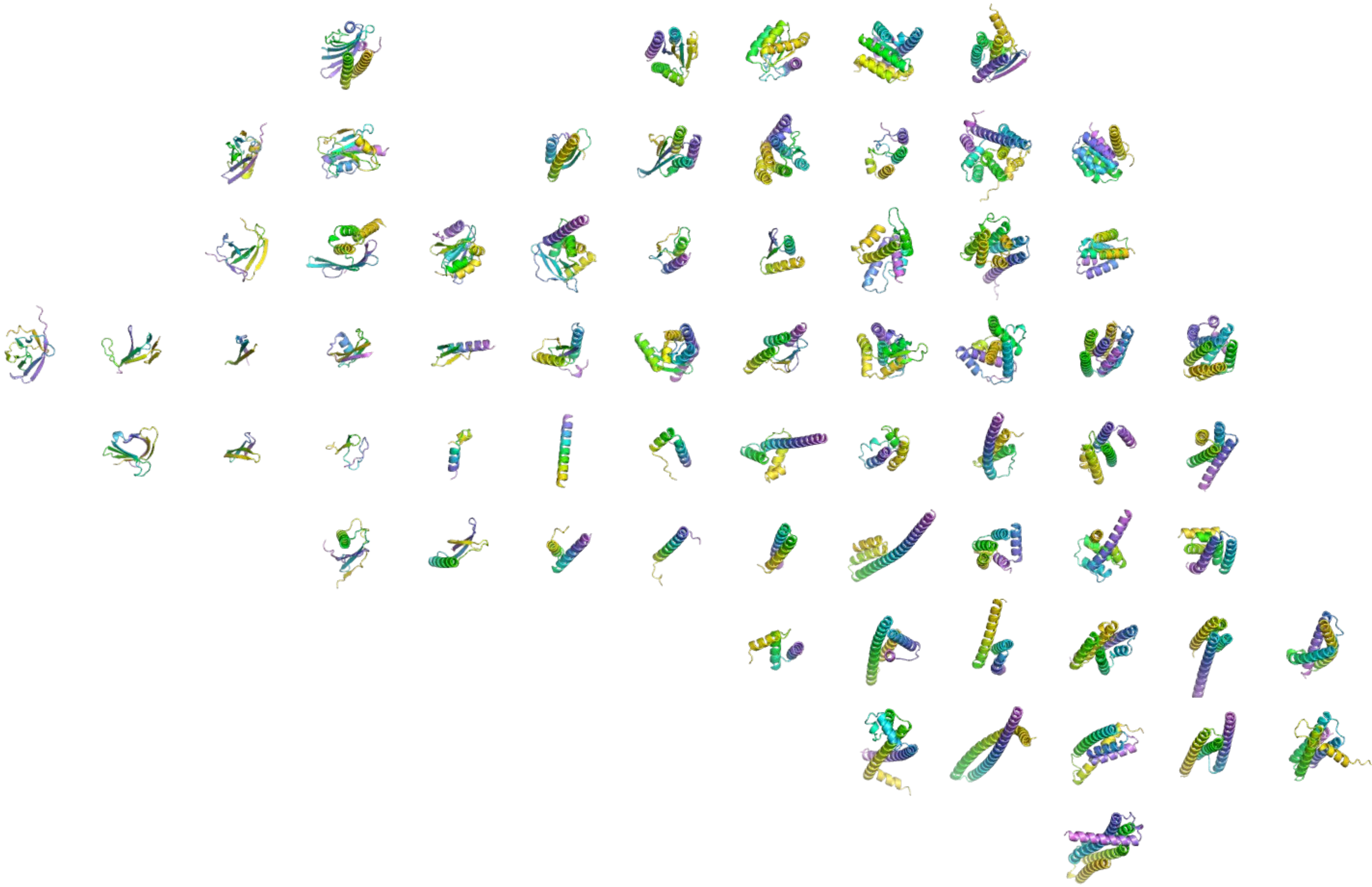

Genie2 Scale 1.0 Undesignable

ProtDomainSegmentor PCA

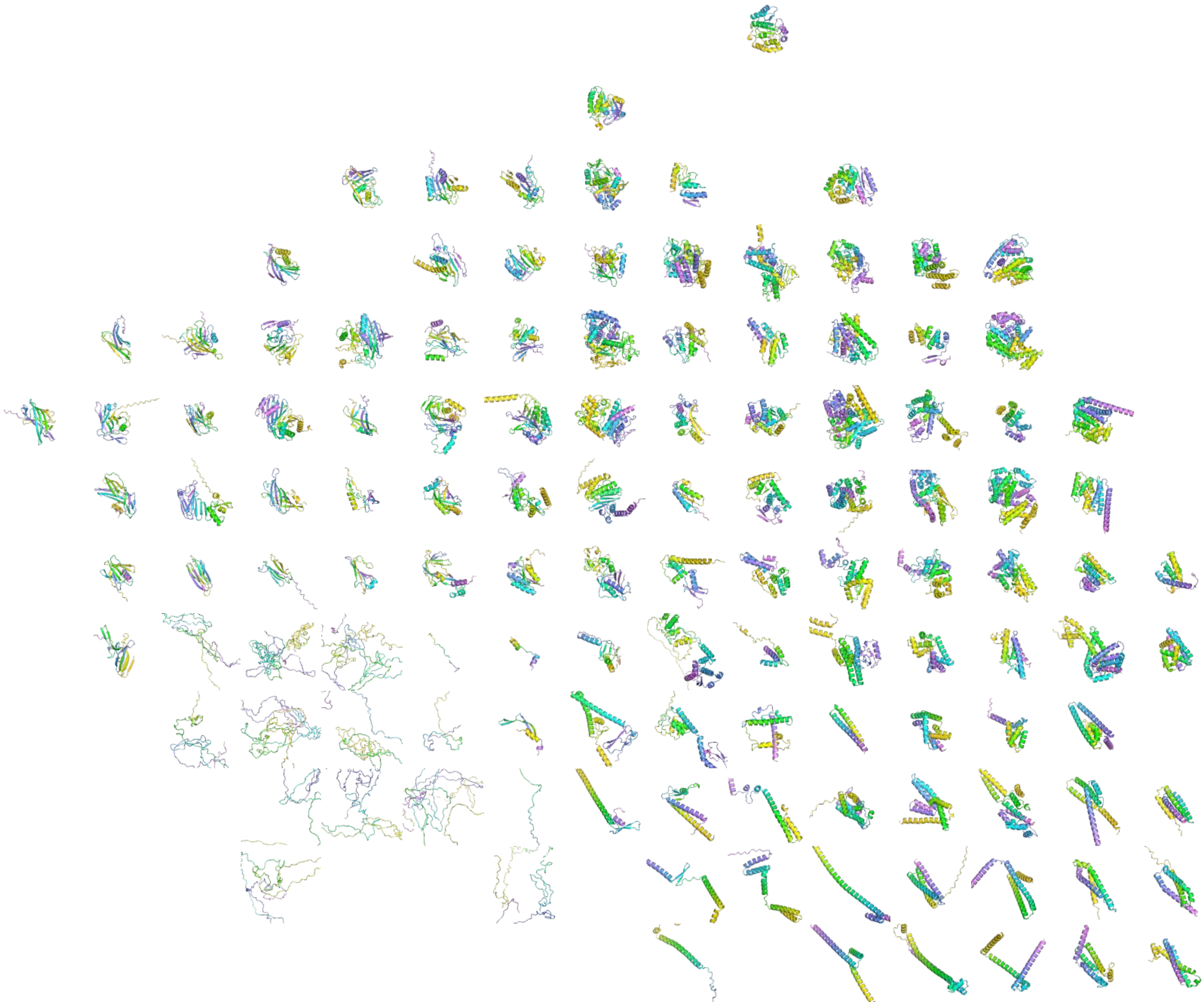

Protpardelle Stepscale 1.2 Designable

ProtDomainSegmentor PCA

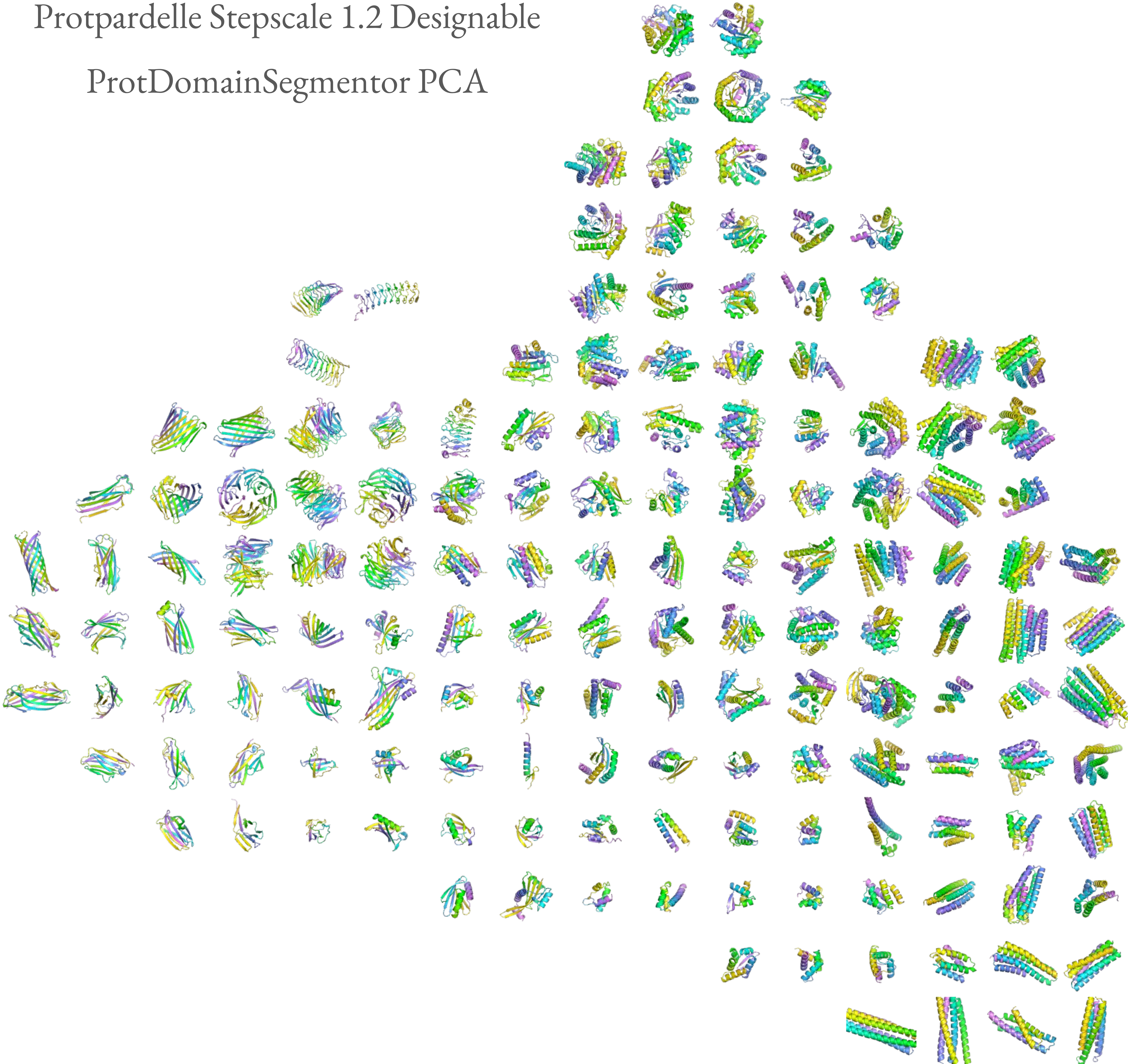

Protpardelle Stepscale 1.2 Undesignable

ProtDomainSegmentor PCA

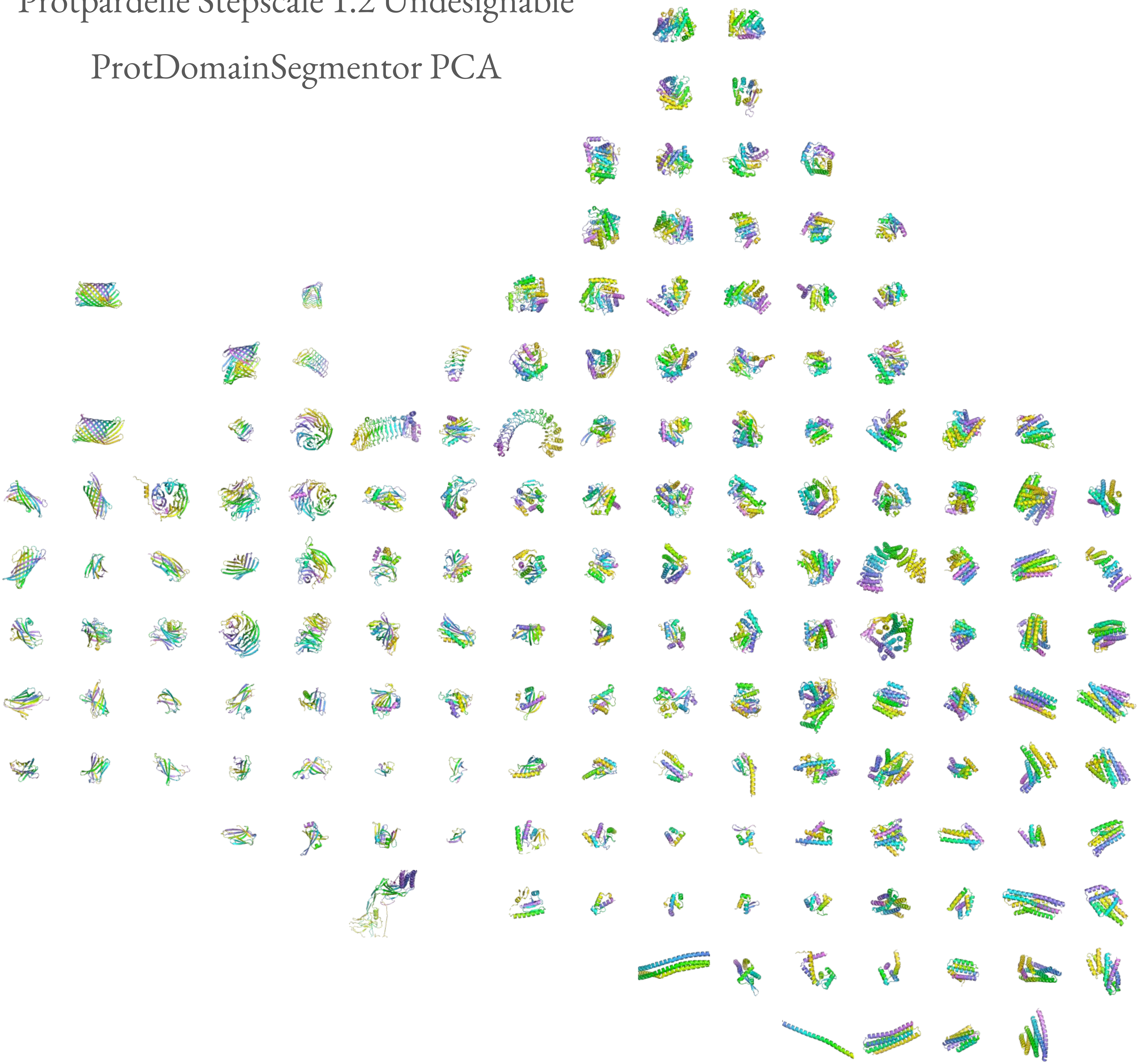

Protpardelle Stepscale 1.0 Designable

ProtDomainSegmentor PCA

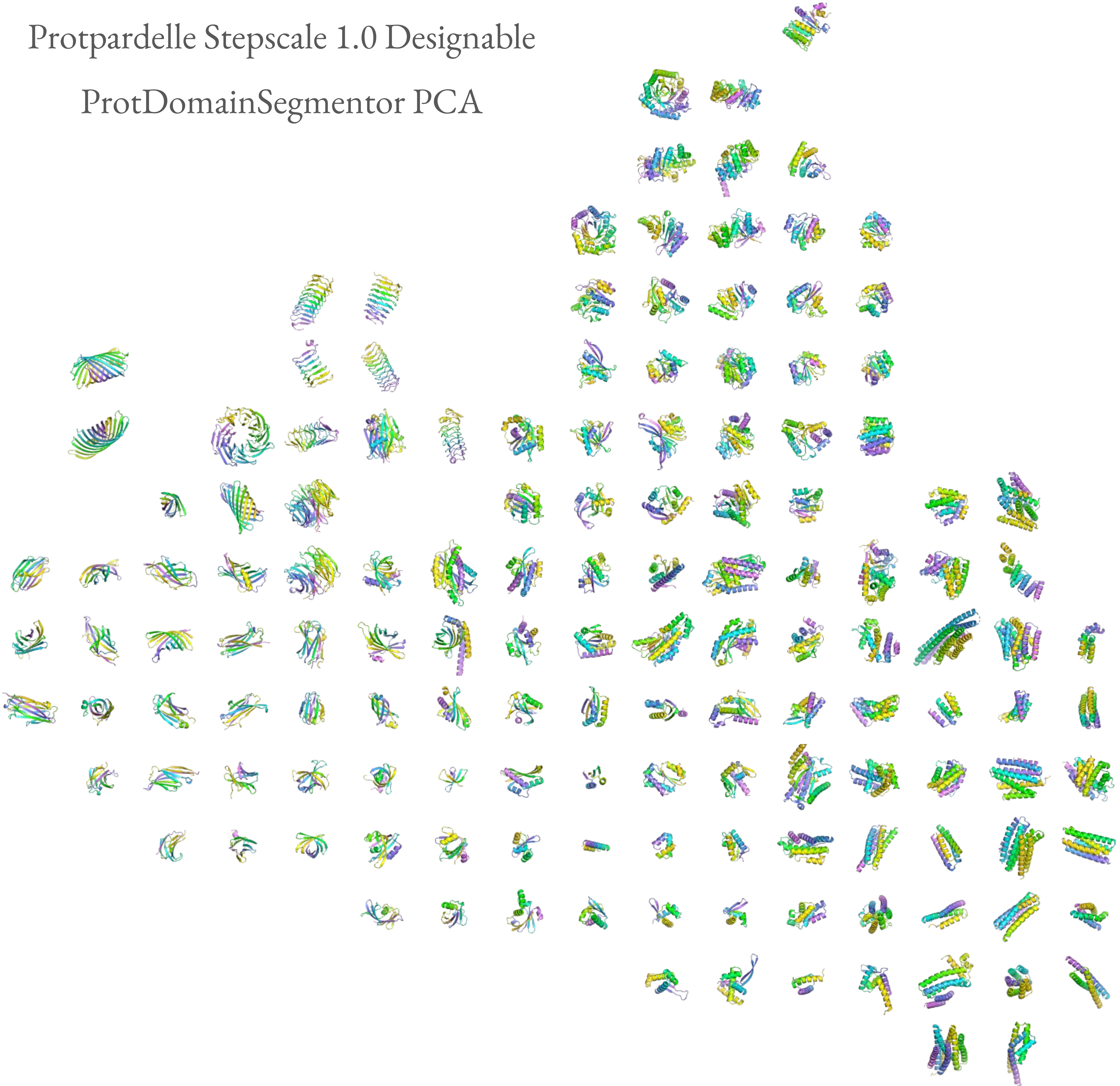

Protpardelle Stepscale 1.0 Undesignable

ProtDomainSegmentor PCA

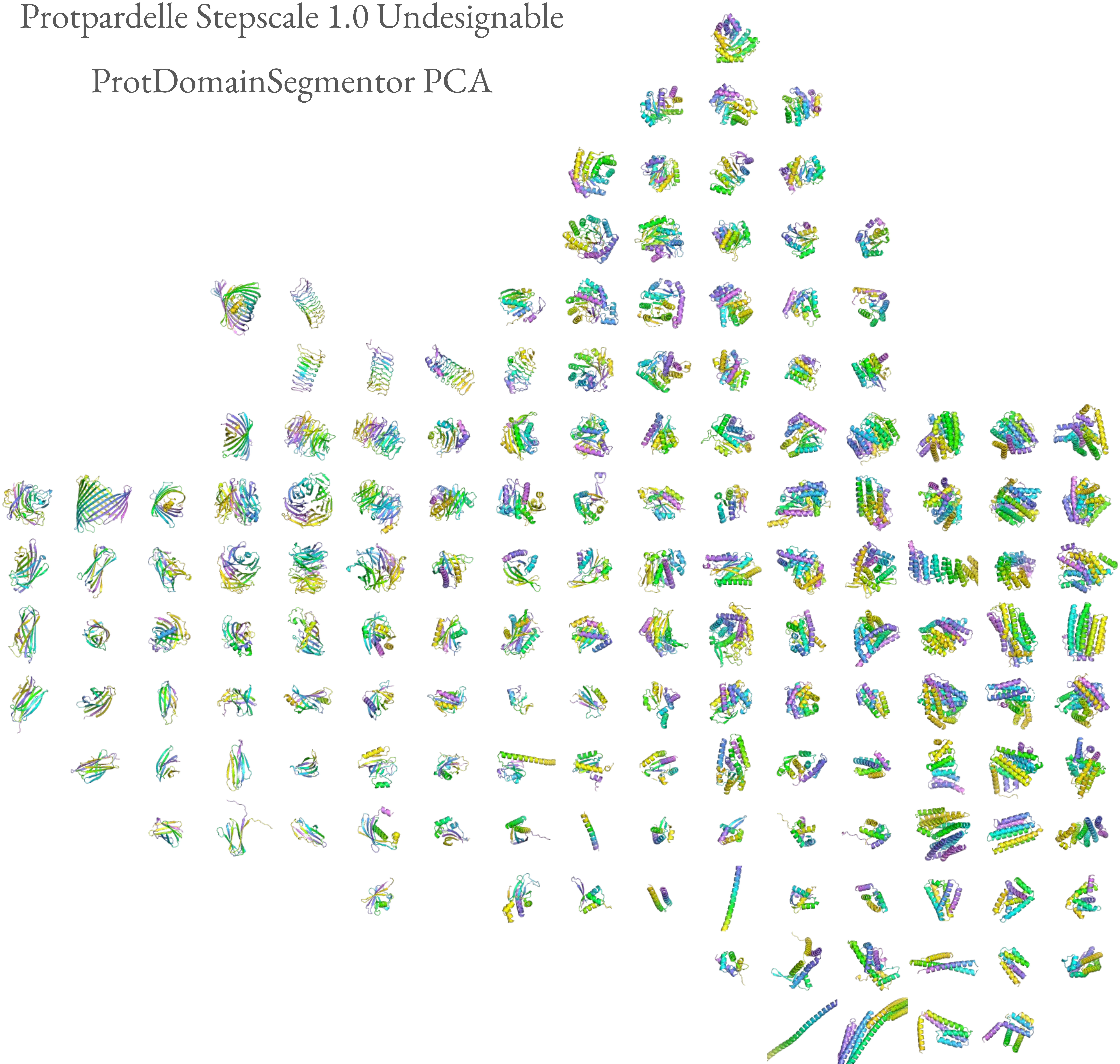

Protpardelle Stepscale 0.8 Designable

ProtDomainSegmentor PCA

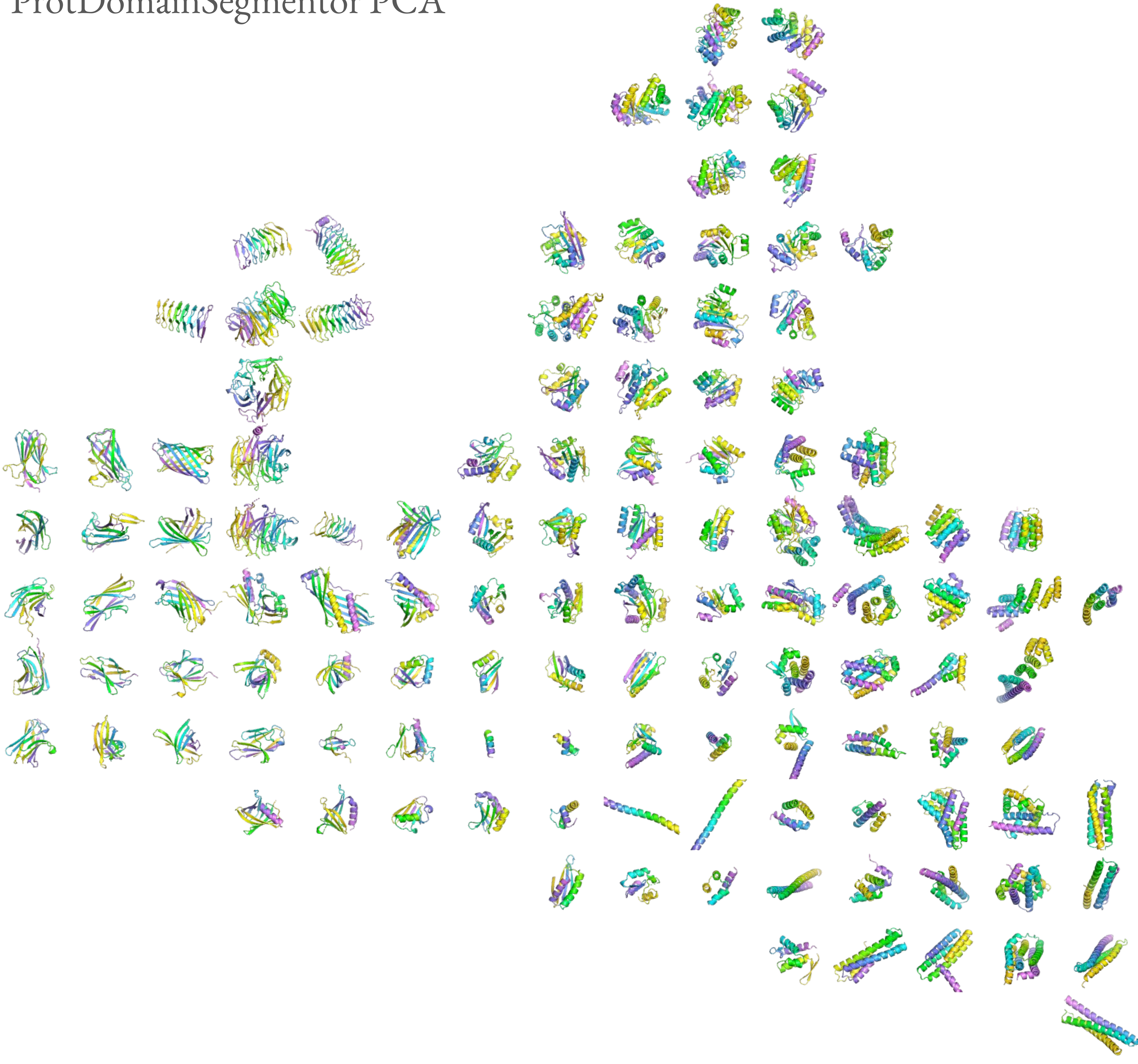

Protpardelle Stepscale 0.8 Undesignable

ProtDomainSegmentor PCA

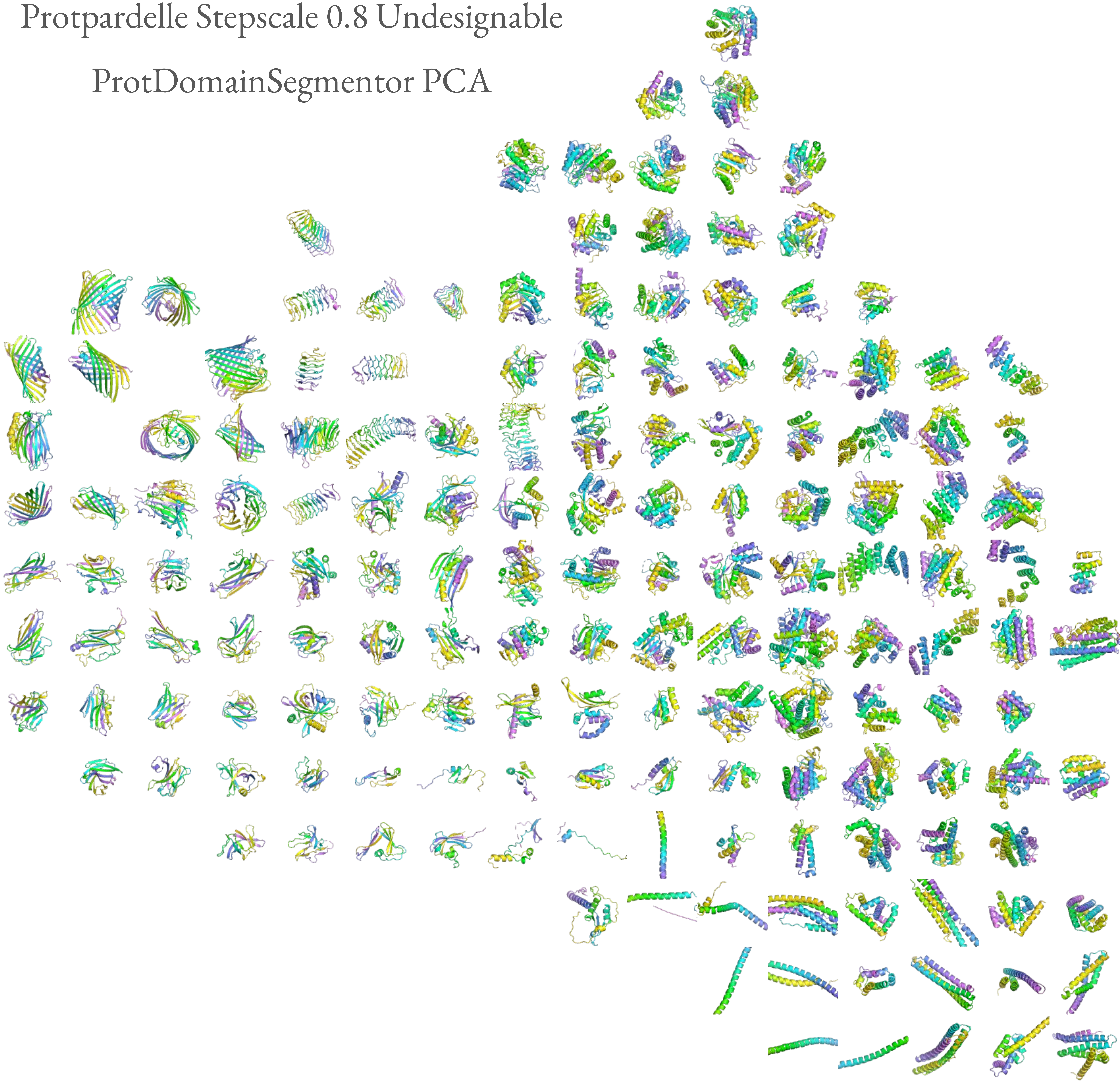

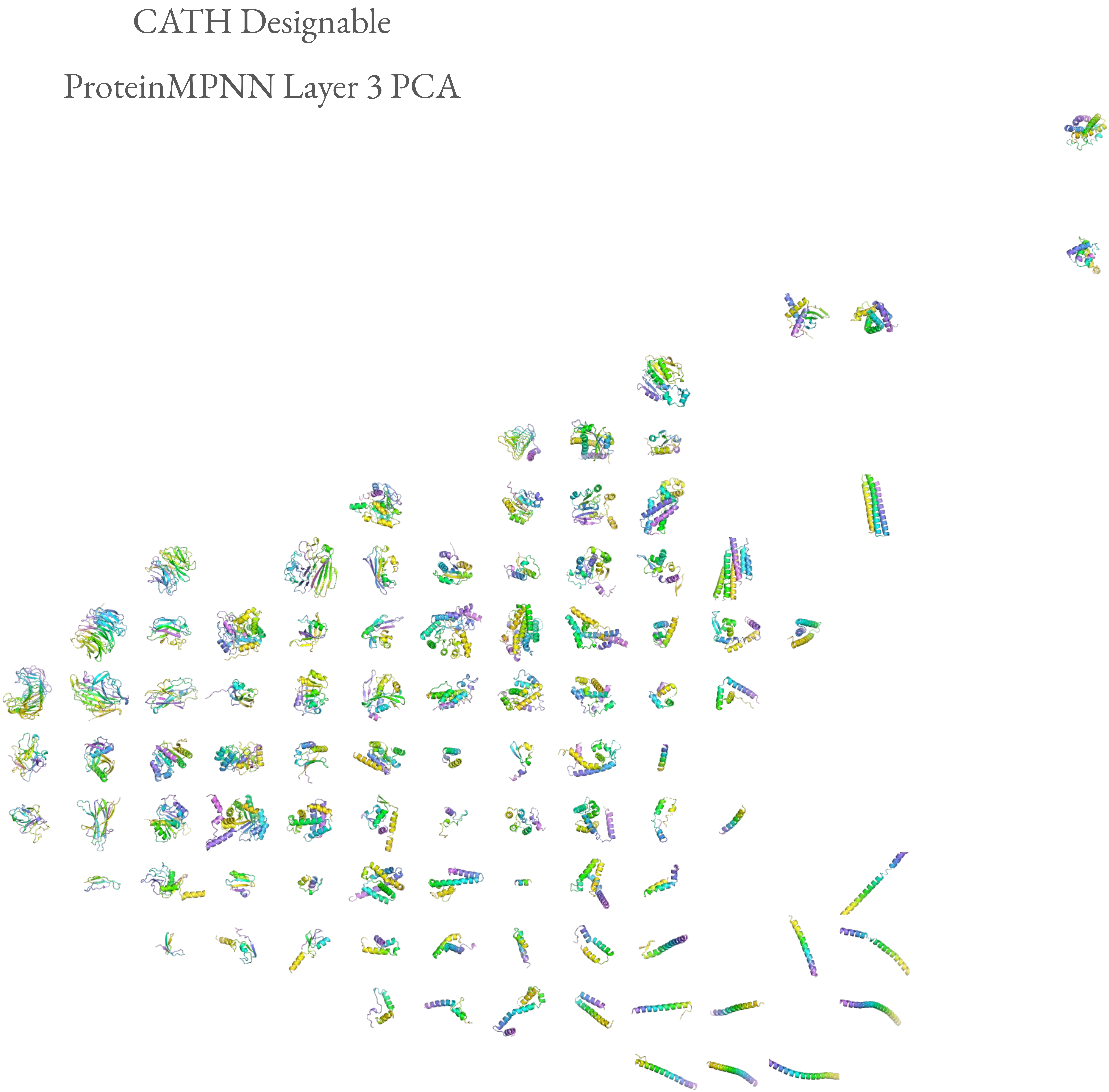
CATH Designable
ProteinMPNN Layer 3 PCA

CATH Undesignable

ProteinMPNN Layer 3 PCA

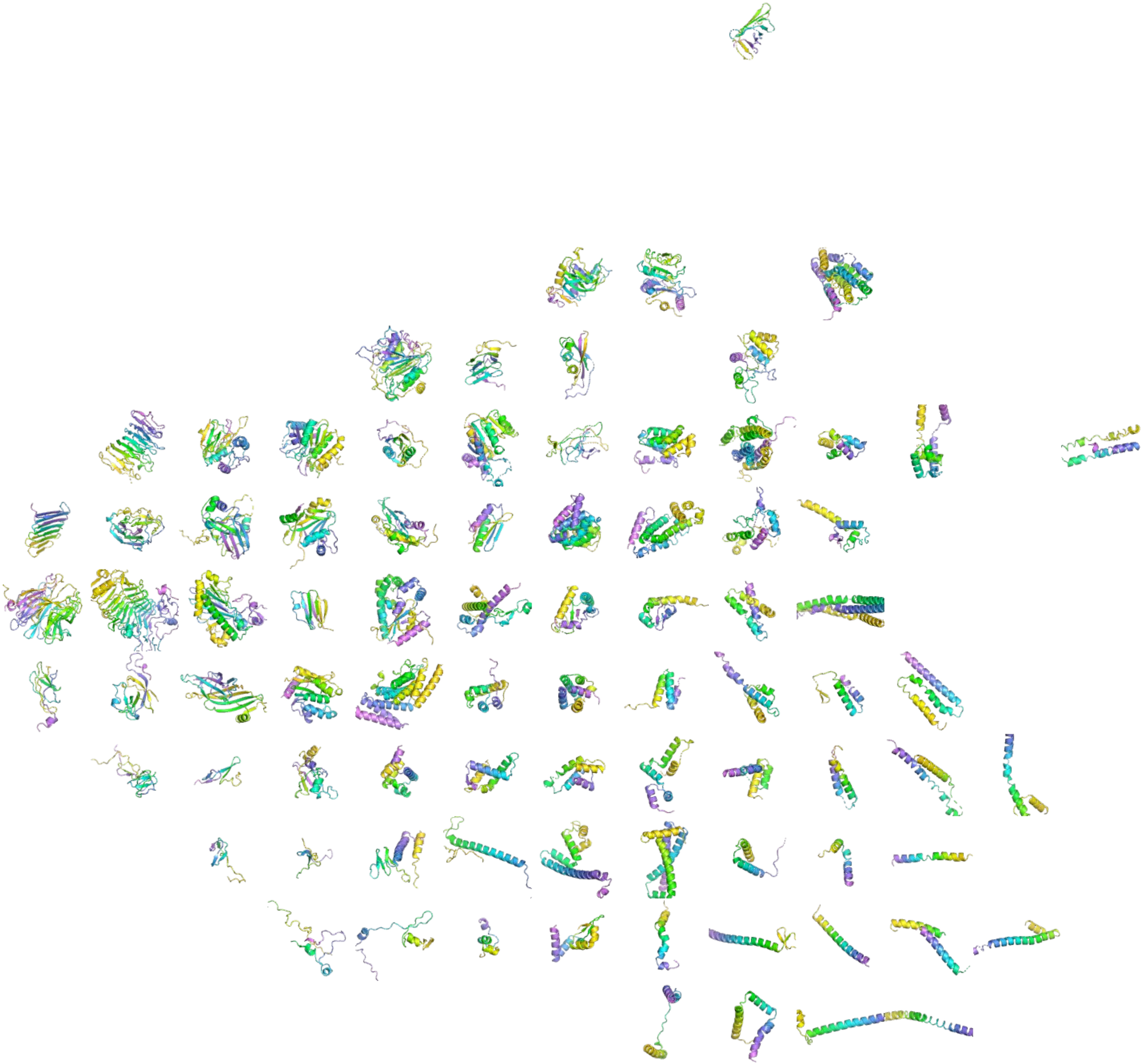

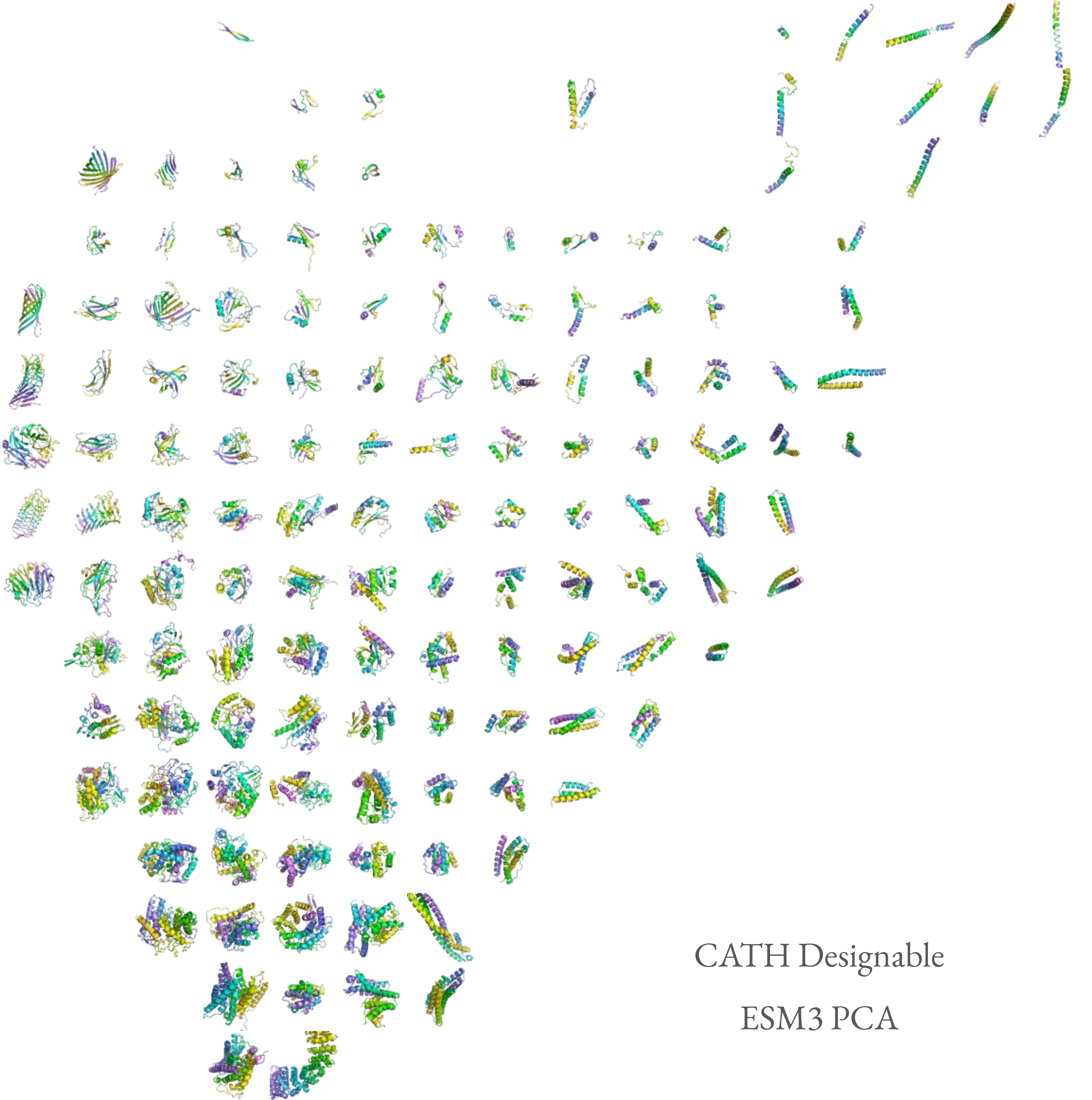

CATH Designable

ESM3 PCA

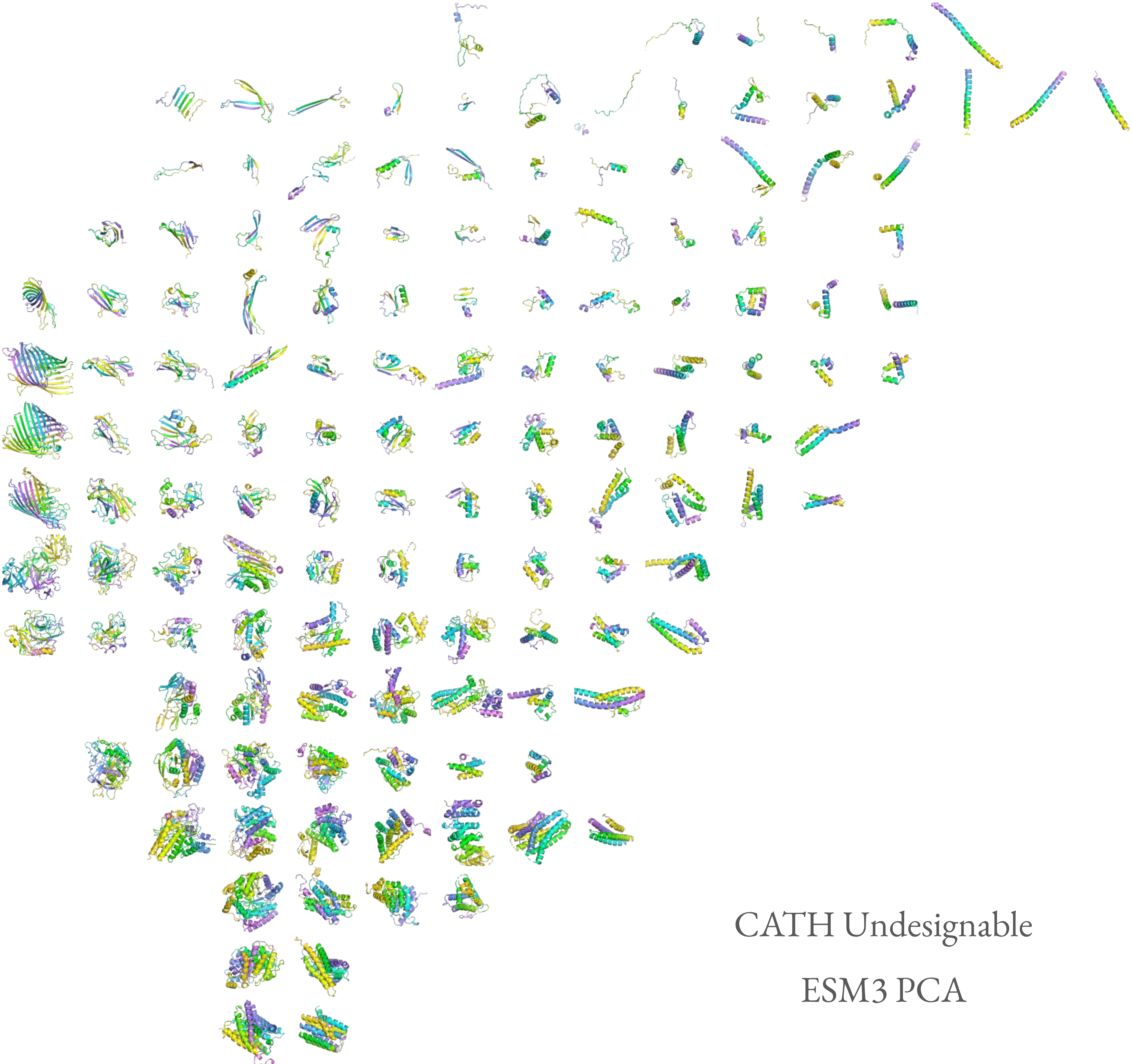

CATH Undesignable

ESM3 PCA

RFdiffusion Designable

ESM3 PCA

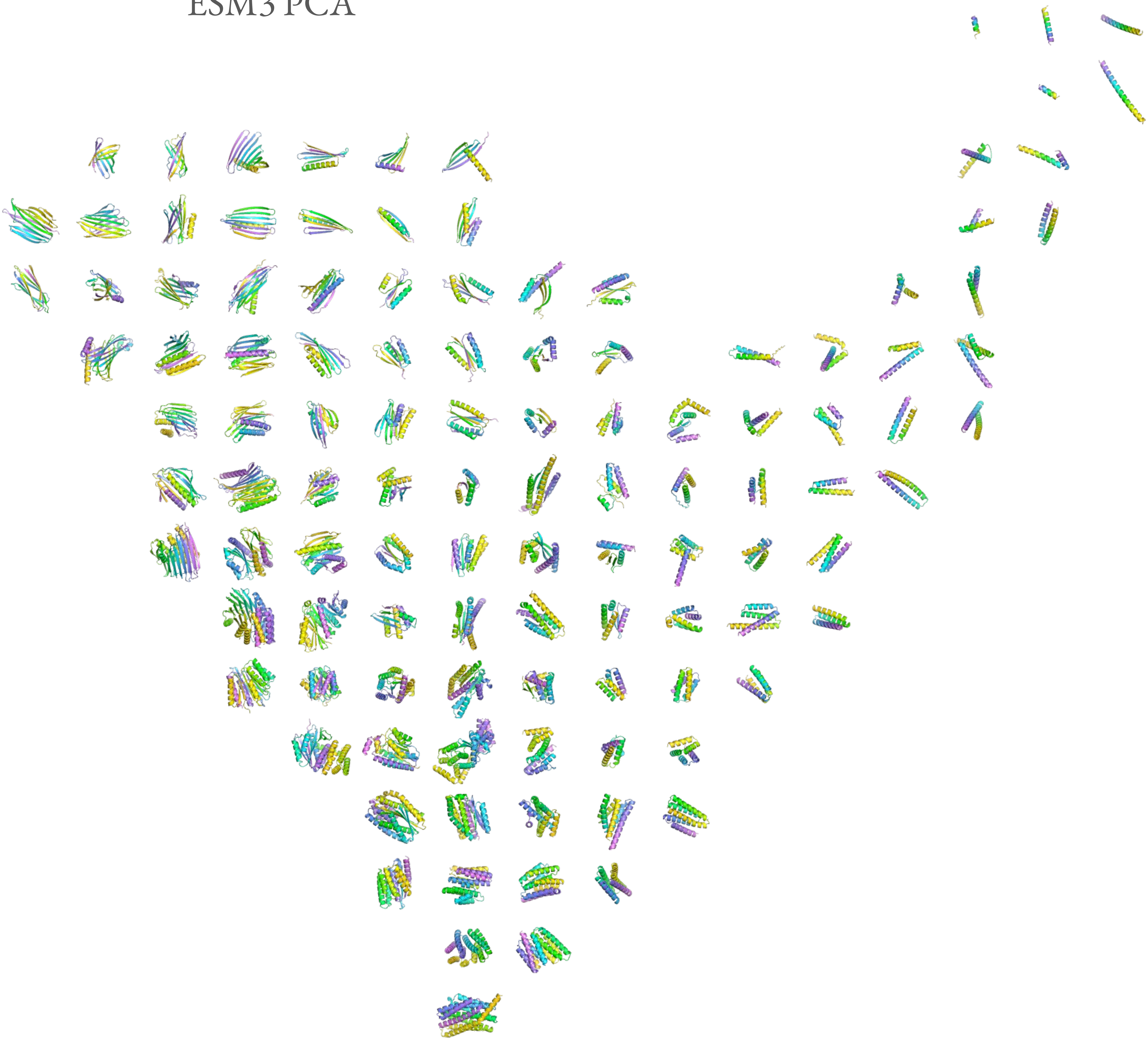

RFdiffusion Undesignable

ESM3 PCA

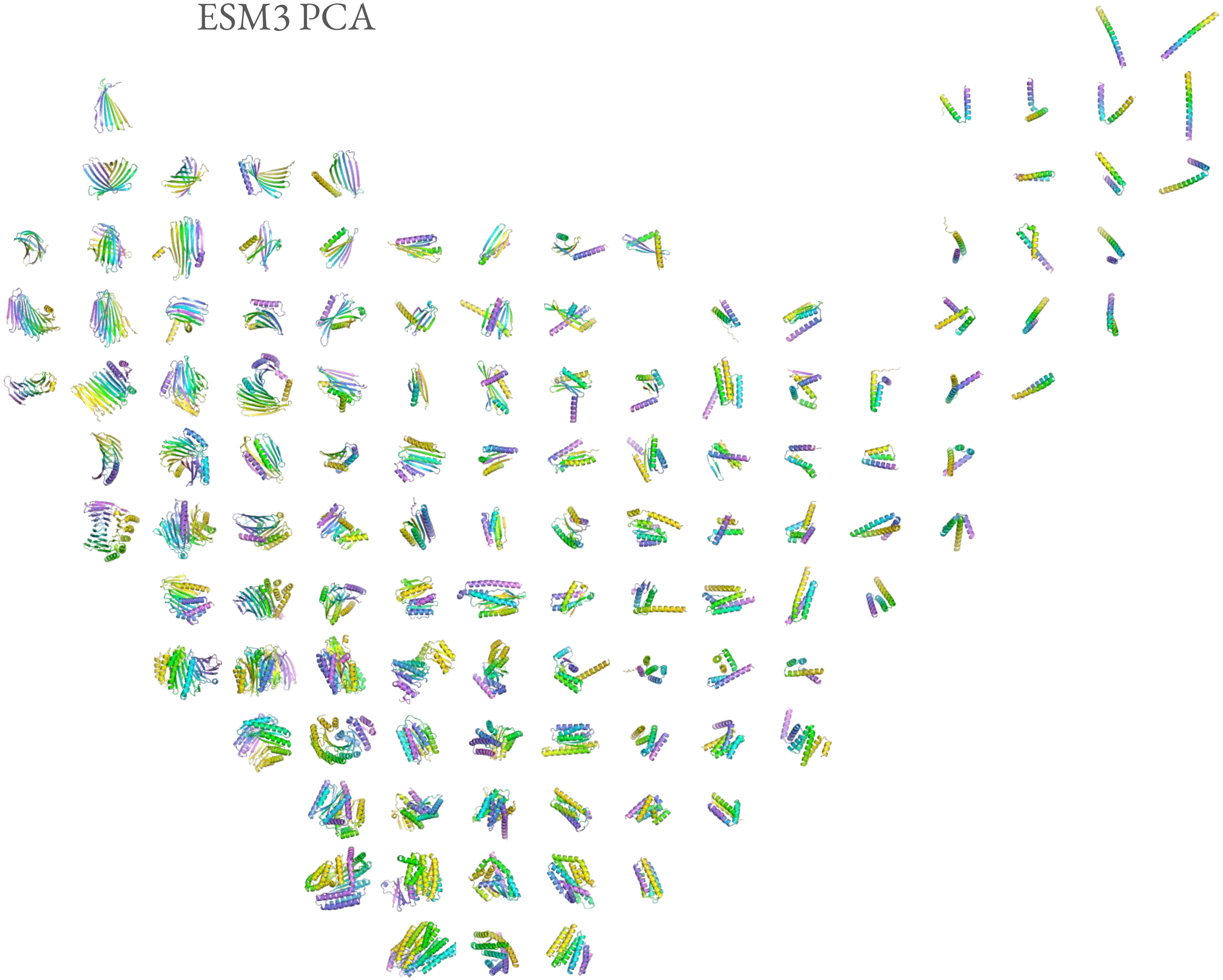

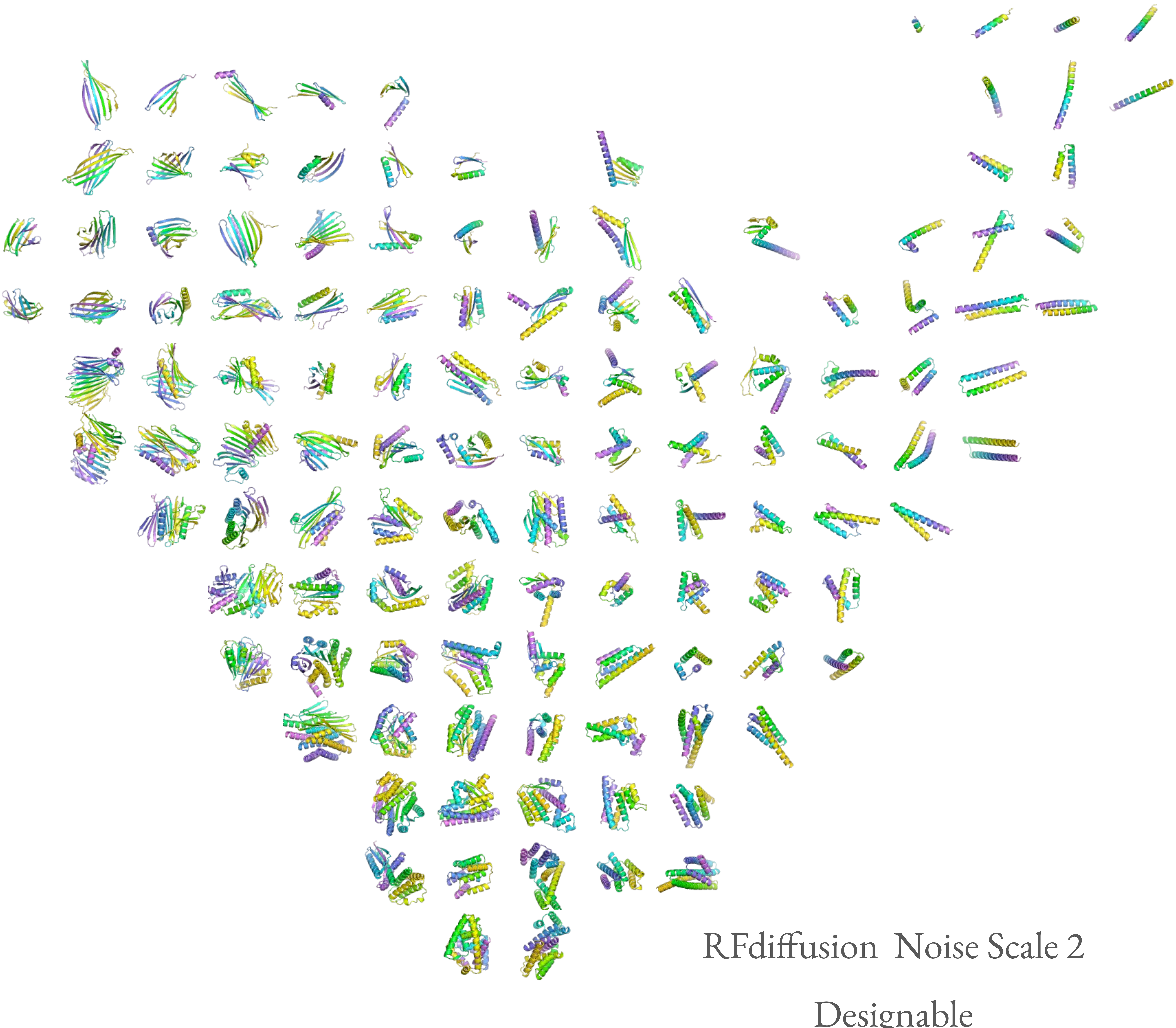

RFdiffusion  Noise Scale 2

Designable

ESM3 PCA

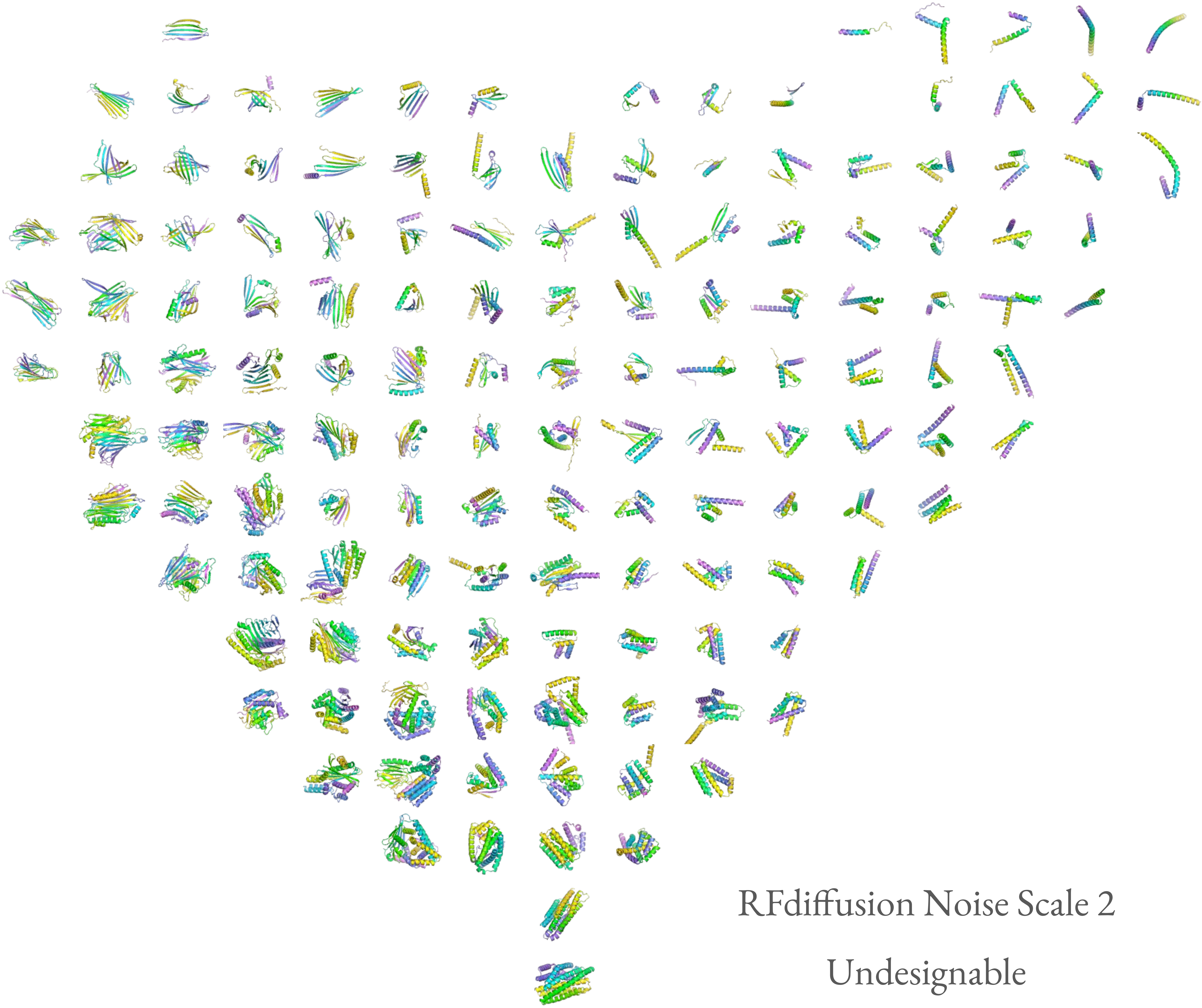

RFdiffusion Noise Scale 2

Undesignable

ESM3 PCA

RFdiffusion Noise Scale 3

Designable

ESM3 PCA

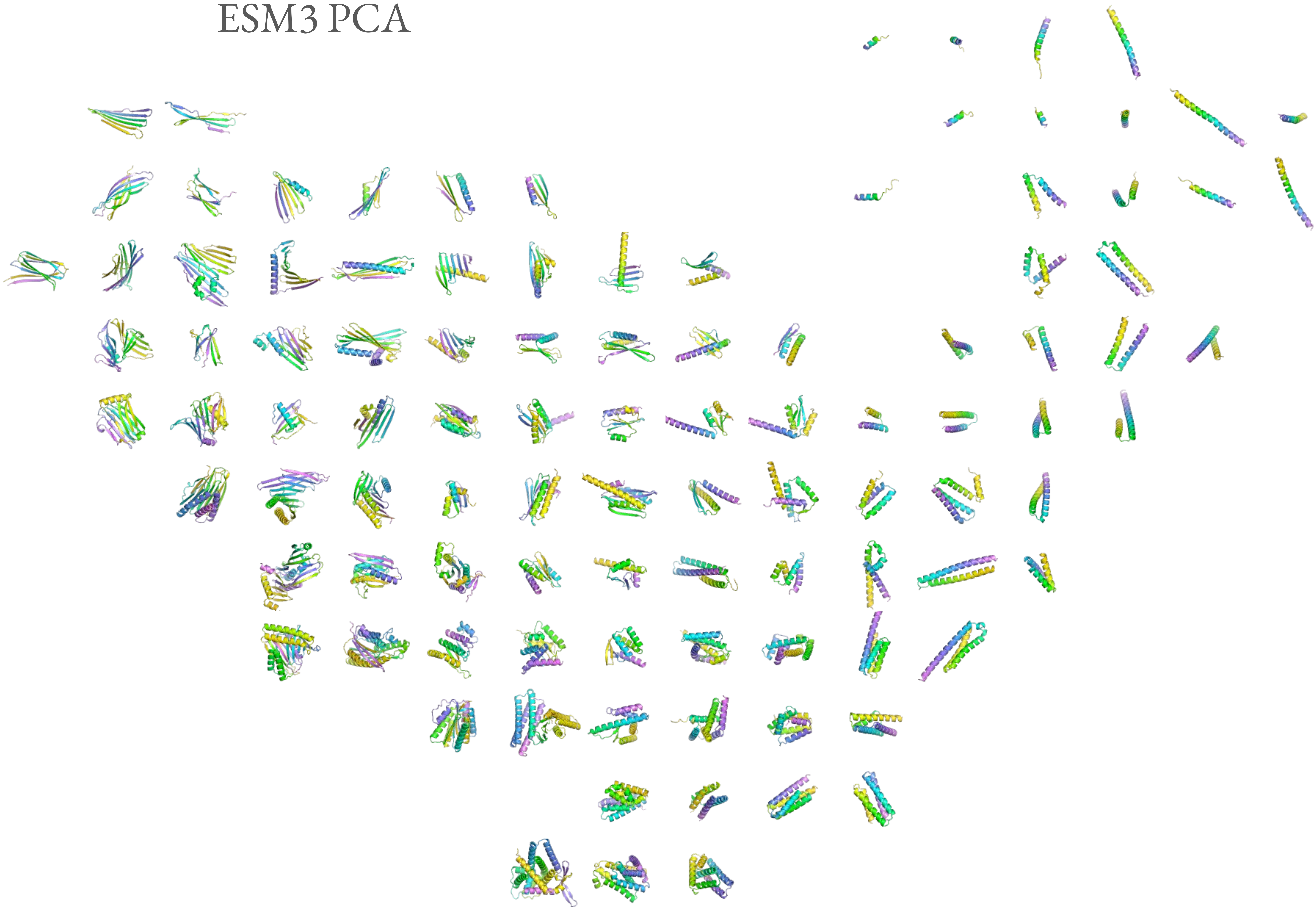

RFdiffusion Noise Scale 3

Undesignable

ESM3 PCA

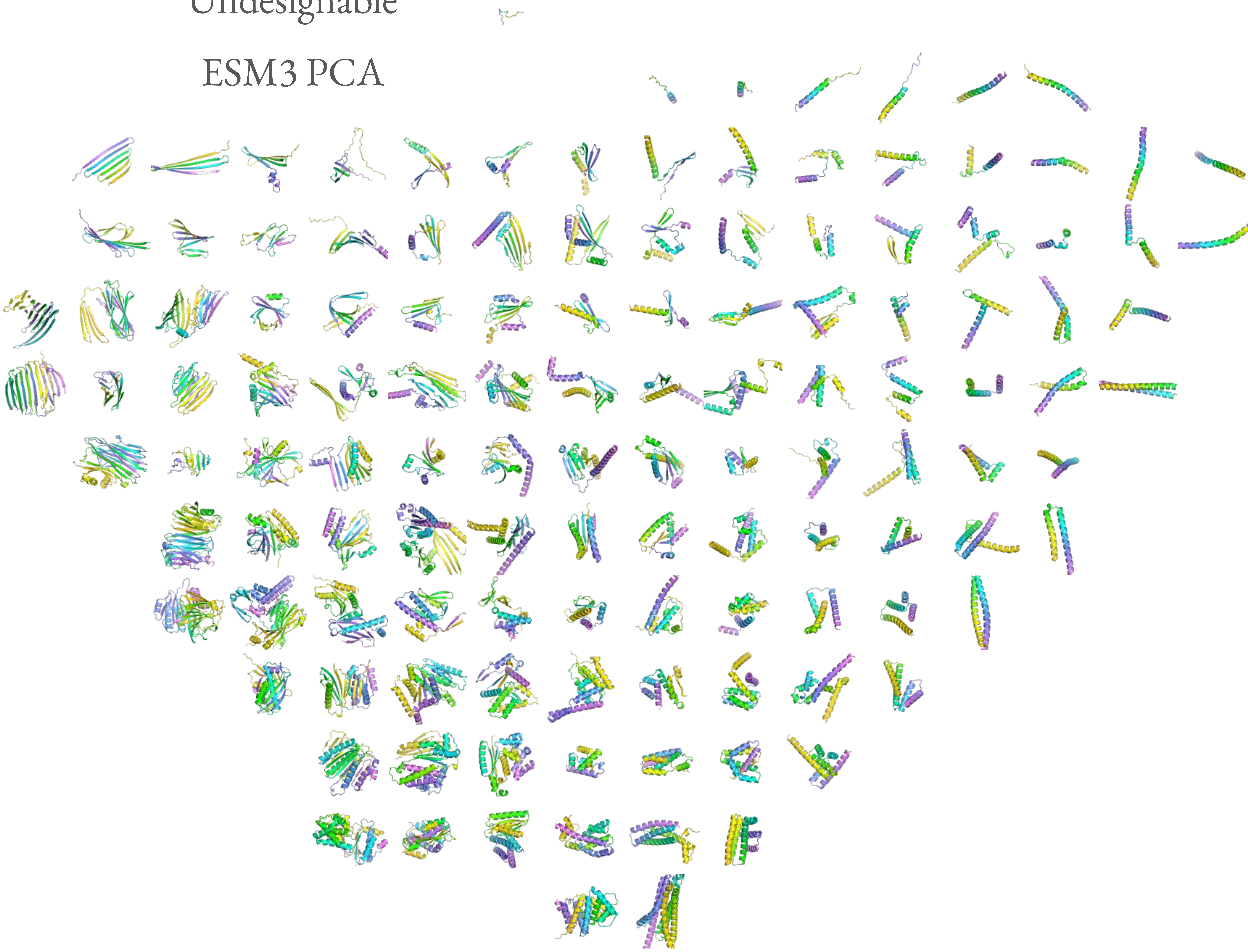

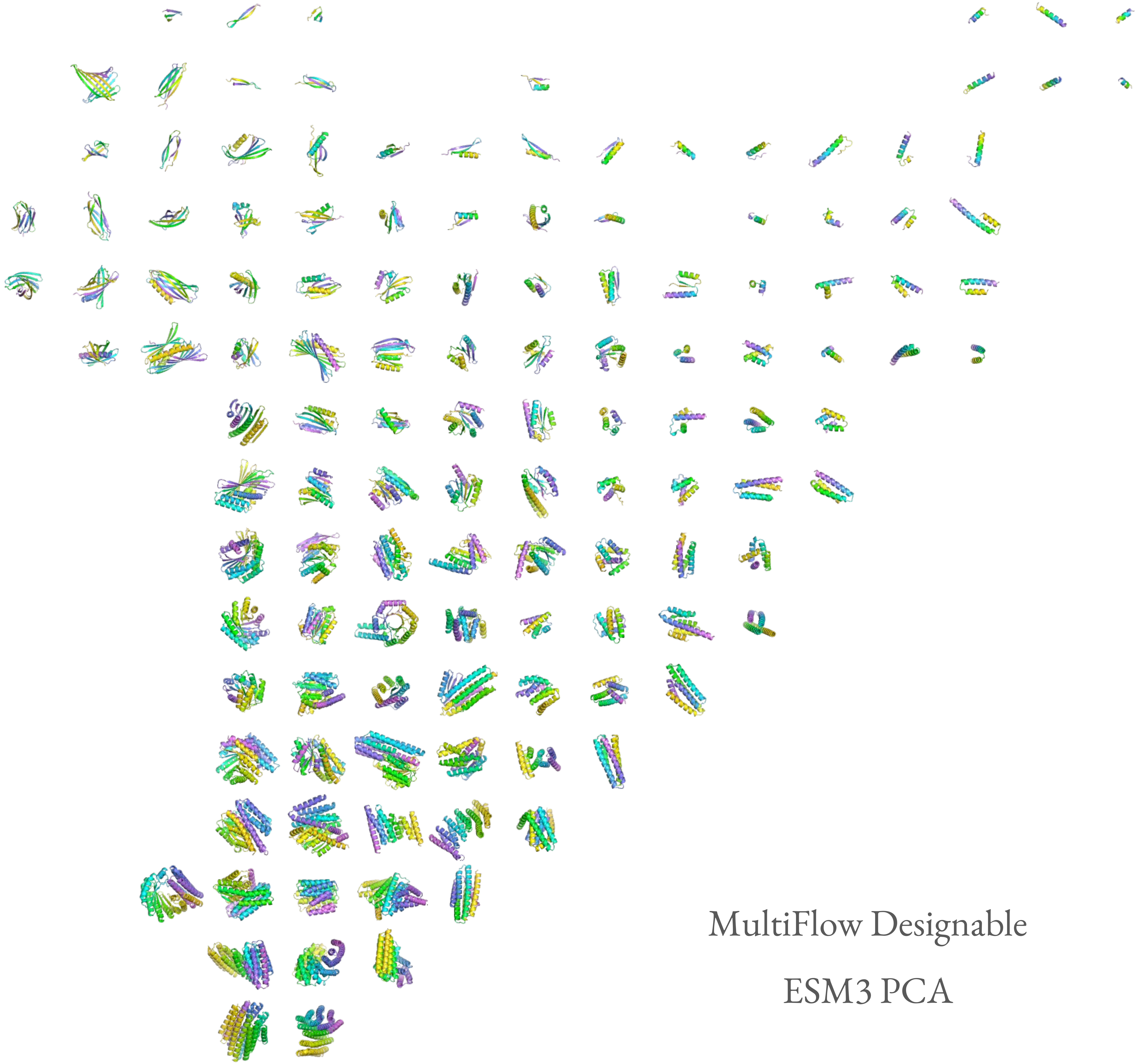

MultiFlow Designable

ESM3 PCA

MultiFlow Undesignable

ESM3 PCA

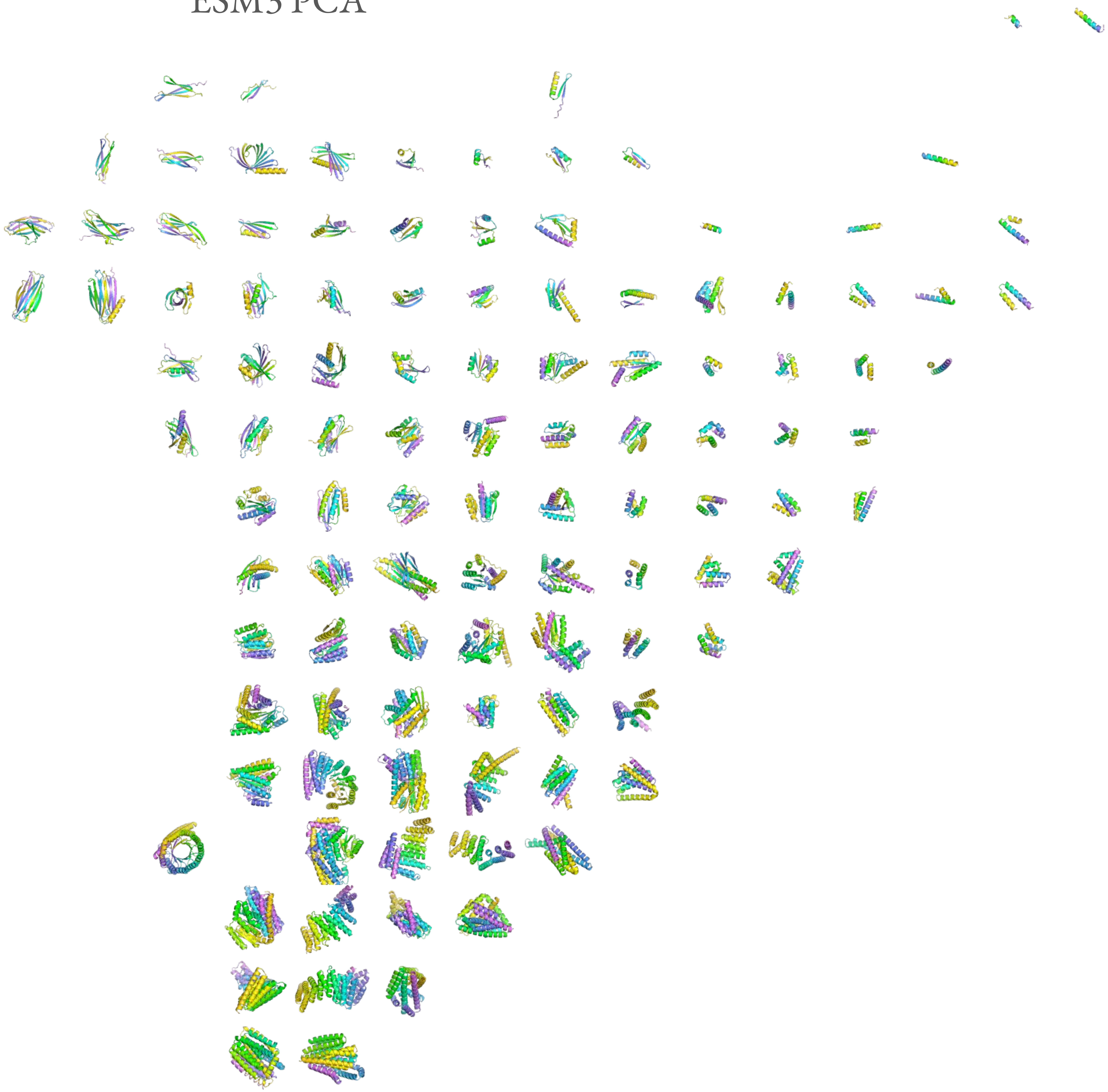

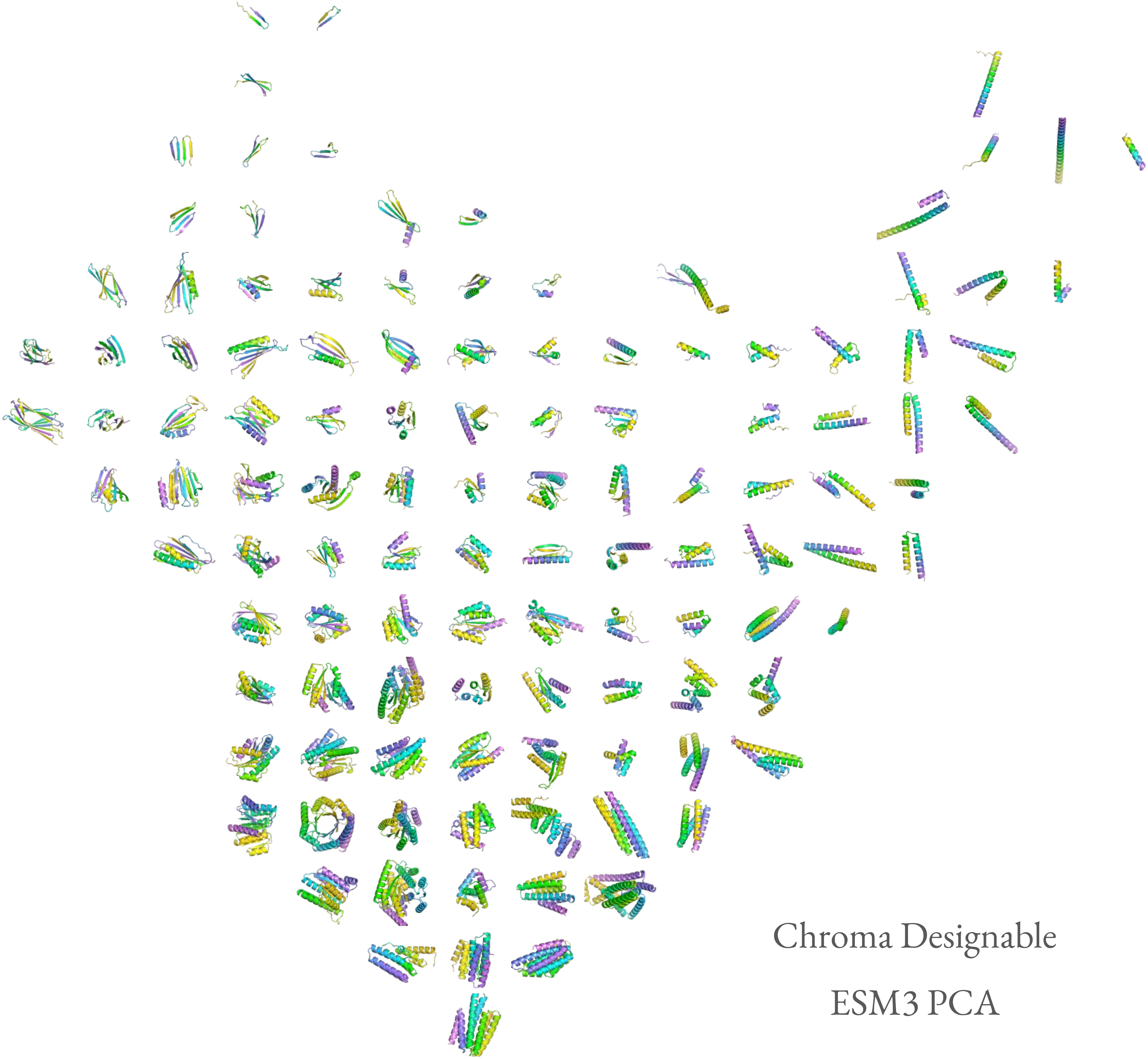
Chroma Designable
ESM3 PCA

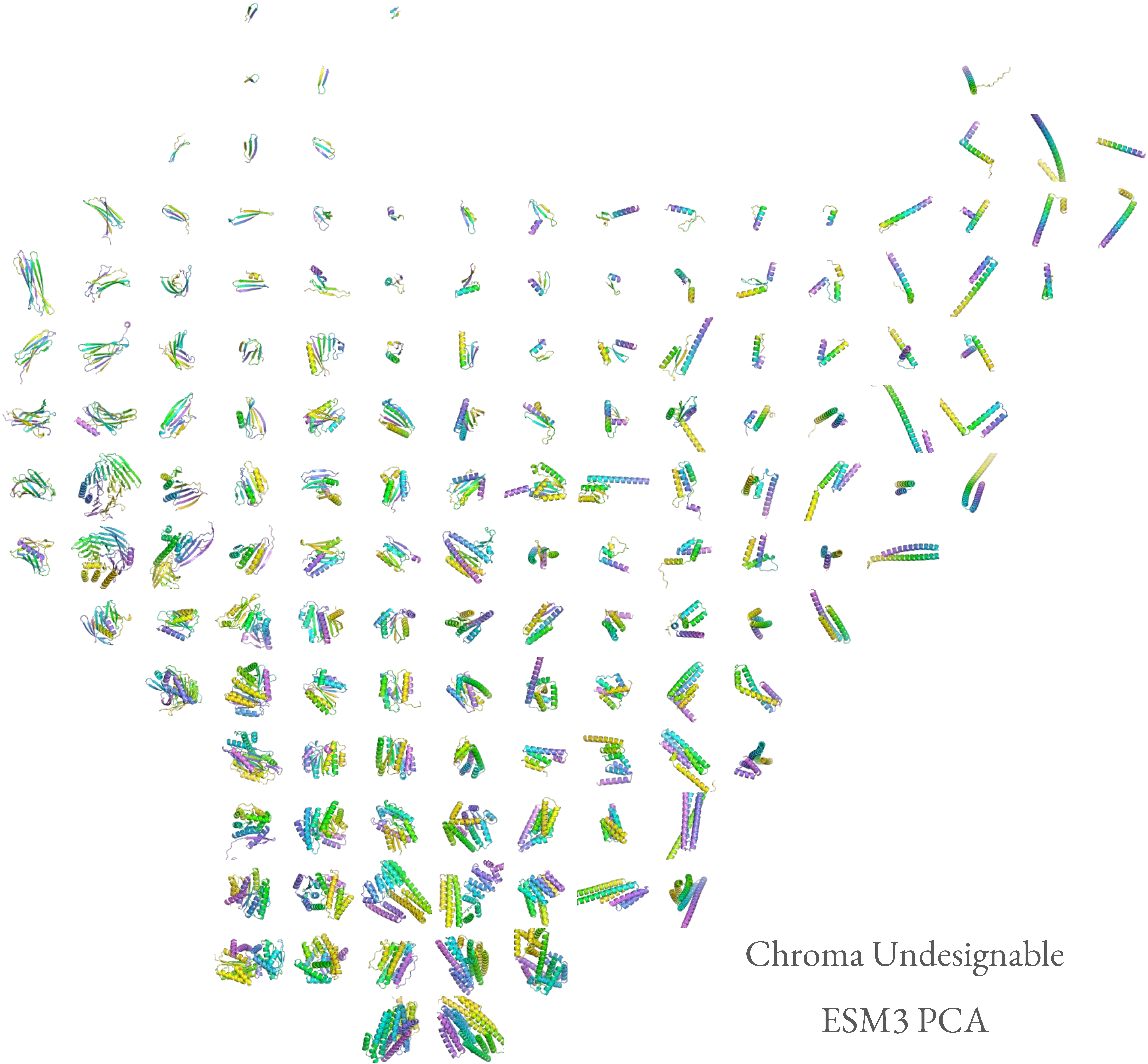

Chroma Undesignable

ESM3 PCA

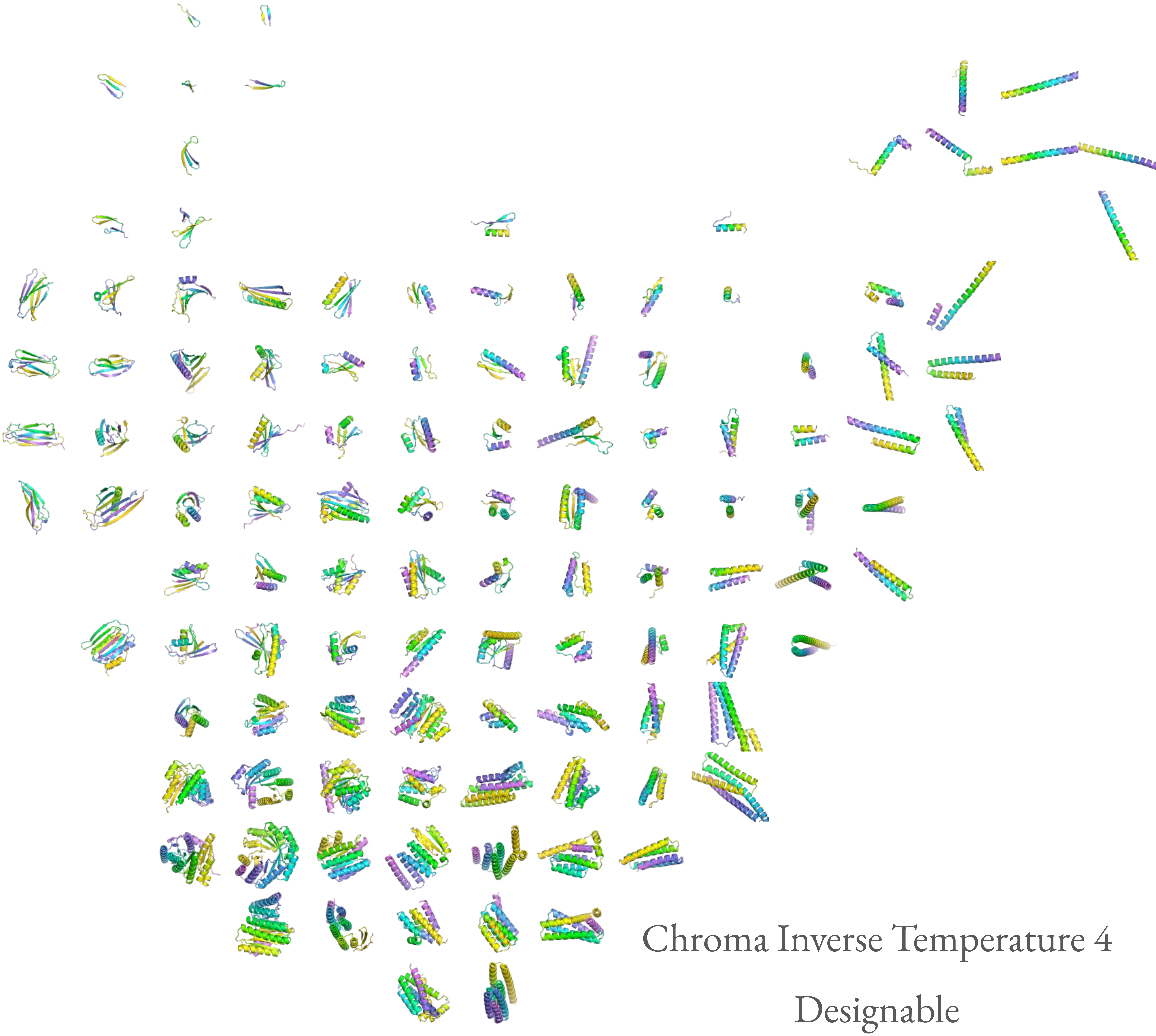

Chroma Inverse Temperature 4

Designable

ESM3 PCA

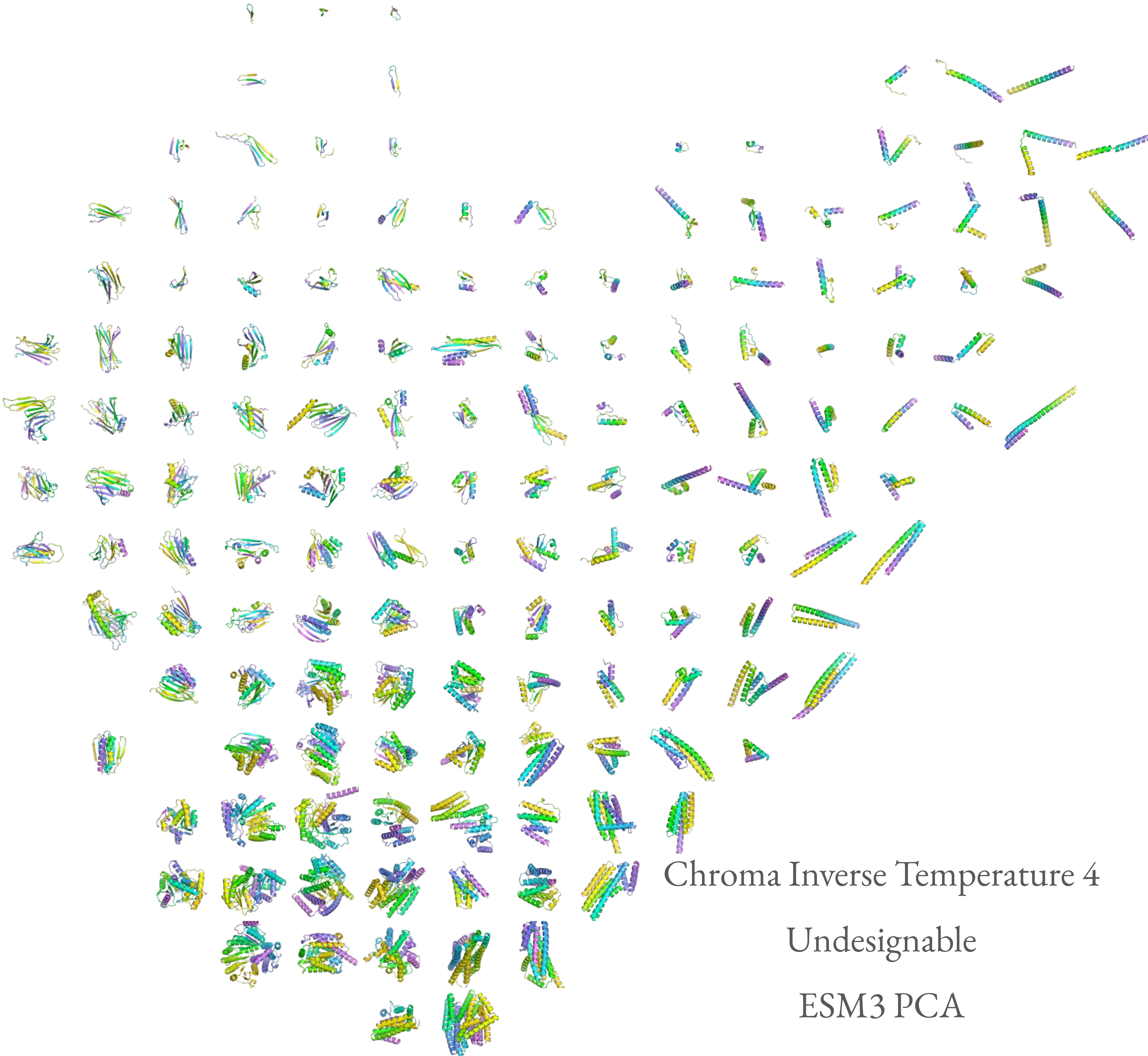

Chroma Inverse Temperature 4
Undesignable
ESM3 PCA

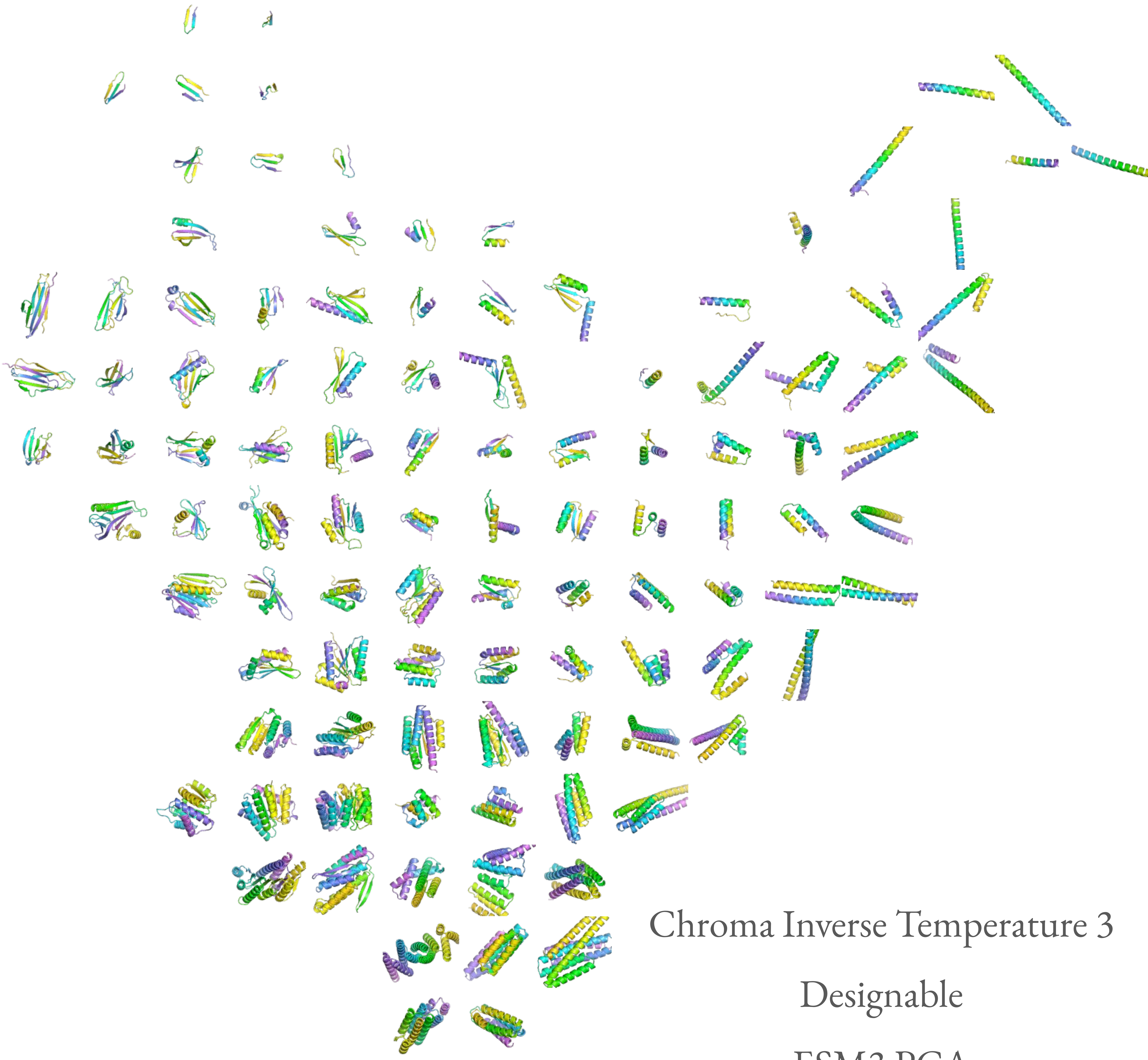

Chroma Inverse Temperature 3

Designable

ESM3 PCA

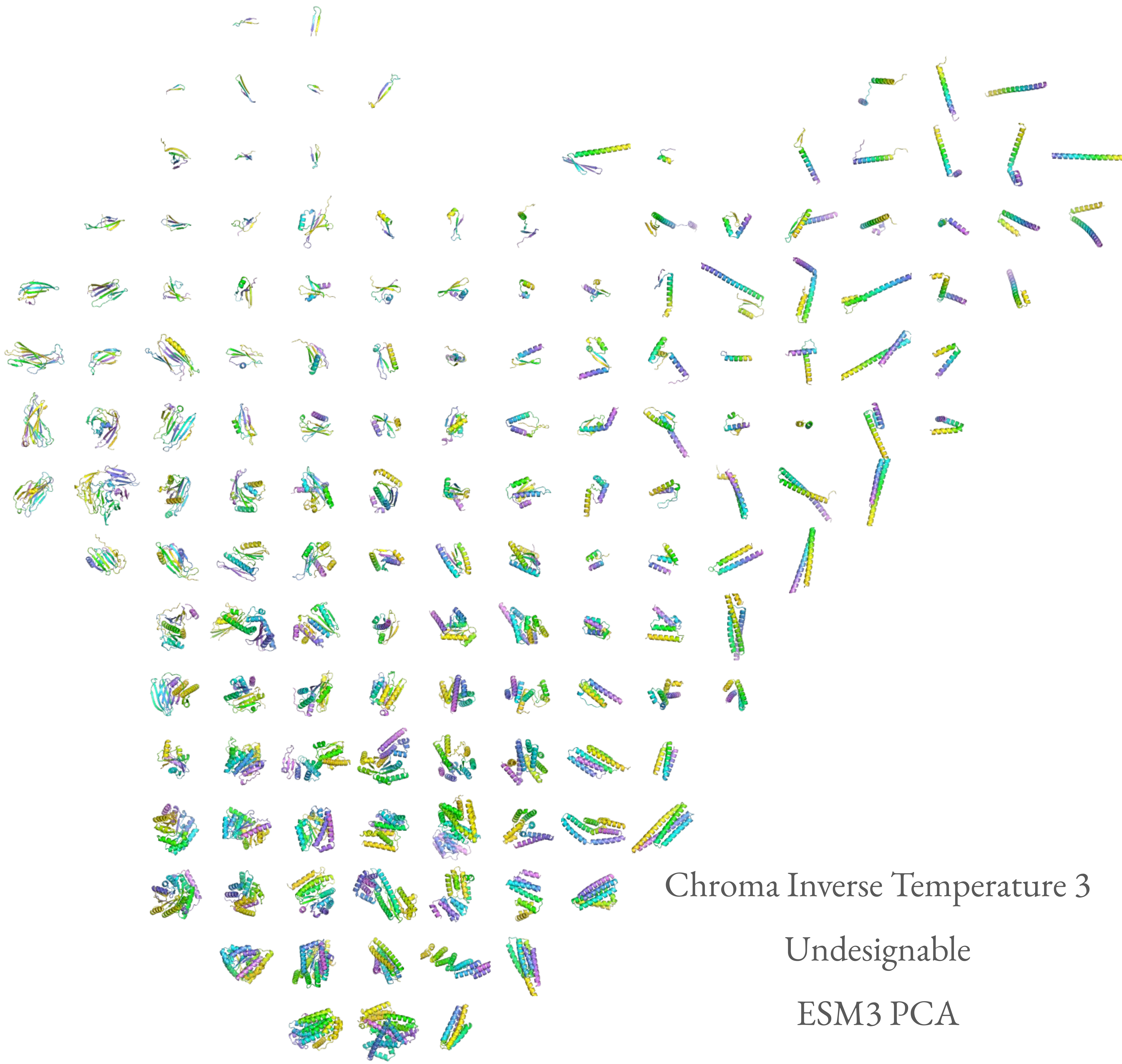
Chroma Inverse Temperature 3
Undesignable
ESM3 PCA

Genie2 Designable

ESM3 PCA

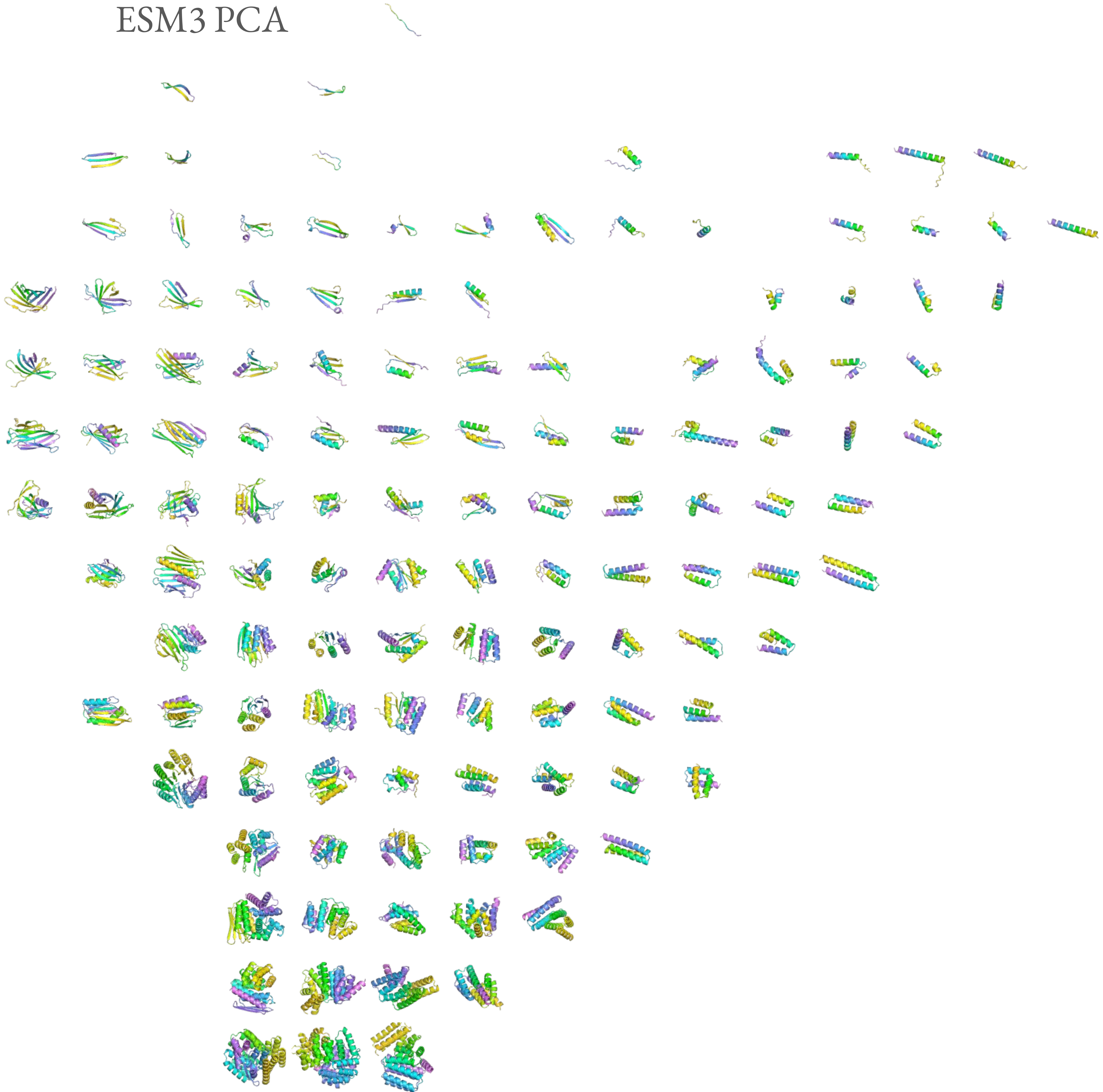

Genie2 Undesignable

ESM3 PCA

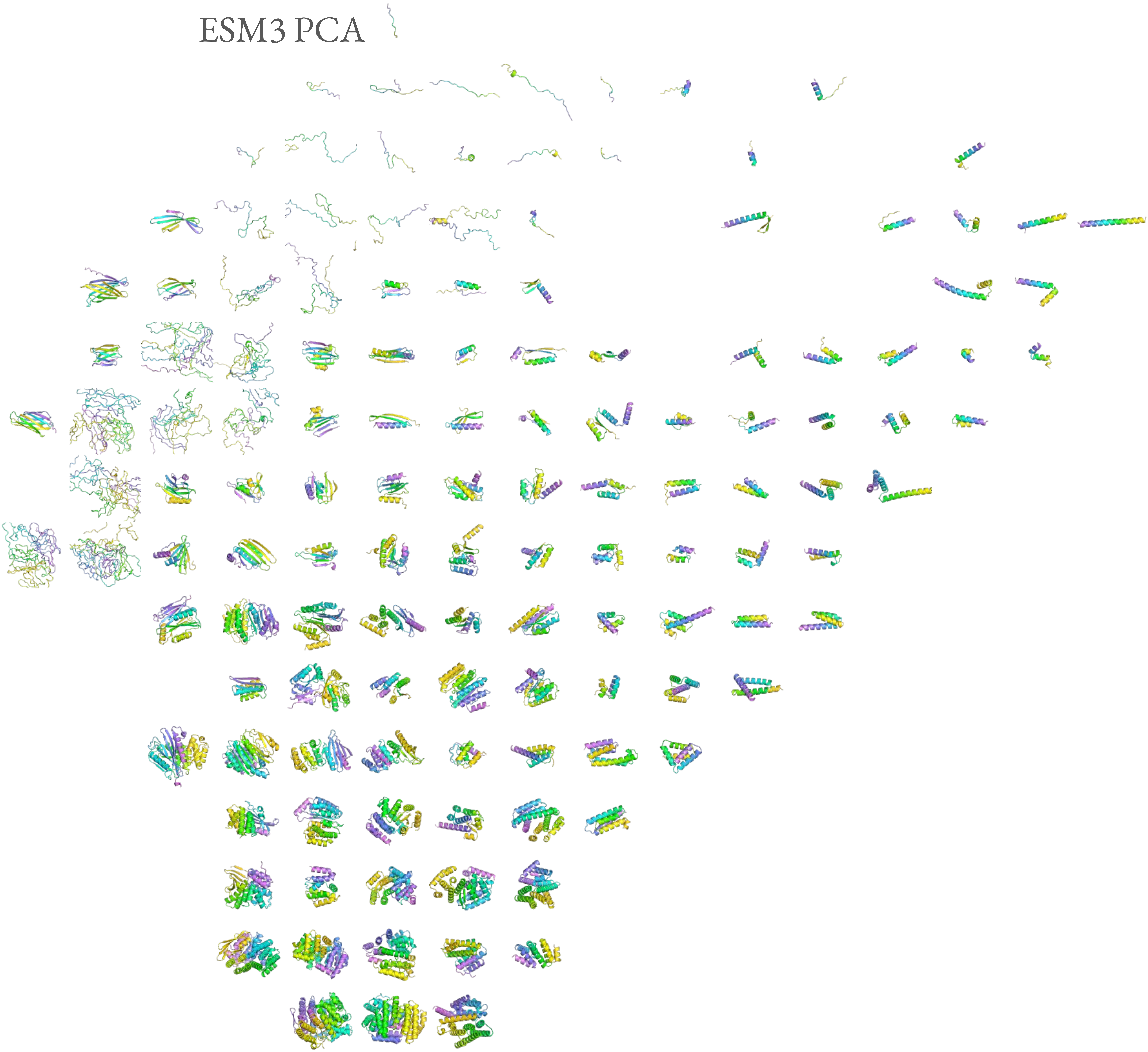

Genie2 Scale 0.8 Designable

ESM3 PCA

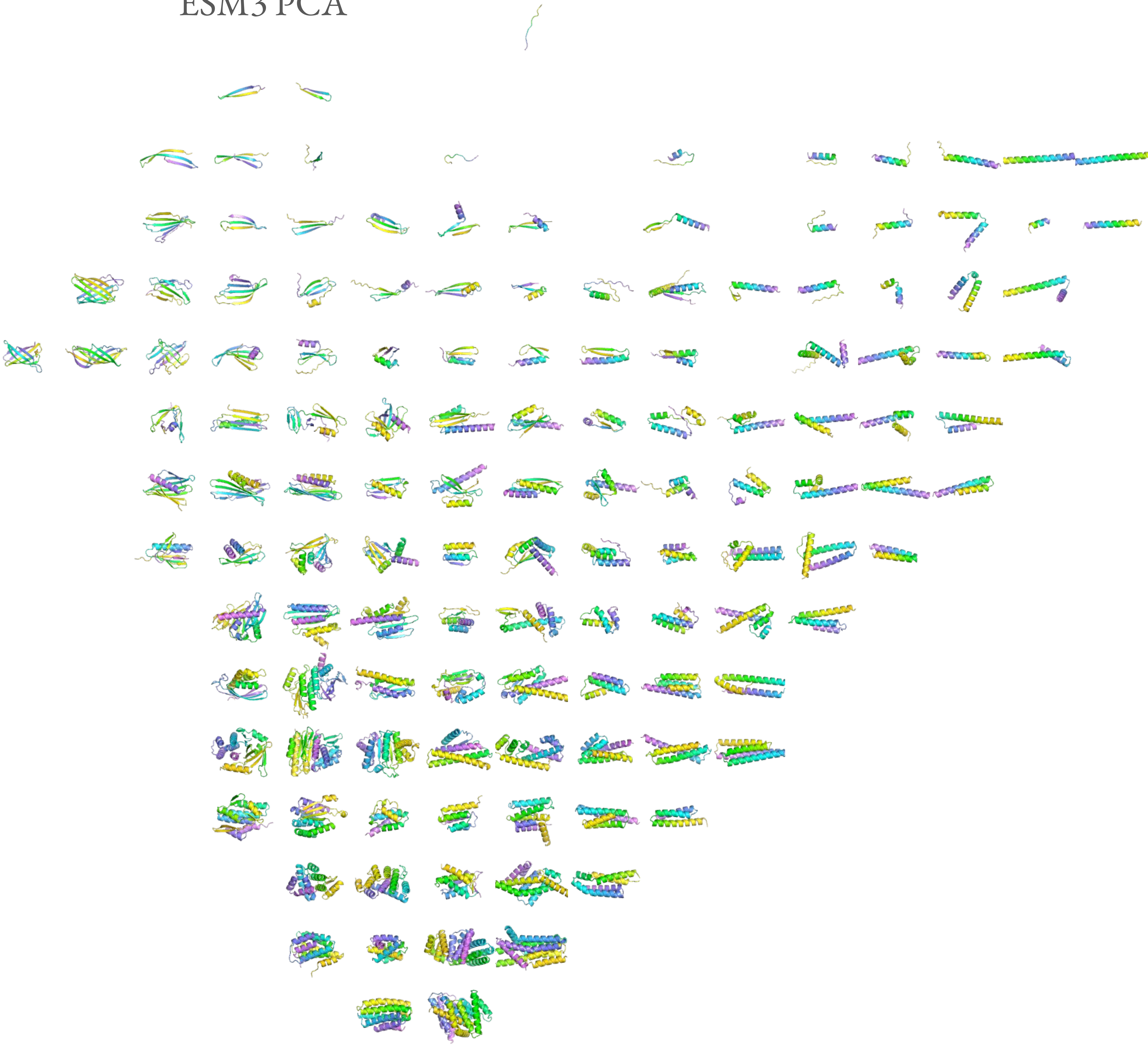

Genie2 Scale 0.8 Undesignable

ESM3 PCA

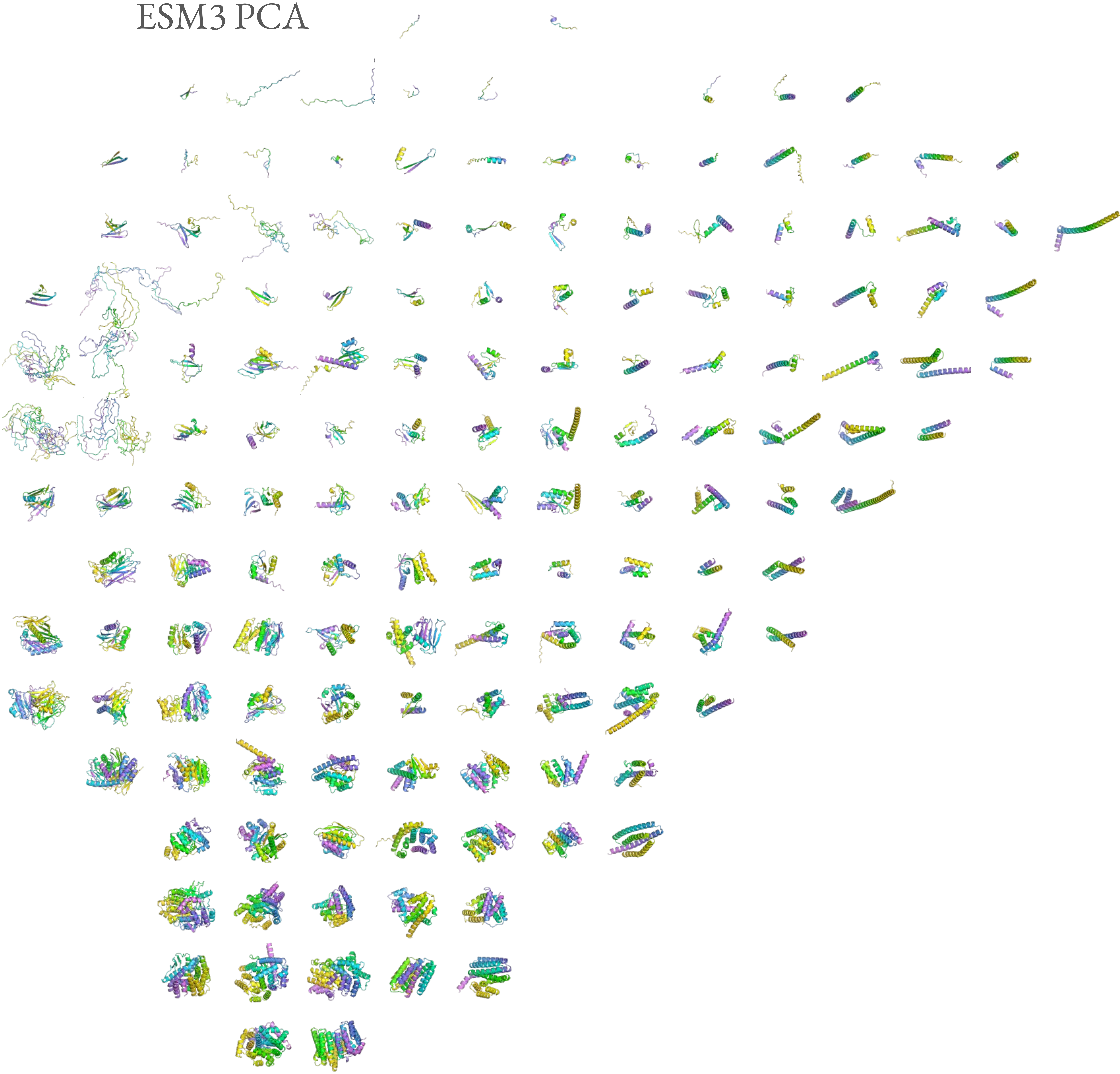

Genie2 Scale 1.0 Designable

ESM3 PCA

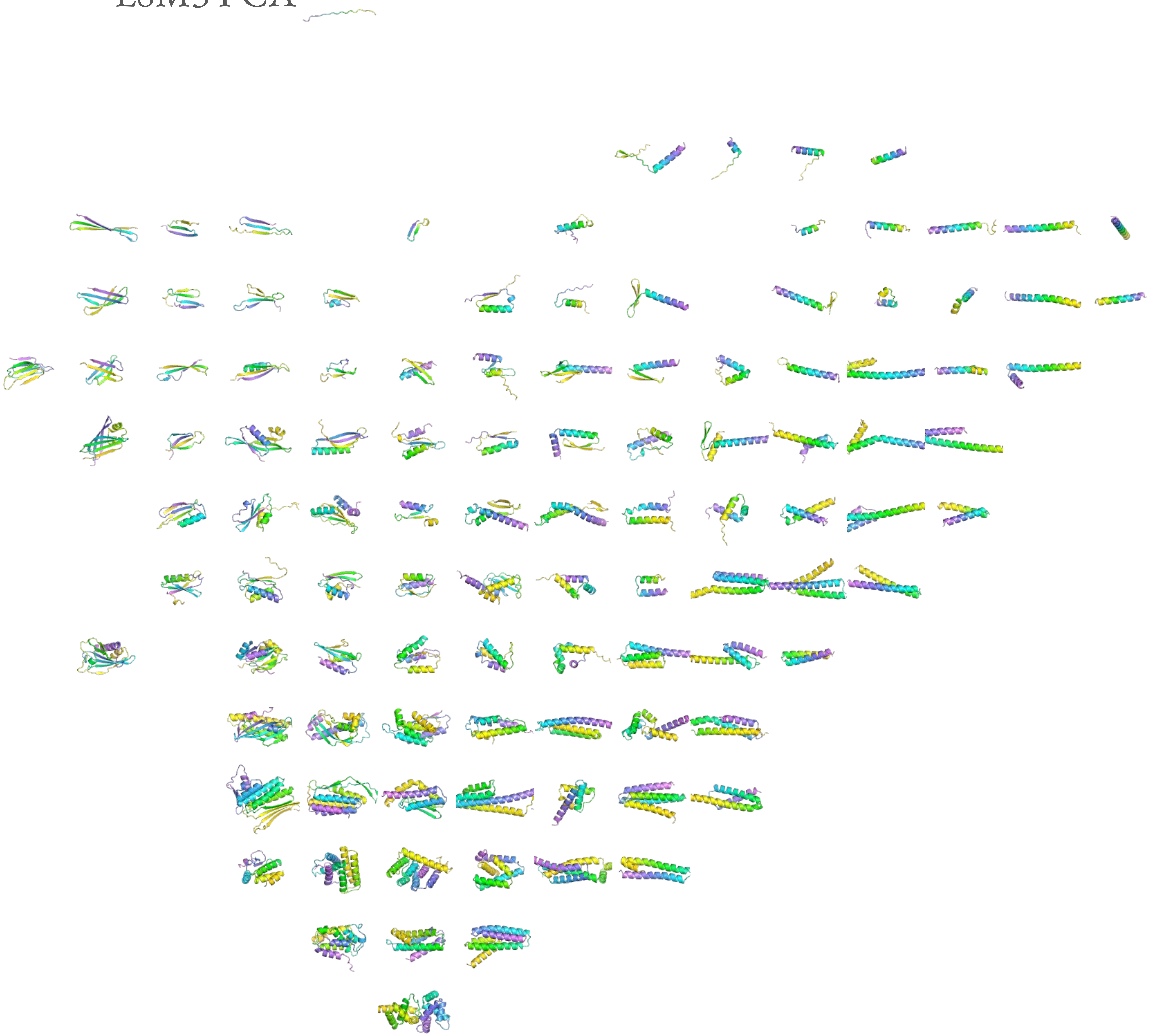

Genie2 Scale 1.0  Undesignable

ESM3 PCA

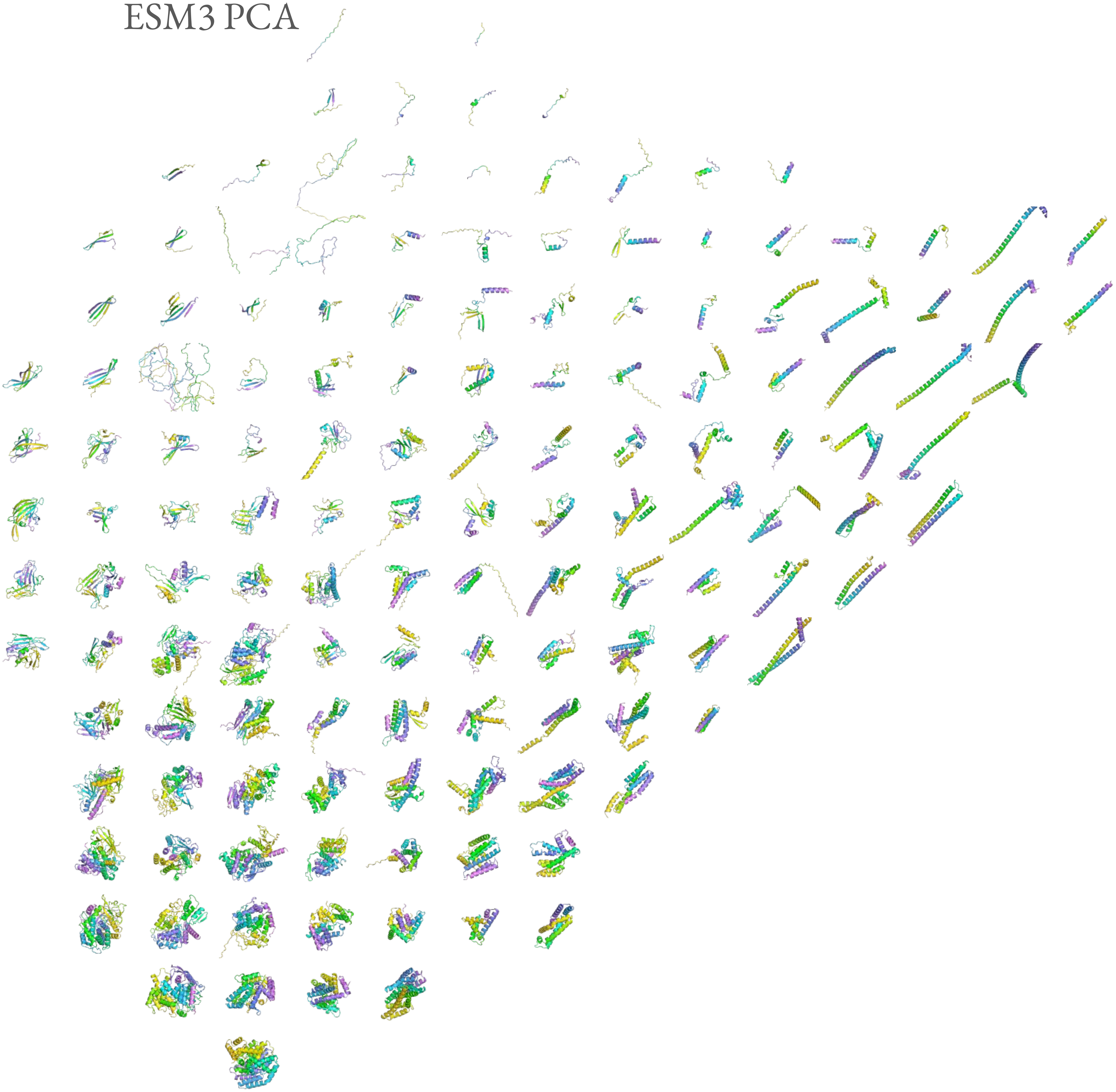

Protpardelle Stepscale 1.2 Designable

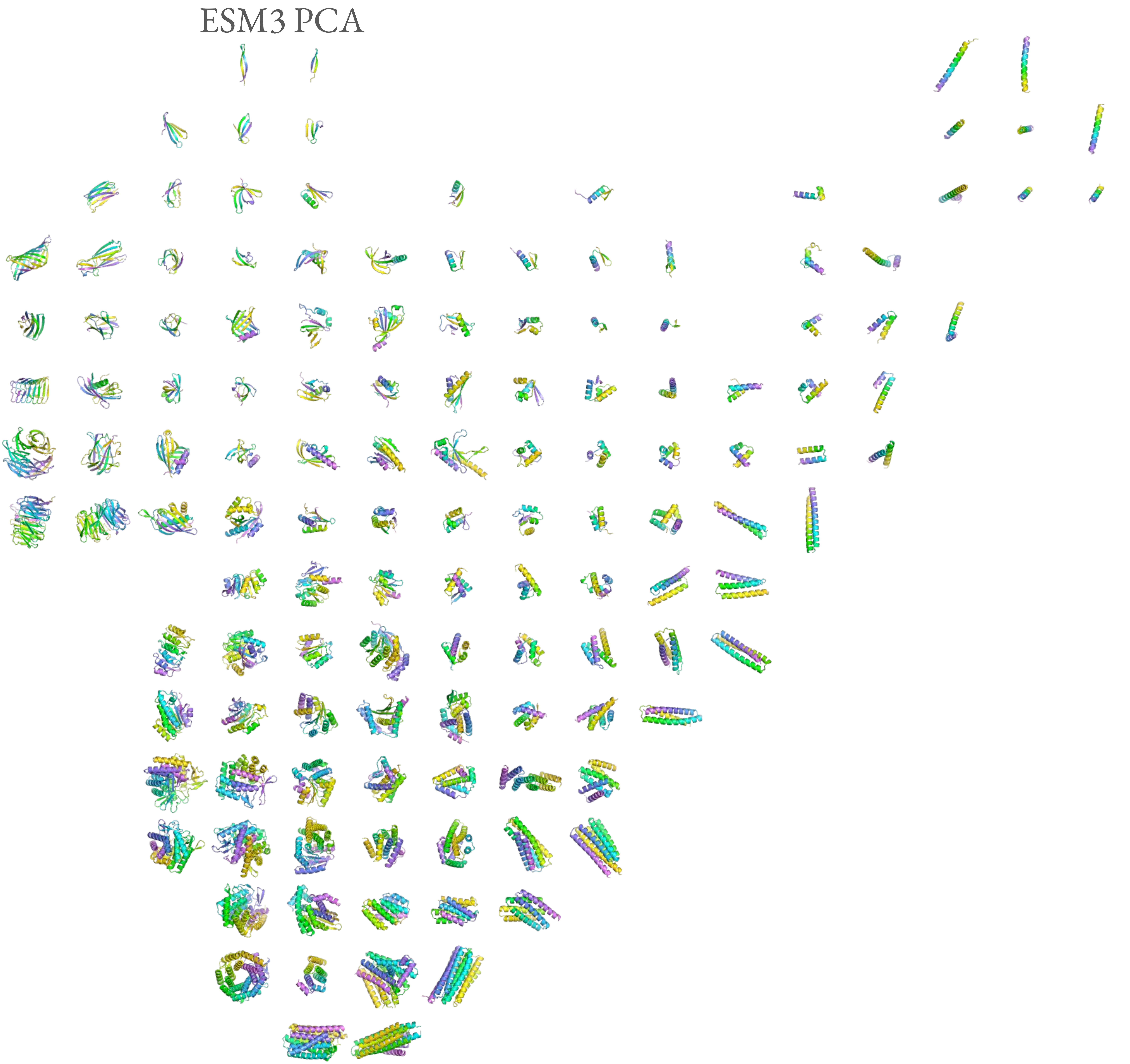

### Protpardelle Stepscale 1.2 Undesignable

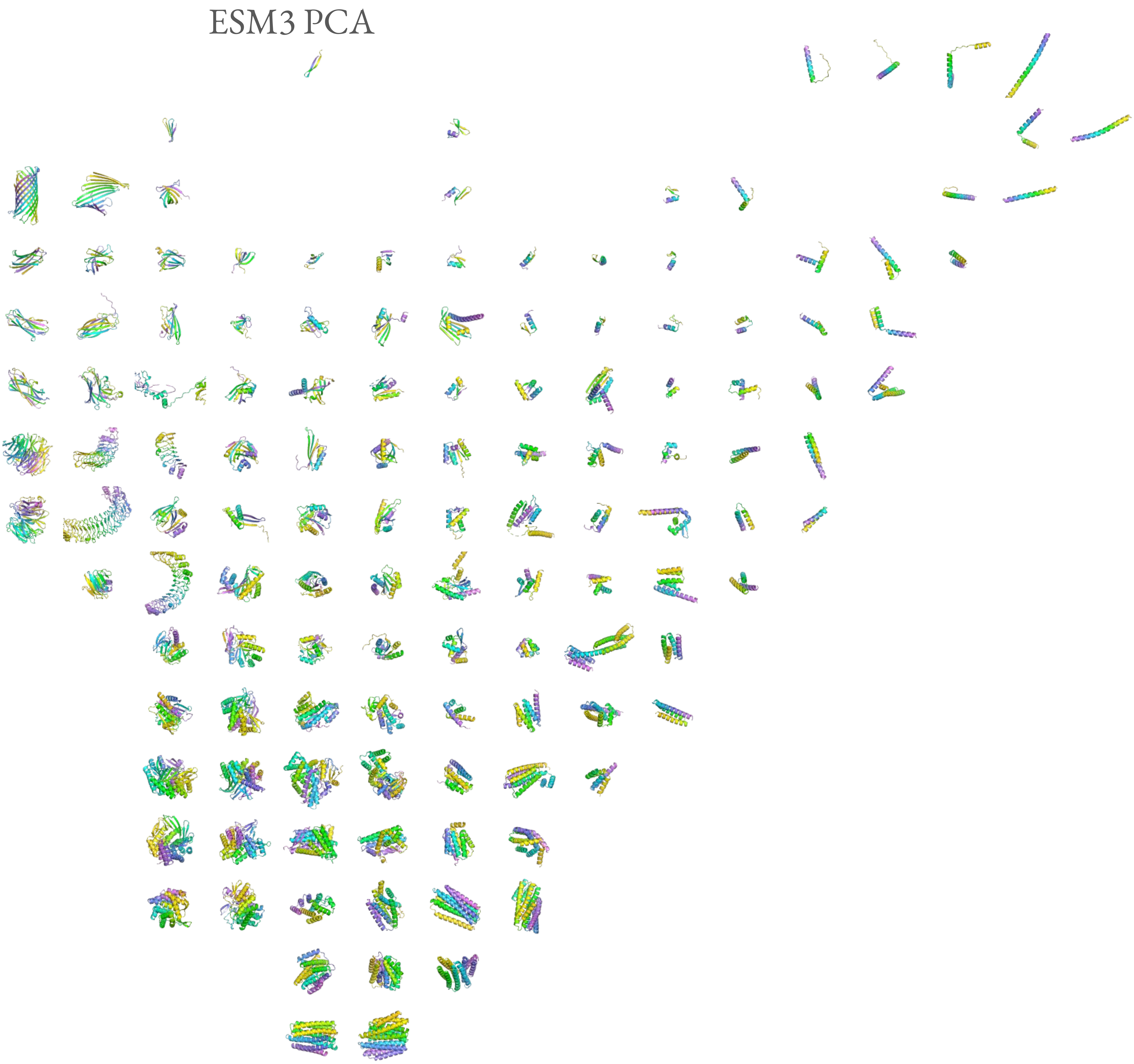

Protpardelle Stepscale 1.0 Designable

ESM3 PCA

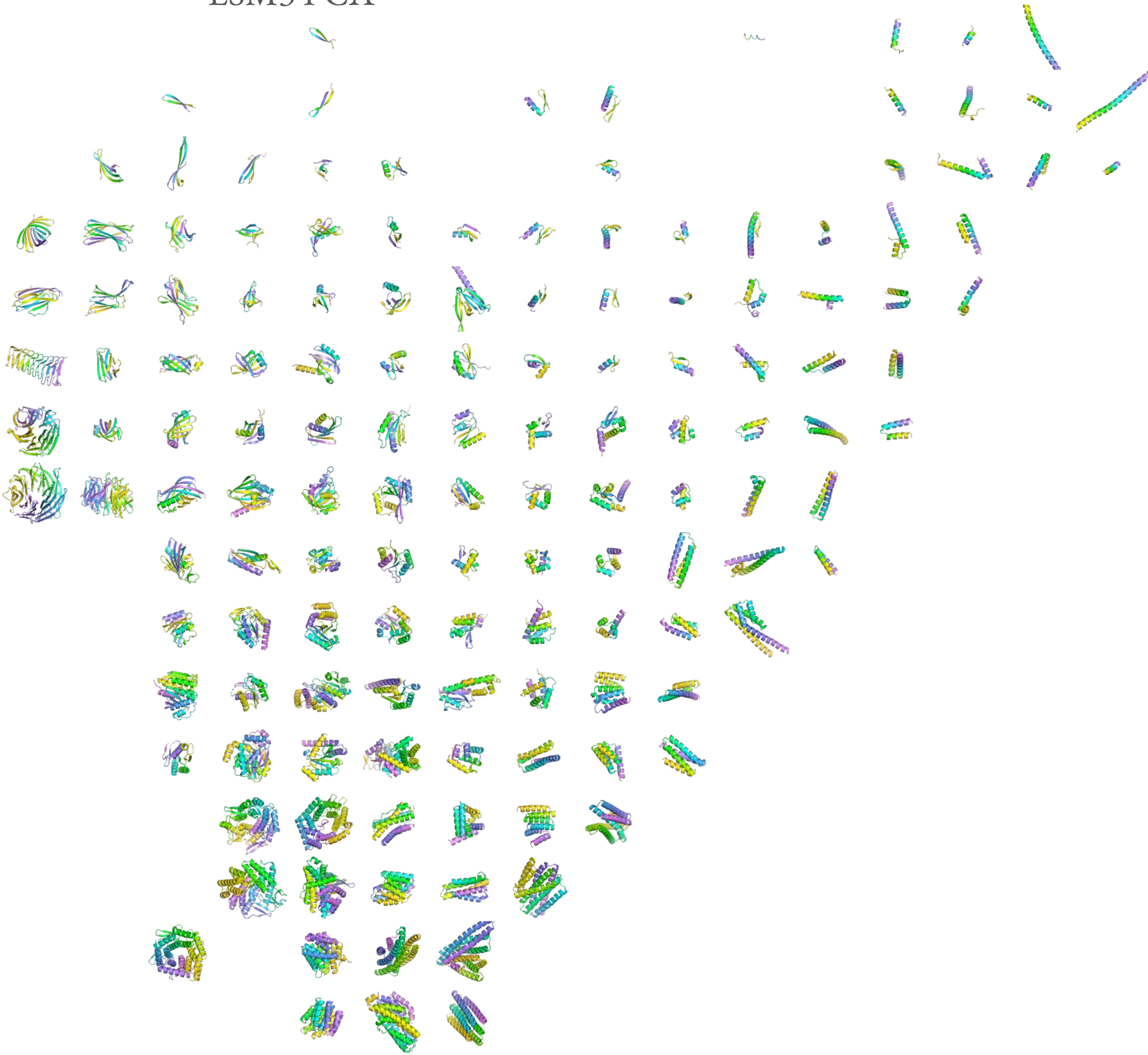

Protpardelle Stepscale 1.0 Undesignable

ESM3 PCA

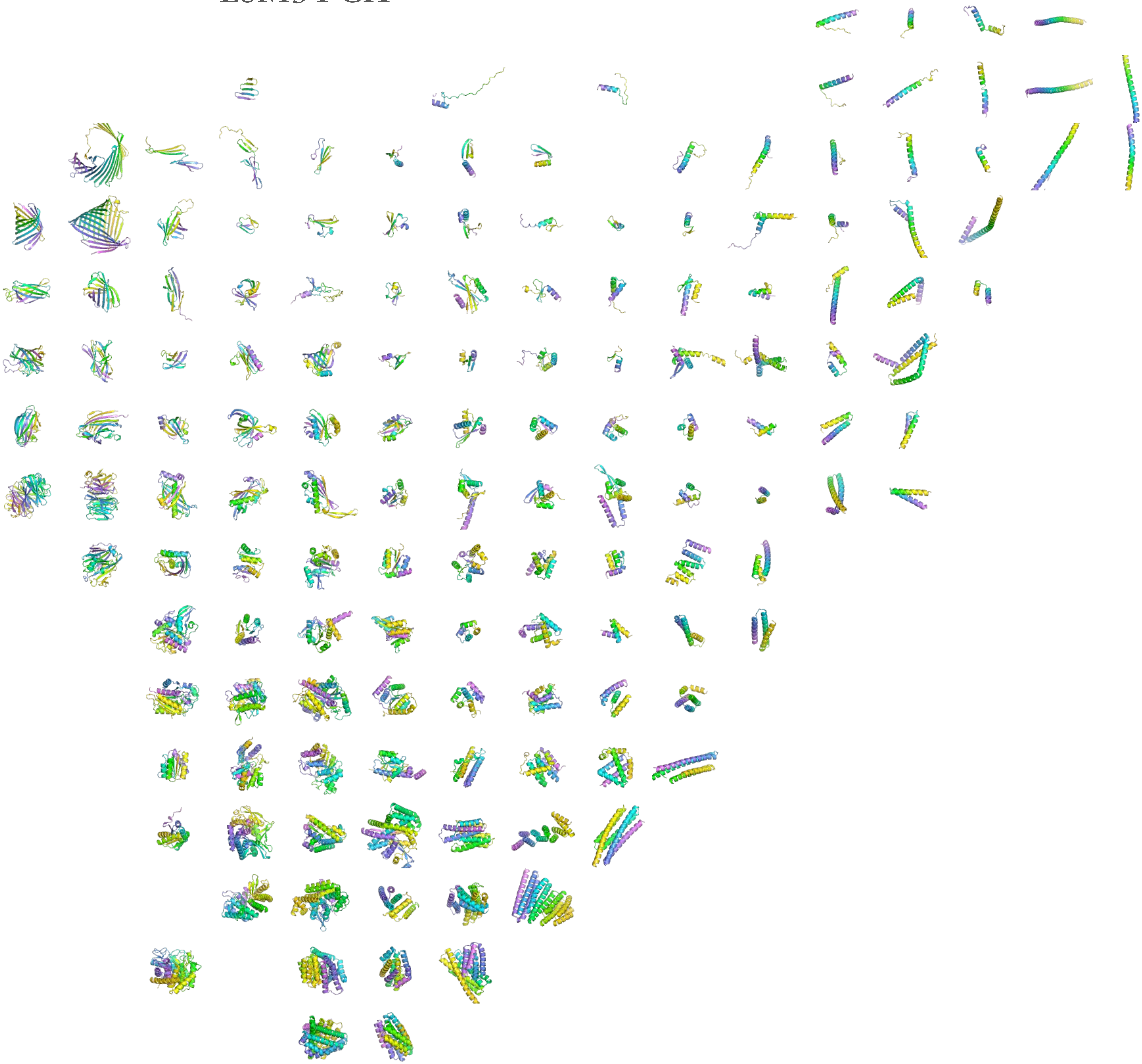

Protpardelle Stepscale 0.8 Designable

ESM3 PCA

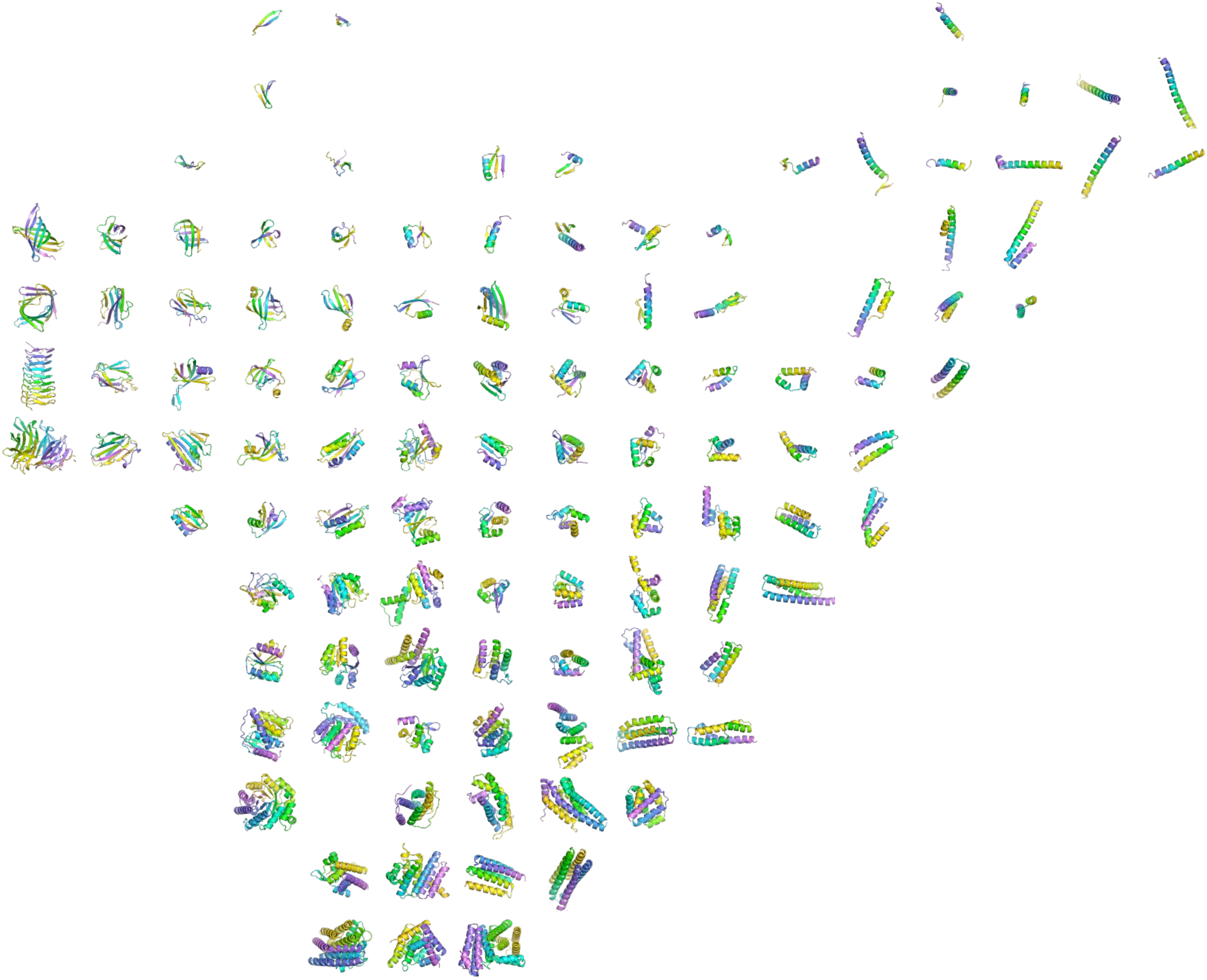

### Protpardelle Stepscale 0.8 Undesignable

#### ESM3 PCA

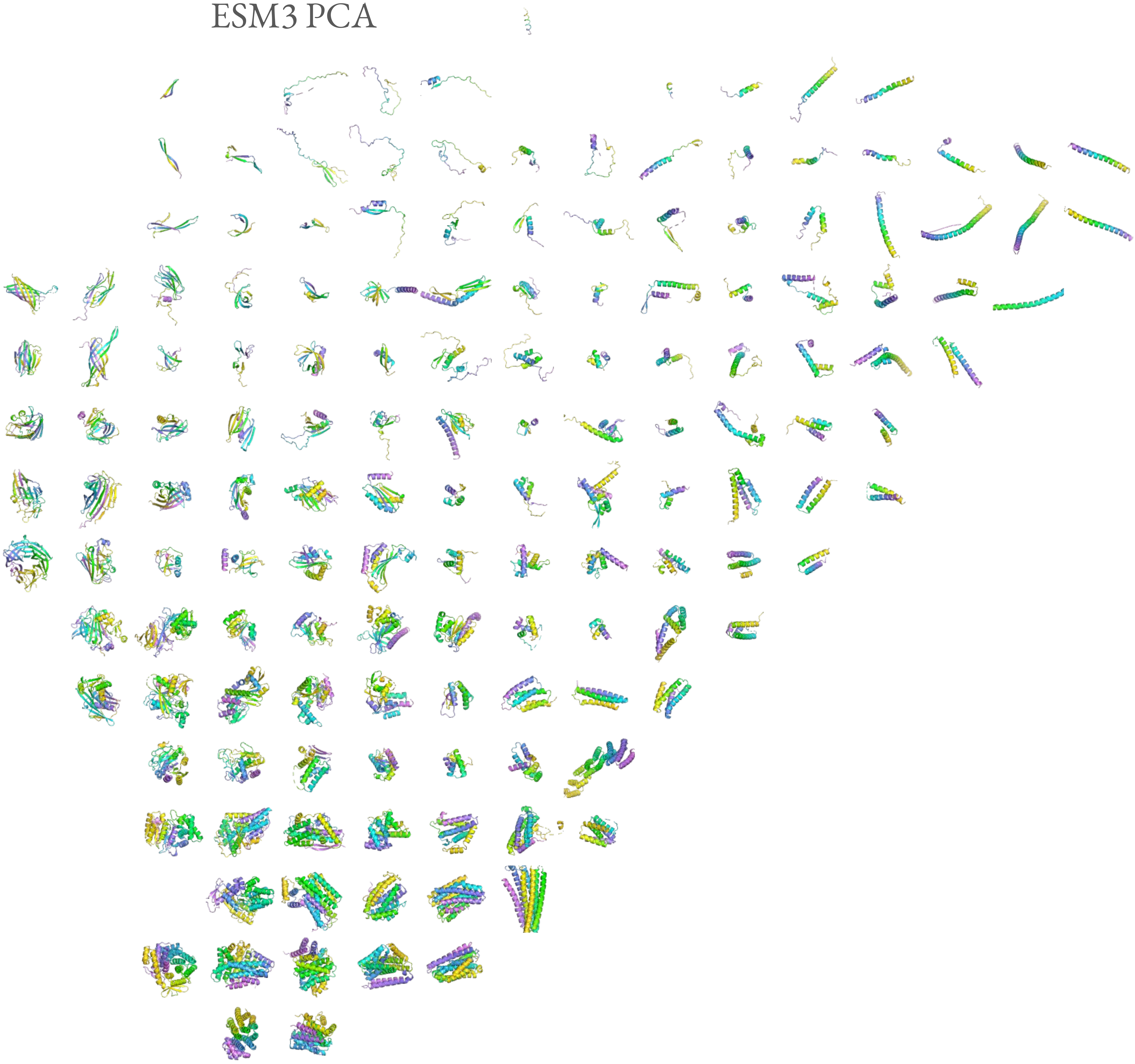
